# The accomplices: Heparan sulfates and N-glycans foster SARS-CoV-2 spike:ACE2 receptor binding and virus priming

**DOI:** 10.1101/2024.02.05.578888

**Authors:** Giulia Paiardi, Matheus Ferraz, Marco Rusnati, Rebecca C. Wade

**Author notes:** Correspondence: Giulia Paiardi, Rebecca C. Wade. **Email:**;. **Author Contributions:** G. P., M. R., and R. C. W. conceptualization; G. P. and M.F. formal analysis; G. P., M.F. and R. C. W. funding acquisition; G. P. and M. F. investigation; G. P. methodology; R. C. W. resources; R. C. W. supervision; G. P. validation; G. P. visualization; G. P. writing—original draft; G.P. and R. C. W. writing—review and editing. **Competing Interest Statement:** The authors declare no competing interests.

## Abstract

Although it is well established that the SARS-CoV-2 spike glycoprotein binds to the host cell ACE2 receptor to initiate infection, far less is known about the tissue tropism and host cell susceptibility to the virus. Differential expression across different cell types of heparan sulfate (HS) proteoglycans, with variably sulfated glycosaminoglycans (GAGs), and their synergistic interactions with host and viral N-glycans may contribute to tissue tropism and host cell susceptibility. Nevertheless, their contribution remains unclear since HS and N-glycans evade experimental characterization. We, therefore, carried out microsecond-long all-atom molecular dynamics simulations, followed by random acceleration molecular dynamics simulations, of the fully glycosylated spike:ACE2 complex with and without highly sulfated GAG chains bound. By considering the model GAGs as surrogates for the highly sulfated HS expressed in lung cells, we identified key novel cell entry mechanisms of spike SARS-CoV-2. We find that HS promotes structural and energetic stabilization of the active conformation of the spike receptor binding domain (RBD) and reorientation of ACE2 toward the N-terminal domain in the same spike subunit as the RBD. Spike and ACE2 N-glycans exert synergistic effects, promoting better packing, strengthening the protein:protein interaction, and prolonging the residence time of the complex. ACE2 and HS binding trigger rearrangement of the S2’ functional protease cleavage site through allosteric interdomain communication. These results thus show that HS has a multifaceted role in facilitating SARS-CoV-2 infection and they provide a mechanistic basis for the development of novel GAG derivatives with anti-SARS-CoV-2 potential.

**Significance Statement:** A key to blocking SARS-CoV-2 infection is to understand why it infects some cell types more than others. Heparan sulfate (HS) proteoglycans are differentially expressed on the surface of host cells and, with ACE2 receptors, provide an entry route for SARS-CoV-2. Here, we used computer simulations to investigate how highly sulfated glycosaminoglycans, a model for HS expressed in lungs, impact the interaction between virus spike and host ACE2. The simulations indicate that HS, together with host and spike N-glycans, stabilizes the spike:ACE2 complex and triggers structural changes, including host protease cleavage, contributing to the SARS-CoV-2 infection mechanism. This study lays the basis for a better understanding of the cell-specificity of SARS-CoV-2 infection and for developing strategies for inhibiting SARS-CoV-2 infection.

## Introduction

Despite the clinical success of vaccines, the continued emergence of SARS-CoV-2 variants highlights the need for a deeper understanding of the infection mechanisms and host tropism for developing targeted therapeutics for COVID-19 patients. The SARS-CoV-2 viral surface spike glycoprotein (spike) is a type I membrane glycoprotein that exists as a homotrimer on the virion surface and mediates the initial steps of infection. Each subunit (S_A,_ S_B,_ S_C_) of the prefusion spike is composed of two regions, S1 and S2. S1, containing the N-terminal domain (NTD) and the receptor binding domain (RBD), wraps around S2, which forms a central helical bundle with two heptad repeats and contains the fusion peptide (FP) ^1-4^. 66 N-glycans are covalently attached to the protein surface of the trimeric spike ectodomain ^5-8^. Beyond generally shielding the protein surface, these N-glycans have been found to control spike activation, modulate the protein:protein interactions, and sterically mask functional epitopes of the viral protein ^9-16^. The prefusion spike has been shown to adopt inactive closed and active open conformations ^1^. In the open, active conformation, the RBD is exposed so that its receptor binding motif (RBm) can interact with the human angiotensin-converting enzyme 2 (ACE2) receptor ^17-19^. The interaction with ACE2 triggers the priming of spike, which entails the cleavage of the S2’ functional site by the human transmembrane serine protease 2 (TMPRSS2) ^20, 21^, the subsequent shedding of S1, and the conformational rearrangement of S2, propelling the FP into the target membrane and initiating membrane fusion ^22, 23^.

Heparan sulfate (HS) proteoglycans (HSPGs) are differentially expressed on the surface of human cells. They are composed of a core protein with covalently attached long, variably sulfated glycosaminoglycans (GAGs). HSPGs are co-receptors for SARS-CoV-2 infection and bind spike via its basic domains ^24-27^. The variability of the sulfation patterns in HS across different individuals, tissues, and cell types may contribute to virus tropism, although this variability hinders their structural characterization and full comprehension of their mechanistic roles ^24, 28-30^. Heparin, a long, linear, highly sulfated GAG, provides a surrogate for HS, such as the highly sulfated lung HS ^24^, in computational and experimental studies due to their structural similarity ^31^. Computational and experimental studies have demonstrated that heparin can act as a potent antiviral agent by competing with HSPGs for binding to spike ^24-27, 32-35^ and, that spike forms a ternary complex by simultaneously binding ACE2 and heparin or HS ^24, 36-38^.

A comprehensive understanding of the dynamics governing the process of SARS-CoV-2 priming upon ACE2 engagement, as well as the roles of N-glycans and HS in viral infection and tissue tropism, has so far evaded experimental characterization, making molecular dynamics (MD) simulations the method of choice for investigating these mechanisms. Therefore, we performed ten 1-µs all-atom conventional MD simulations followed by a total of over 300 ns of random acceleration molecular dynamics (RAMD) to investigate how the interaction between SARS-CoV-2 (Wuhan variant) prefusion spike in the active, open conformation and human ACE2-RBD (hereafter referred to as ACE2) is affected by the binding of highly sulfated GAG chains and how N-glycans synergistically interact with these GAG chains. We also investigate the structural dynamics and allosteric force transmission between ACE2 and spike upon binding of GAG chains. In this study, we thus aim to unveil the mechanisms by which highly sulfated HS, as expressed in cells susceptible to SARS-CoV-2, facilitates viral infection. Therefore, for interpreting these simulations, we consider the GAG chains to provide a model for the highly sulfated HS expressed in lung cells ^24^ and, by coupling these results with extensive available experimental data, lay the basis for a mechanistic understanding of SARS-CoV-2 tropism and host cell susceptibility and the development of novel GAG derivatives with anti-SARS-CoV-2 activity.

## Results and discussion

### Highly sulfated GAGs stabilize the open-spike:ACE2 RBD complex by direct and indirect interactions

We performed molecular modelling and replica all-atom conventional MD simulations to generate representative ensembles of equilibrium complexes of one ACE2 bound to a fully glycosylated spike homotrimer with and without three highly sulfated GAG chains bound (Fig.1). Each simulated system consisted of over 870000 atoms (see Material and Methods for details).

**Figure 1.**
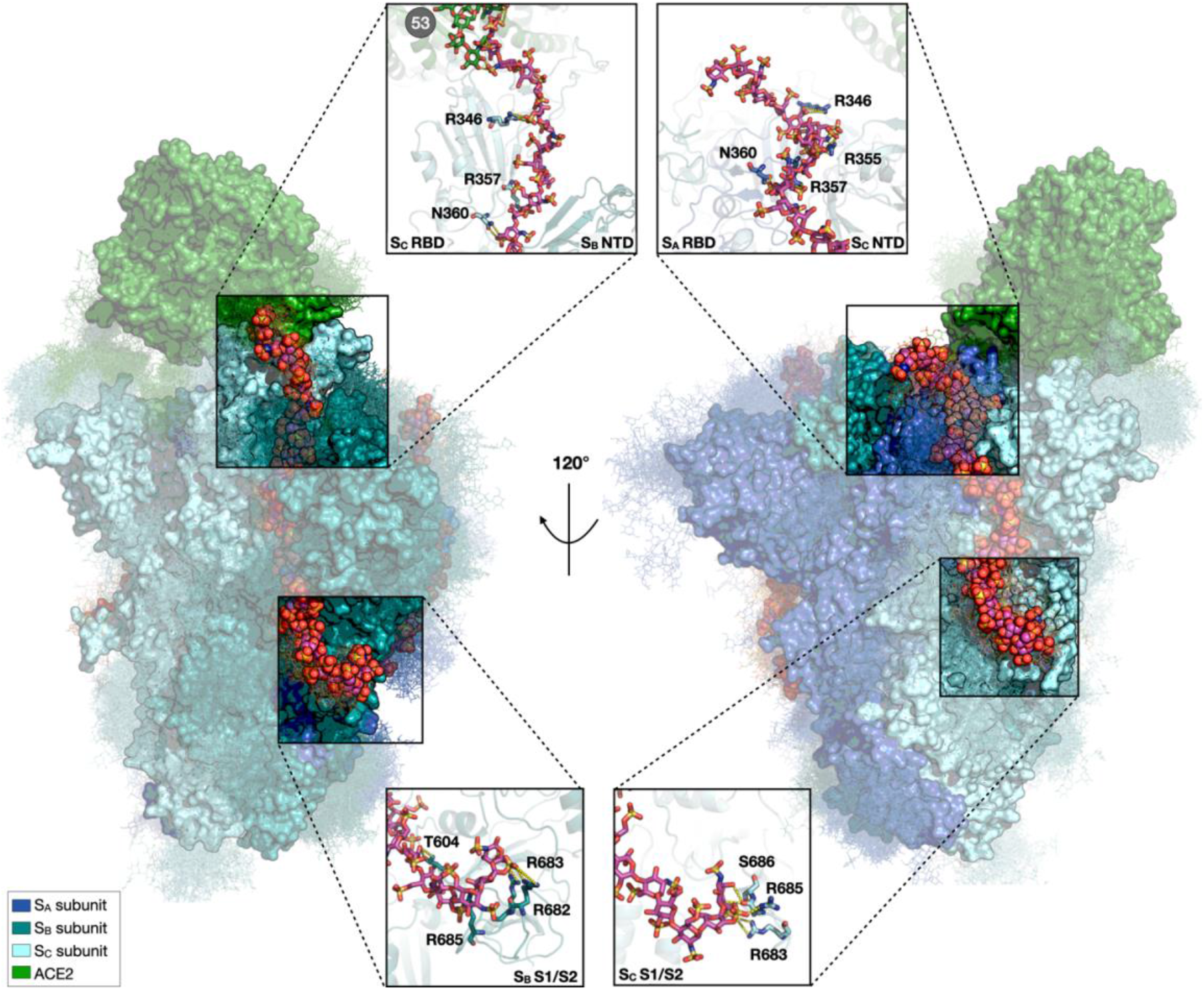
Structural model of the complex of the open, active spike homotrimer, the ACE2 RBD and three GAG chains, showing stabilizing interactions of two GAG chains and N-glycans with spike and ACE2. Two side views of a representative structure obtained from the last snapshot of one of the MD simulation replicas are shown. S_A,_ S_B_, and S_C_ subunits and ACE2 are shown as molecular surfaces in blue, teal, cyan, and green, respectively. N-glycans covalently attached to the spike and ACE2 are shown in line representation, colored according to the subunit to which they are attached. 40 frames of the N-glycan structures collected at intervals of 25 ns from the simulation are shown. The 31mer GAG chains are depicted as spheres colored by element with magenta carbons. On the left, GAG-2 spans from the S_C_ up-RBD to the S_B_ S1/S2 multifunctional domain. On the right, GAG -3 follows a similar path, simultaneously binding the S_A_ down-RBD and the S_C_ NTD and S1/S2.

The open spike:ACE2 complex, both with and without the GAG chains, remained structurally stable during all the conventional MD simulations with no dissociation noticeable by visual inspection. The computed root mean squared deviation (RMSD) confirms the convergence of the systems, which were reasonably equilibrated after 200ns (Fig. S1A). Overall, the systems with GAG chains equilibrated faster than those without and they were more stable along the trajectories. The RMSD values of the systems show an overall lower deviation for S_C_ in the presence of GAGs, most likely due to the direct interaction of the Sc RBD, which is in the open up-conformation (up-RBD), with ACE2. Accordingly, the root mean squared fluctuation (RMSF) shows a lower fluctuation of the S_C_ NTD and RBD in the presence of GAGs (Fig. S1B-C), suggesting a stabilizing effect on the spike:ACE2 interaction.

The highly sulfated GAG chains maintained stable H-bond interactions mainly with residues in the basic domains of spike (Tab. 1). GAG chains 1 and 3 (hereafter referred to as GAG-1 and GAG-3) interact with the down-RBD of one subunit, the NTD and the S1/S2 site on the adjacent subunit and with N-glycans of both subunits (Fig. 1, Tab. 1, and Fig. S2). GAG chain-2 (hereafter referred to as GAG-2) shares the same binding mode although it interacts with the up-RBD (S_C_ RBD) and, intriguingly, the monosaccharides bound to the up-RBD interact tightly with the ACE2 N53 glycan, suggesting a potential stabilizing role, see next sections (Fig. 1, Tab. 1 and Fig. S2), while none of the GAG chains interact with ACE2 residues, in line with previous SPR data ^24^. The computed average molecular mechanics-generalized Born surface area (MM/GBSA) binding free energy (ΔG_bind_) between spike and ACE2 over the last 100 ns of the simulations is - 105.9±5.8 kcal/mol and -123.2±37.0 kcal/mol in the absence and presence of the polysaccharide chains, respectively (Tab. 2), indicating that GAG binding energetically stabilizes the spike:ACE2 complex. Importantly, the strong electrostatic repulsion between highly sulfated GAGs and ACE2, which are both negatively charged, is countered by favorable van der Waals interactions, primarily with the N-glycans, which result in the greater energetic stability of the glycoprotein complex.

**Table 1.** Residues in functional sites in open spike and N-glycans on spike and ACE2 make H-bonds with the GAG chains. H-bond interactions were computed over the 6 replica MD simulations in the presence of GAGs and are given for each spike functional site (RBD, NTD, S1/S2, N-glycans) of each subunit (S_A_, S_B_ or S_C_) as well as for ACE2 N-glycans. No direct H-bond interactions were observed between the GAG chains and ACE2 amino-acid residues. Residues are shown in the 3D structure in Fig. 1, and H-bond occupancies are given in Fig. S2.

|  | **GAG-1** | **GAG-2** | **GAG-3** |
| --- | --- | --- | --- |
| *Spike RBD* | $S_B$ (**down-RBD**): T345, R346, R357, S359, N360 | $S_C$ (**up-RBD**): R346, R357, N360 | $S_A$ (**down-RBD**): R346, R355, R357, N360 |
| *Spike NTD* | $S_A$: T167, T208 | $S_B$: Q173, L176, T286, D287 | $S_C$: N280, N282, T284 |
| *Spike S1/S2* | $S_A$: R682, R683, R685, S689 | $S_B$: T604, R682, R683, R685 | $S_C$: R683, R685, S686 |
| *Spike N-glycans* | $S_A$: N122, N149, N165, N282 | $S_B$: N109, N149, N296 | $S_C$: N122, N282 |
| | $S_B$: N331 | $S_C$: N331 | $S_A$: N331 |
| *ACE2 N-glycans* | - | ACE2: N53 | - |

**Table 2.** Computed binding free energies indicate that the three GAG chains and N-glycans energetically stabilize open spike:ACE2 complexes. Their effect is exerted primarily through favorable van der Waals packing interactions. Computed MM/GBSA energies (kcal/mol) and their energy components are given for the spike:ACE2, S_C_ RBD:ACE2, and S_C_ RBD+NTD:ACE2 interactions in the absence and presence of the GAG chains. GAG-binding energetically stabilizes the complexes, except for the spike RBD:ACE2 complex, in line with measurements from native mass spectrometry coupled to gas-phase ion chemistry ^44^. The ΔG^el^ contribution to ΔG^tot^ for S_C_ RBD:ACE2 with GAG-2 (GAG chain-2) is more repulsive than when other subdomains are included. This observation, coupled with the computed charges of ACE2 and the GAG-2 chain (−27e and -58e, respectively), supports the hypothesis that the decreased binding affinity of S_C_ RBD:ACE2+GAG-2 is due to electrostatic repulsion. – or + denotes that glycans are excluded or included in the computation. The averages and standard deviations of the energy values are computed from the replica simulations of the spike:ACE2 (4 trajectories) and spike:ACE2:GAGs (6 trajectories) systems. Note that while it is well known that absolute values computed with MMGBSA can be overestimated with respect to the experimental observations, the relative values and components are expected to be meaningful ^69, 70^.

|  |  | *MM/GBSA energies (kcal/mol)* |  |  |  |
| --- | --- | --- | --- | --- | --- |
| *N-glycans* | *System* | $\Delta G^{total}$ | $\Delta G^{el}$ | $\Delta G^{vdW}$ | $\Delta G^{solv}$ |
| + | *Spike:ACE2* | -105.88 ± 5.79 | -2777.50 ± 109.17 | -305.19 ± 37.06 | 2976.81 ± 137.04 |
|  | *Spike:ACE2+GAGs* | -123.20 ± 37.03 | 17725.09 ± 693.54 | -352.81 ± 73.55 | -17541.18 ± 733.45 |
| + | *S<sub>C</sub> RBD:ACE2* | -69.96 ± 3.12 | -881.69 ± 37.71 | -171.41 ± 7.74 | 971.64 ± 47.65 |
|  | *S<sub>C</sub> RBD:ACE2+GAG-2* | -49.18 ± 13.13 | 6302.11 ± 358.87 | -147.61 ± 15.12 | -6205.29 ± 404.03 |
| + | *S<sub>C</sub> RBD+NTD:ACE2* | -72.17 ± 12.94 | -1457.42 ± 111.47 | -236.48 ± 46.85 | 1621.74 ± 138.99 |
|  | *S<sub>C</sub> RBD+NTD:ACE2+GAG-2* | -86.04 ± 35.94 | 5649.30 ± 435.47 | -260.60 ± 57.99 | -5477.17 ± 460.36 |
| - | *S<sub>C</sub> RBD+NTD:ACE2* | -33.21 ± 12.94 | -1232.56 ± 80.39 | -116.08 ± 21.45 | 1315.43 ± 99.09 |
|  | *S<sub>C</sub> RBD+NTD:ACE2+GAG-2* | -43.76 ± 16.08 | 5564.62 ± 431.00 | -140.07 ± 24.37 | -5471.27 ± 443.07 |

Overall, the atomic-detail model of the open spike:ACE2:GAGs complex is consistent with surface plasmon resonance and mass photometry data, supporting the ability of the open spike to simultaneously bind to ACE2 and the highly sulfated GAG chains of heparin or HS ^24, 36-38^. Furthermore, these simulations underscore the ability of a single spike bound to ACE2 to accommodate up to 3 GAG chains, in agreement with stoichiometry measurements ^36^. The data support the hypothesis that at the cell surface, highly sulfated HS orchestrates the initial step of cellular attachment by latching onto spike and bringing the virus close to the epithelial cell membrane while exposing the spike RBm and thus promoting encounter with ACE2 and the formation of the ternary spike:ACE2:HS complex. In contrast, the poorly sulfated GAG chains may fail to promote this mechanism ^24, 28^ due to weaker electrostatic forces driving the spike:HS interactions, supporting the hypothesis that HS may be a key driver of SARS-CoV-2 tropism.

### Highly sulfated GAGs enhance the open spike RBD:ACE2 interactions and cross-correlated motion

In the context of SARS-CoV-2 infection, it is necessary to consider all the possible strategies exploited by the virus to optimize the interaction with the host cell receptor. As the simulations show that GAGs stabilize the protein:protein complex, we investigated the effect of their binding at the S_C_ RBm:ACE2 interface. At the interface between S_C_ RBm and ACE2 in the absence of GAGs, residue-residue interactions observed in crystal structures are well-maintained during the trajectories, as shown by H-bond analysis and computed contact occupancies (Fig. S3-4) (Tab. S1). GAG binding results in the loss of the polar interaction between S_C_ N501 and ACE2 Y41 (Fig. 2A), which is not compensated by the formation of interactions with the polysaccharide (Fig. S2). However, their binding leads to the formation of new polar contacts between ACE2 and the spike S_C_ RBm (D355-G502; K353-Y495; H34-S497; S19-G476, respectively) (Fig. 2B, Figs. S3-4, Tab. S1), thereby strengthening the protein:protein interaction.

**Figure 2.**
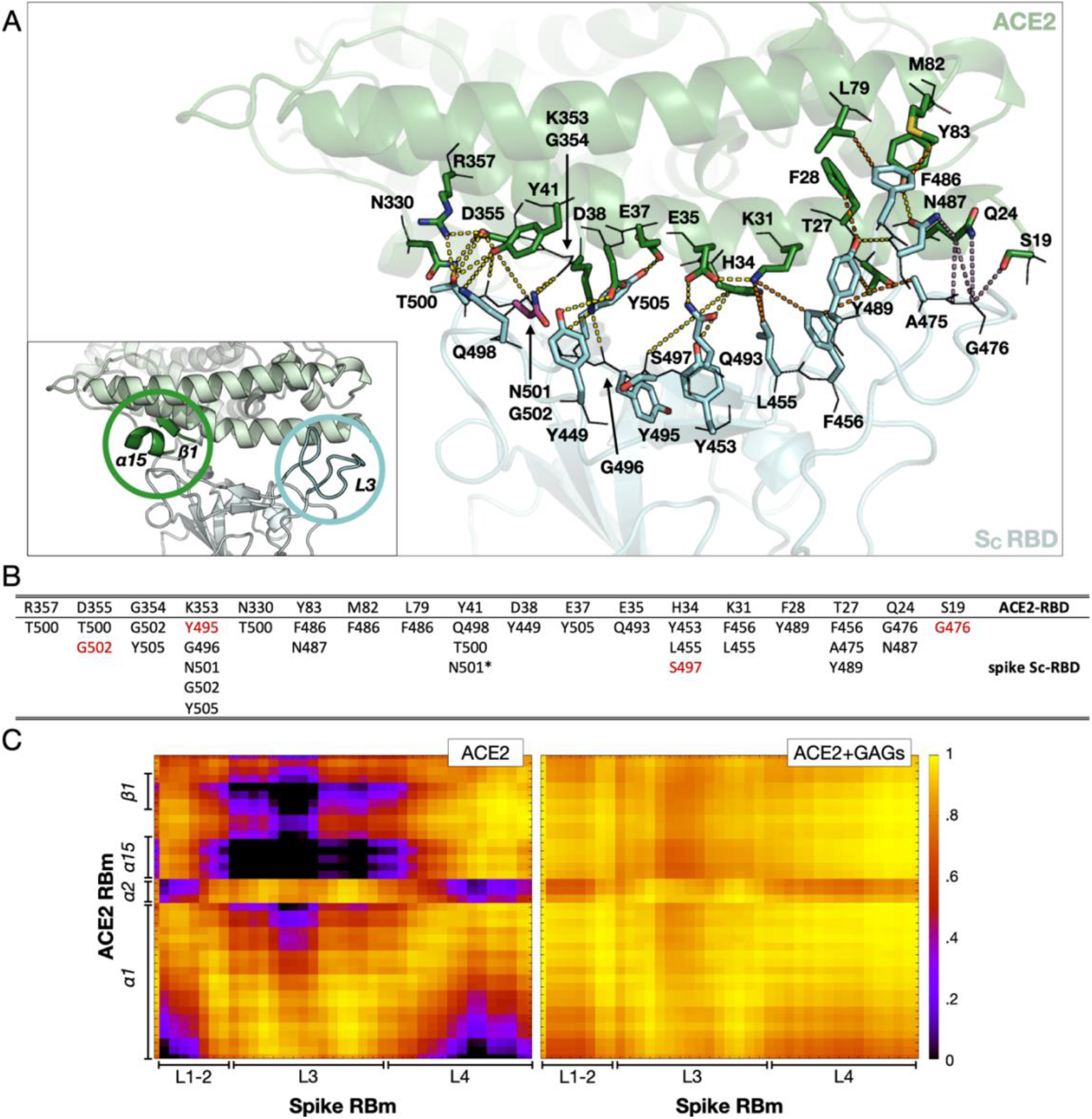
Binding of GAGs alters the interactions at the interface between the spike S_C_ subunit (up-RBD) and ACE2. **(A)** View of the S_C_ RBD – ACE2 interface extracted from the last snapshot of the replica 1 trajectory simulated in the presence of GAGs. The S_C_ RBD and ACE2 are shown as cartoons colored cyan and green, respectively. GAG-2 is omitted for clarity. Residues involved in the interactions are labeled and depicted as lines (main chain) and sticks (side chain) colored by element with oxygen, nitrogen, and sulfur in red, blue, and yellow, respectively. N501 is shown as magenta stick colored by element. Dashes connect interacting residues are colored yellow, orange, and violet to differentiate between polar, hydrophobic, and other types of contact, respectively. Contact matrices for each replica are given in Fig. S4. The inset shows the location in the 3D structure of two regions of spike S_C_ RBD and ACE2 that are dynamically coupled in the presence of GAGs (see C). (B) Interactions between ACE2 (upper line) and S_C_ RBD. Residues colored red gained the respective interactions during the simulations (both with and without GAGs) compared to the interactions reported in the literature ^3, 68^; see Tab. S1. The asterisk for the Y41-N501 interaction indicates that it is lost only in simulations with GAG chains. (C) Dynamic crosscorrelation matrices of residues of S_C_ RBm (x-axis) and ACE2-RBm (y-axis) for the spike:ACE2 (left) and spike:ACE2:GAGs (right) systems. Values range from 0 (black, uncorrelated motion) to +1 (yellow, correlated motion). The plots are shown for the replica 1 trajectory; plots for all replica trajectories are available in Fig. S5.

To investigate the dynamically coupled motion of the complex and how it is affected by the GAG chains, Dynamic Cross-Correlation Matrices were computed for the S_C_ RBm and ACE2 (Fig.2C, Fig.S5). GAG binding increases the correlated movement of the S_C_ RBm L3-loop (residues 472-490) with the ACE2 α15-helix (residues 324-330) and β1-strand (residues 354-357) (Fig. 2A-inset, 2C and Fig. S6). In agreement, the RMSF of the S_C_ RBD and L3-loop, which is known to be highly flexible ^39, 40^, is lower when GAGs are bound, confirming their stabilizing effect (Fig. S7). Notably, Verkhivker and co-workers ^41^ identified a negative cross-correlated movement of these residues in the newer omicron spike variants upon ACE2 binding in the absence of GAGs, while Kim and colleagues ^36^ showed that omicron spike interactions with HS and ACE2 are significantly enhanced compared to the Wuhan variant. Consistent with these and our findings, newer omicron spike variants could require HS to stabilize the protein:protein interface for subsequent steps in the infection process.

In summary, the observed energetics and dynamics of the spike RBD and ACE2 indicate that at the cell surface, the highly sulfated HS of the lung may have a stabilizing effect on the open spike:ACE2 complex while the newer omicron spike variant may exploit the same mechanism with higher affinity.

### Highly sulfated GAGs energetically stabilize the reorientation of ACE2 towards the spike-NTD

To investigate the ability of GAGs to promote rearrangements beyond the S_C_ RBm:ACE2 interface, the RMSD and RMSF of ACE2 (Fig. S1) and orientational vectors between ACE2 and spike were monitored along the trajectories (Fig. 3). The structure of ACE2 is well preserved in simulations with and without GAGs (Fig. S1). However, the distance ***d*** between the center of mass (COM) of ACE2 and the COM of S_C_ NTD is reduced upon GAG binding (Fig. 3A,B). The **α** and **β** angles, defining the positions of ACE2 and the S_C_ NTD with respect to the long axis of the spike homotrimer (Fig. 3A), show a narrower distribution upon GAG binding, suggesting that ACE2 has more freedom to reorient on spike in the absence of the polysaccharide chains and is constrained in a well-defined location upon GAG binding (Fig.3C,D and S8). Overall, the data point to a putative involvement of the S_C_ NTD in the S_C_ RBD:ACE2 interaction, in agreement with a recent report highlighting that mutations in the hypervariable NTD regions increase protein-mediated fusion and cell entry ^42^.

**Figure 3.**
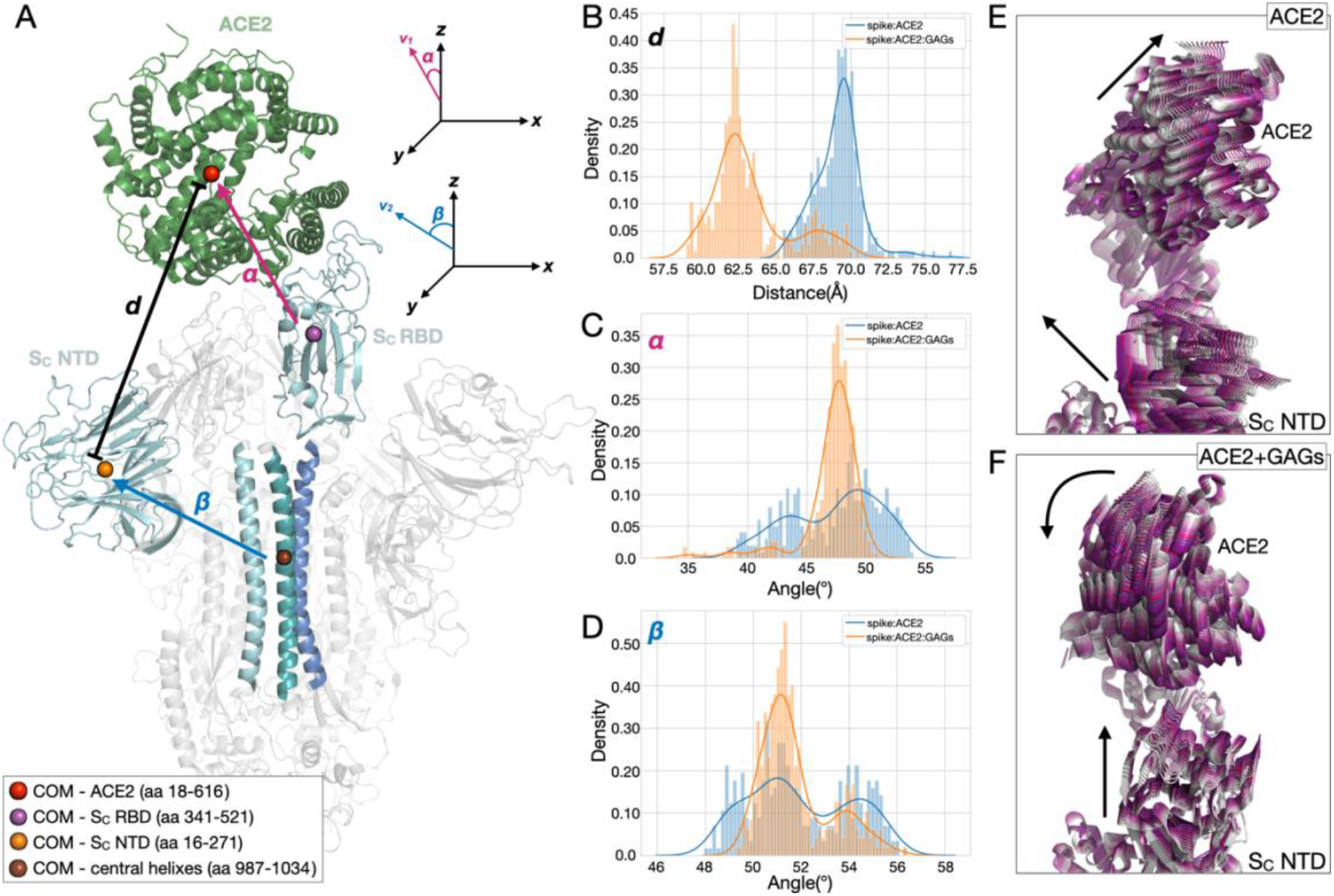
ACE2 and the S_C_ NTD approach each other in the presence of GAGs. (A) Side view of a representative structure of the spike head ectodomain in the open conformation (S_C_ with up-RBD) in complex with ACE2. Spike and ACE2 are shown as cartoons colored grey and green, respectively. The S_C_ RBD, NTD, and central helix are colored cyan. The S_A_ and S_B_ central helices are colored marine and teal, respectively. The spheres represent the center of mass (COM) calculated for each domain and are colored red, violet, orange and brown for ACE2 (residues 18-616), S_C_ RBD (residues 341-521), S_C_ NTD (residues 16-271), and the three spike central helices (residues 987-1034), respectively. To define the approach of ACE2 and the spike, the complex was oriented perpendicular to the xy plane. The distance **d** and vectors **v1** and **v2** were defined as follows: **d** (black) is the distance between the COM of ACE2 and the COM of S_C_ NTD; **v1** (magenta) points from the COM of the S_C_ RBD to the COM of ACE2; **v2** (blue) points from the COM of the three central helices to the COM of the S_C_ NTD. The relative position of ACE2 is defined by (B) the distance **d**, and the angles α (C) and β (D) between the z-axis and **v1** and **v2**, respectively. The distributions (B-D) were calculated from the MD trajectories for all replicas and are colored according to system: spike:ACE2 (blue) and spike:ACE2-GAGs (orange). The motion of ACE2 and the S_C_ RBD in the absence (E) and presence (F) of GAGs is shown by the superimposition of ten conformations extracted at equal time intervals along a representative trajectory (from grey to purple) and projected onto the first essential dynamics eigenvector. ACE2 and S_C_ RBD are shown in cartoon representation, with GAGs omitted for ease of visualization. The corresponding plots for all the replica trajectories are similar and are shown in Fig. S8.

To further investigate the role of the S_C_ NTD, essential dynamics (ED) analysis of the S_C_ subunit and ACE2 in the complexes with and without GAGs was performed. Projection of S_C_:ACE2 dynamics onto the first eigenvector (the vector with the largest variance in the trajectory) shows that, in the absence of GAGs, the motion of ACE2 and S_C_ NTD is randomly oriented (Fig. 3E and S9), in agreement with cryo-EM data ^43^. In contrast, upon GAG binding, S_C_:ACE2 dynamics along the first eigenvector indicate the reorientation of ACE2 and the S_C_ NTD towards each other, in agreement with the reduced distance between their COMs (Fig. 3B, F and S9), indicating that the spike NTD and GAGs may together contribute to the increased affinity of SARS-CoV-2 to ACE2.

To quantify the contribution of the S_C_ NTD to ACE2 binding, the binding free energy of the system was decomposed by calculating the MM/GBSA interaction energy of ACE2 with the S_C_ RBD versus the S_C_ RBD+NTD both in the absence and presence of GAGs. The average ΔG_bind_ between S_C_ RBD and ACE2 is -70.0 ± 3.1 kcal/mol and -49.2 ± 13.1 kcal/mol in the absence and presence of GAG-2, respectively (Tab.2). GAG-2 binding thus has an unfavorable effect on the spike RBD:ACE2 complex, in line with measurements from native mass spectrometry coupled to gas-phase ion chemistry ^44^, due to electrostatic repulsion between the strongly anionic polysaccharide and the negatively charged ACE2. S_C_ NTD does not significantly affect the ΔG_bind_ in the absence of GAG-2, with an average ΔG_bind_ of -72.2 ± 12.9 kcal/mol. Conversely, the S_C_ NTD significantly contributes to the interaction energy with ACE2 in the presence of GAG-2, resulting in an average ΔG_bind_ of -86.0 ± 35.9 kcal/mol. Notably, this S_C_ RBD+NTD:ACE2 interaction energy is almost 70% of the binding free energy calculated for the simulated full-length spike (Tab.2).

Overall, our data suggest that the binding to the highly sulfated HS of the lung may result in closer interactions between ACE2 and the S_C_ NTD, reducing the orientational motion of ACE2 bound to spike, and driving the reorientation of ACE2 toward the S_C_ NTD, resulting in a more compact spike S1 up-subunit. Notably, site-directed mutagenesis and cell assays demonstrated that the NTD allosterically modulates spike-mediated functions, including TMPRSS2-dependent viral entry and fusogenicity ^42, 45, 46^.

### N-glycans of spike and ACE2 concur with highly sulfated GAGs in strengthening the open spike:ACE2 interactions and extending the residence time of the complex

To assess the ability of N-glycans to stabilize the complex, the MM/GBSA S_C_ RBD+NTD:ACE2 interaction energies were decomposed to compute the individual contributions of the protein and the N-glycans (Tab. 2). Interestingly, both with and without GAGs, only half of the total interaction energy comes from protein:protein interactions, while the remaining half is due to glycans. Notably, the same trend is observed for the van der Waals term of the MM/GBSA interaction energy (Tab. 2), indicating that a key role of the N-glycans in stabilizing the spike:ACE2 complex is to fill intermolecular space and make favorable packing interactions. In addition, the buffering effect between the S_C_ NTD and ACE2 is due to a dense H-bond network (Fig. S10 and Movie S1). In the absence of GAGs, H-bonds are mainly established by S_C_ N17, N165 and N343 glycans simultaneously interacting with ACE2 N546 glycan and by spike S_C_ N343 interacting with ACE2 N322 glycan (Fig. S10). GAG binding promotes the interaction of S_C_ N17 with ACE2 N322 and of S_C_ N165 with ACE2 N90 and N322 glycans and with multiple residues of ACE2 (Fig. 4A and S10). Intriguingly, independently of the presence of GAGs, the ACE2 N90 glycan inserts in the pocket between the back of the S_C_ RBD (up-RBD) and S_A_ RBD (Fig. 4A). In the absence of polysaccharide chains, the ACE2 N90 glycan interacts with residues of S_A_ and S_C_ RBDs and S_C_ N165 glycan (Fig. S10). GAG binding strengthens the interaction between ACE2 N90 glycan and S_A_ RBD residues by increasing the occupancy of the H-bonds and introducing further contacts (S_A_ V445, S_A_ S494). Interactions with S_C_ RBD residues are overall maintained with the occupancy of some interactions reduced in favour of others, including the newly gained H-bond with S_C_ G416 and the increased interaction with S_C_ N165 glycan (Fig.4A and S10). These results provide a mechanistic explanation for previous reports of the importance of the N17, N165 and N343 glycans of spike and the N53, N90 and N322 glycans of ACE2 for their interaction ^12, 47-50^, by showing that spike and ACE2 N-glycans establish stable contacts and fill the intermolecular space, promoting more favorable protein:protein interactions. No experimental data are available concerning interactions of heparin or HS with N-glycans, making the model a potential starting point for further investigations.

**Figure. 4.**
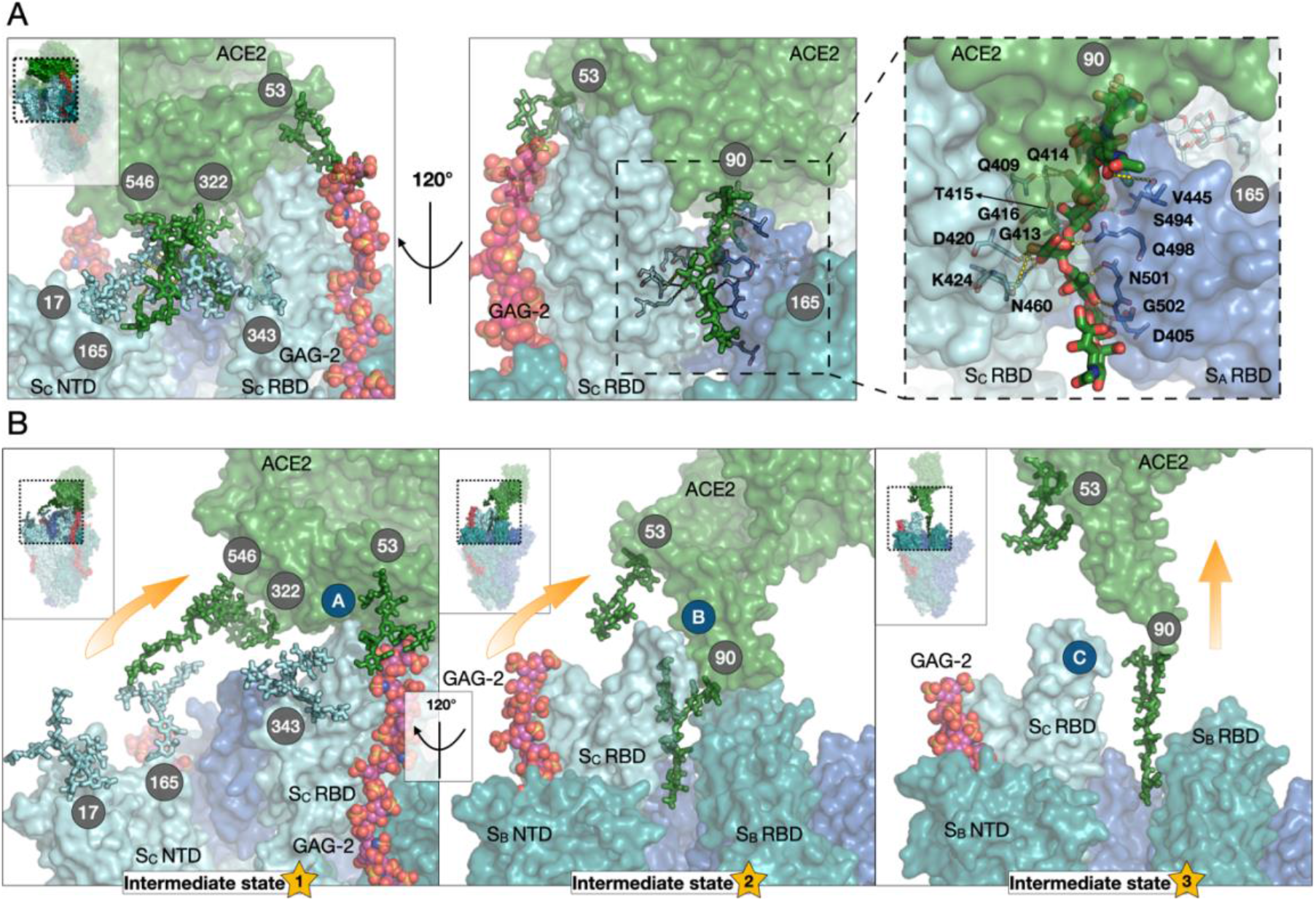
N-glycans and GAGs contribute to stabilization of the open spike:ACE2 complex and hinder its dissociation. (A) Interactions established by the N-glycans in the conventional MD simulations: buffering effect of spike and ACE2 N-glycans at the interface between ACE2 and S_C_ NTD (left); ACE2 N53 glycan:GAG-2 interactions and insertion of ACE2 N90 glycan at the interface between S_C_ RBD and S_A_ RBD (center) with an enlargement showing the interactions established by the N90 glycan with the spike residues (right). (B) The dissociation pathway of ACE2 from spike in the presence of GAGs observed in the RAMD trajectories has three intermediate states. Yellow arrows show the dissociation route of ACE2 as a fan-like rotational movement in the first two intermediate states and a translating movement in the third. Blue circles (labelled A-C) highlight changes in the protein:protein interactions along the dissociation pathway schematically. S_A,_ S_B_, and S_C_ subunits and ACE2 are shown as surfaces in blue, teal, cyan, and green, respectively. Mechanistically key N-glycans covalently attached to spike and ACE2 are labeled with grey circles and shown in stick representation, colored according to the subunit to which they are attached. The 31mer highly sulfated GAG-2 chain bound to the S_C_ up-RBD is depicted as spheres colored by elements with magenta carbons. For further details of the dissociation pathway, see Figs. S11-S13.

To further investigate the mechanistic role of the N-glycans and GAGs, the dissociation of ACE2 from spike was investigated by performing RAMD simulations ^51, 52^. Although the large size of the simulated systems (see Material and Methods for details) meant that fully statistically converged RAMD residence times could not be calculated, the computed RAMD residence times of the spike:ACE2 complex without and with GAG chains bound of 2.2±0.2 and 3.1±1.2 ns, respectively, (Fig. S11) point to the stabilizing effect of the polysaccharide chains that results in increased affinity and residence time of the complex, in line with TR-FRET measurements ^37^. Visual inspection and analysis of the dissociation pathways unveiled the mechanisms by which N-glycans and GAGs prolong the residence time. Dissociation of ACE2 in the absence of polysaccharide chains is in line with results from pulling MD simulations ^53^. Upon GAG binding, ACE2 follows the same dissociation path as without them, visiting multiple intermediate states, although the dissociation is slower (Fig.4B, Fig.S12-13 and Movie S2):

1. In the first intermediate state, interactions between the buffering N-glycans at the interface between the S_C_ NTD (N165, N17, N343) and ACE2 (N322, N546) are disrupted while conserving interactions between the S_C_ RBm and ACE2-RBm (Fig. 4B, Fig. S12-13).
2. In the second intermediate state, the protein:protein dissociation evolves with a fan-like movement of ACE2, starting with the loss of polar interactions between L1-loop and L4-loop of spike and the respective interactors on the α1 helix of ACE2. Interactions between spike RBD L3-loop and the distal residues of the ACE2 N-terminus are still present (Fig. 4B, Fig. S12-13).
3. In the third and last intermediate state, contacts between the spike RBD L3-loop and the distal portion of ACE2 are lost, subsequently inducing the loss of interactions between ACE2 N90 glycan and the S_C_ RBD until ACE2 is completely detached following an upward sliding movement (Fig. 4B, Fig. S12-13).

The second intermediate state is visited for a longer period in the presence of GAGs due to direct interactions between the ACE2 N53 glycan and GAG-2 (Fig. 4B, Fig. S10,12,13). Similarly, in the third intermediate state, the longer dissociation time in the presence of GAGs is due to stronger interactions established between the ACE2 N90 glycan and the spike RBDs, which result in a narrower crevice where the N-glycan is inserted (Fig. S10, S14).

Overall, our data support the hypothesis that N-glycans and highly sulfated HS may synergistically interact, strengthening the spike:ACE2 interaction and packing by extending the residence time of the system.

### Allosteric communication pathways induced by ACE2 binding and modulated by highly sulfated GAGs regulate the rearrangement of the spike-TMPRSS2 site

The pronounced change in the dynamics of the systems upon ACE2 and GAG binding suggests that the binding partners not only affect spike residues at their immediate binding site but also induce long-range effects within spike. To reveal the residues and interactions specifically altered by the binding of ACE2 and GAGs, Force Distribution Analysis (FDA) ^54^ was carried out by calculating the differences in the residue-based pairwise forces in the (i) open spike with and without ACE2 bound; (ii) open spike with ACE2 bound and with and without GAGs bound. Fig. 5*A* shows the connected networks of residue pairs that exhibit changes in pairwise forces that exceed a given threshold. Changes in the force distribution pattern extend through the RBD domain of the S_C_ subunit (bound to ACE2) to the S_A_ RBD, from where the forces further propagate through the central helices of the S_A_ and S_B_ subunits, to the S1/S2 multifunctional domain and furin cleavage site ^55^ of S_B_ and the TMPRSS2 cleavage site of S_A_. Represented as edges of the networks, N-glycans are only marginally involved in the propagation of the forces and are only visible upon application of a high pairwise force threshold (Fig. 5*A*).

**Fig. 5.**
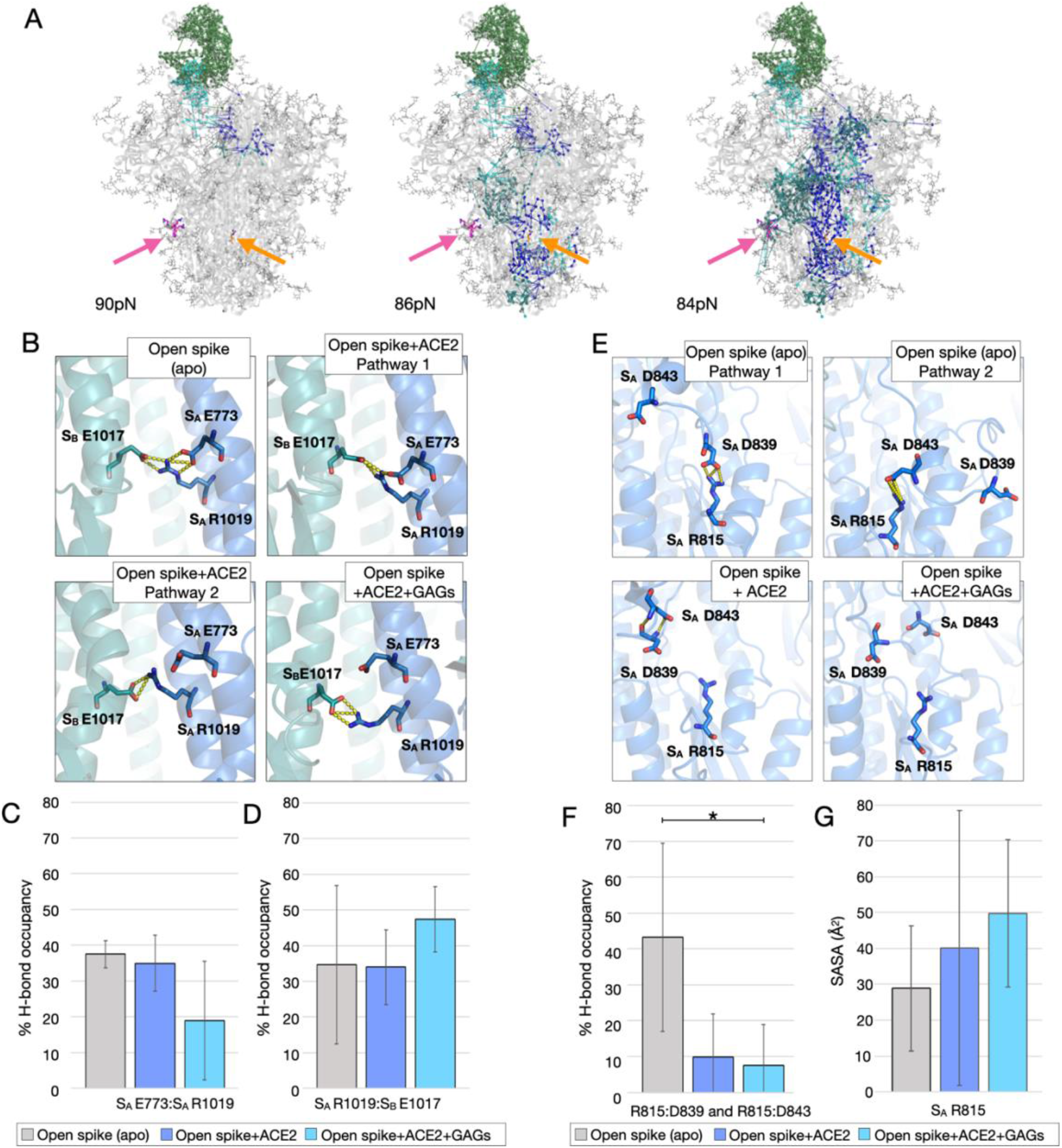
The binding of ACE2 and GAGs allosterically affects the spike protease cleavage sites, promoting virus priming. (A) Residue-based pairwise force differences in the open spike protein when bound or not bound to ACE2 at three force thresholds (lines). The residue-based pairwise force difference upon GAG binding (not shown for clarity) follows the same distribution pattern. The spike and ACE2 glycoproteins are shown as grey cartoons with lines representing the N-glycans. Pairwise force vectors are depicted as lines colored according to the location of the initial point residue (marine, teal, cyan, and green for S_A_, S_B,_ S_C,_ and ACE2, respectively). R815 and the S1/S2 functional site residues are shown in orange and magenta, respectively. (B) Proposed force transduction from S_A_ E773 to S_A_ R1019 and S_A_ E1017 (in the central helices – yellow dashed lines represent H-bonds) and (C-D) the relative trajectory-averaged H-bond occupancy calculated for the last 400ns of the MD simulations. (E) Proposed force transduction from S_A_ R815 (cleavage site of TMPRSS2) to S_A_ D839 and S_A_ D843 (in the fusion peptide - yellow dashed lines represent the H-bonds) and (F) the relative trajectory-averaged H-bond occupancy calculated for the last 400ns of the MD simulations. The solvent-accessible surface area (SASA) of S_A_ R815 along the corresponding trajectories is reported in (G) and shows that binding of ACE2 alone or ACE2 and GAGs increase the exposure of the residue. Data are presented as mean ± SEM. *p< 0.05, two-tail ANOVA.

To further explore the perturbation and potential allosteric effects on the prefusion spike, the changes in the time-averaged punctual stresses on each residue upon ACE2 binding to spike and upon GAG binding to the spike:ACE2 complex were computed. Notable differences in punctual stress were identified upon ACE2 or ACE2 and GAG engagement at residues within the NTD, RBD and central helices of spike even though none of these residues was in direct contact with ACE2 or GAGs (Fig. S15, S16). Visual inspection, H-bond and accessible surface analysis of residues in the spike NTD and RBD did not show significant differences (Fig.S17), suggesting that the stress differences are driven by long-range electrostatic interactions between the negatively charged ACE2 or GAGs and the charged residues of spike in proximity to the binding partners.

Particularly important for the intermolecular allostery upon either ACE2 or GAG binding are the residues located in the distal region of the central helices of each spike subunit (Fig. S15,S16), which have been suggested to have a stabilizing role on the S2 region in the spike prefusion conformation ^56, 57^. Visual inspection and computation of H-bond occupancies revealed a bifurcated salt-bridge network between S_A_ E773, S_A_ R1019 and S_B_ E1017 in spike (Fig. 5B). Upon ACE2 binding, the S_A_ E773 - S_A_ R1019 interaction becomes more transient (Fig. S18), while upon ACE2 and GAG binding, the occupancy of the S_A_ E773 - S_A_ R1019 H-bond decreases in favor of S_A_ R1019 – S_B_ E1017 (Fig. 5C,D), suggesting that the GAGs concur in destabilizing the packing of the central helices and triggering subsequent rearrangements. Interestingly, another important residue for the intermolecular allostery upon ACE2 binding located in proximity to S_A_ E773 is S_A_ R815 (Fig. 5E and Fig. S15), the target of TMPRSS2 cleavage ^21^. Visual inspection of S_A_ R815 and computation of H-bond occupancies revealed that it forms previously unrecognized salt bridges with S_A_ D839 and S_A_ D843 of the adjacent intrinsically disordered FP, that were disrupted upon ACE2 or ACE2 and GAG binding (Fig. 5E,F). These observations are in line with increased exposure of S_A_ R815 upon binding of ACE2 and binding of ACE2 and GAGs (Fig. 5G). This explains how R815 becomes available for priming by the TMPRSS2 protease, or for the binding of specific antibodies directed to this site only upon ACE2 binding ^20, 22, 38, 58^. No punctual stress or H-bonding rearrangement was recorded for R815 of S_B_ and S_C_ upon ACE2 binding (Fig. S15), suggesting that priming of the S1 portion and the subsequent rearrangement of the FP portion is propagated from the up-RBD of the subunit bound to ACE2 via the NTD and central helices to the S2’ of the adjacent spike subunit. The reduced bonding cohesion between the interacting residues in the central helices increases the flexibility of the proximal loop containing R815, triggering the disruption of H-bonding and the subsequent exposition of the S2’ cleavage site.

The S_B_ N657 glycan, which is proximal to S_A_ R815, was the only glycan identified as an additional punctual stress site upon either ACE2 or GAG binding (Fig. S19). Interestingly, site-directed mutagenesis demonstrated that the N657Q/D point mutation increased infectivity for reasons not yet clarified ^13, 59^. Visual inspection and H-bond analysis show that the N657 glycan gains interactions with D843 solely upon ACE2 binding (Fig. S20), and therefore when the residue loses its interactions with S_A_ R815. The simulations suggest that upon ACE2 binding, the N657 glycan interacts with D843, possibly slowing down the exposition and rearrangement of the spike FP, explaining the increased infectivity of the virus after mutation. Furthermore, GAG binding induces loss of interactions between N657 glycan and D843, promoting the sequestering of the N657 glycan via allosteric effects (Fig. S2 and S19), and thereby supporting spike infection by unmasking the spike FP and making it available for subsequent rearrangements.

In conclusion, ACE2 and the highly sulfated HS directly promote viral infection by making spike accessible to cleavage by the human cell-surface proteases, which then leads to the activation of the viral protein. The higher affinity of Delta and Omicron spike for HS may enhance this mechanism. The spike and ACE2 N-glycans do not play a significant role in the propagation of the forces through spike upon ACE2 or GAG binding.

## Conclusions

The infection mechanism of SARS-CoV-2 and the role of HS in priming virus attachment to the host cell receptor ACE2, initiating viral cell entry, has been studied by a range of experimental approaches ^17-19, 24-26, 28, 33, 34, 36, 44^. Surface plasmon resonance and mass photometry showed the formation of ternary spike:ACE2:heparin ^24^ and spike:ACE2:HS complexes ^24, 36, 37^. Nevertheless, the atomic detail structure and dynamics of these complexes have so far evaded experimental characterization, leaving open questions regarding the mechanistic role of the HS co-receptor in SARS-CoV-2 infection and their involvement in the host cell susceptibility and viral tropism.

Here, we adopted a three-pronged strategy to address this gap and investigate the mechanisms exerted by highly sulfated GAGs in promoting the SARS-CoV-2 infection, laying the foundation for a mechanistic understanding of viral tropism and susceptibility to COVID-19. First, by performing microsecond-long conventional MD simulations of the spike:ACE2 complex without or with three highly sulfated GAG chains bound (as a model for highly sulfated lung HS), we demonstrate the ability of GAGs and N-glycans to synergistically concur in dynamically and energetically stabilizing the open spike:ACE2 complex. The polysaccharide chains modulate the conformational dynamics of ACE2 and the spike NTD, promoting better packing of the complex. The simulations reveal that spike N17, N165 and N343 glycans and ACE2 N322 and N546 glycans act as a stabilizing buffer at the spike NTD:ACE2 interface. Consistently, previous reports showed that blocking spike N-glycan biosynthesis significantly reduced spike:ACE2 binding (by 40%) and dramatically reduced (>95%) viral entry into ACE2-expressing cells ^12^. Moreover, experiments show that de-glycosylation of the spike N343 glycan drastically reduces spike RBD binding to ACE2 ^60^ and that ACE2 N322 glycan interacts with spike glycoprotein ^8, 49, 61^.

Next, by performing enhanced sampling RAMD simulations, we demonstrate that GAG binding extends the residence time of the open spike-ACE2 complex, in agreement with ^37^. In addition to the handshake role of the ACE2 N90 glycan described by others ^8, 30, 49, 61, 62^, we find that stronger interactions between the spike and the ACE2 N90 glycan upon GAG binding and the direct interaction between the GAG chain and the ACE2 N53 glycan slow the dissociation of the complex, supporting the hypothesis that HS and the N-glycans may together contribute to strengthening the spike:ACE2 interaction.

Finally, by performing FDA analysis, we identified salt-bridge networks that, upon individual or combined substitution could yield prefusion-stabilized spikes that are more potent for inducing antibodies that neutralize SARS-CoV-2, as previously proposed for other mutations ^56, 57^. Furthermore, we found that an intricate allosteric communication pathway triggered by ACE2 modulates the exposition of R815 and TMPRSS2-dependent viral entry, in agreement with previous reports ^20, 45, 46, 58^. Notably, we show that highly sulfated GAGs concur with ACE2 in modulating this allosteric communication pathway without shielding the previously identified binding site for antibodies targeting the fusion peptide ^58^.

SARS-CoV-2 attachment and infection mechanisms are HS-dependent ^24, 28^. HS expression, composition, and sulfation patterns vary across different cell types and within the same cells according to health or disease conditions. Lung cells and lung cancer patients, who are highly susceptible to COVID-19, have increased HS sulfation levels ^28^. Based on the structural similarity between our model GAG chains and the HS in lungs, we infer that HS and N-glycans exert a synergistic stabilizing effect on the spike:ACE2 complex by multiple mechanisms, explaining the tropism of the SARS-CoV-2 toward highly sulfated cells (see Fig. 6): (1) HS interacts with spike, facilitating the encounter of the virions with ACE2, although HS might be able to shield the ACE2 receptor from the virions ^63^; (2) once spike is in the proximity of the human host cell receptor, the ACE2 N90 and N53 glycans have a pivotal role in spike:HS engagement and interaction, thereby stabilizing the protein:protein complex; (3) subsequently, HS guides the motion of ACE2 toward the NTD of the spike subunit to which ACE2 is bound, while the N-glycans of spike and ACE2 fill the space between the proteins, favoring a more densely packed S1:ACE2 complex and prolonging the residence time of the ACE2-spike complex; (4) ACE2 and HS binding prime the virus infection by promoting the exposition of the fusion peptide residues and the consequent cleavage of spike by TMPRSS2, thus destabilizing the packing and facilitating the subsequent release of the S1:ACE2 complex.

**Figure 6.**
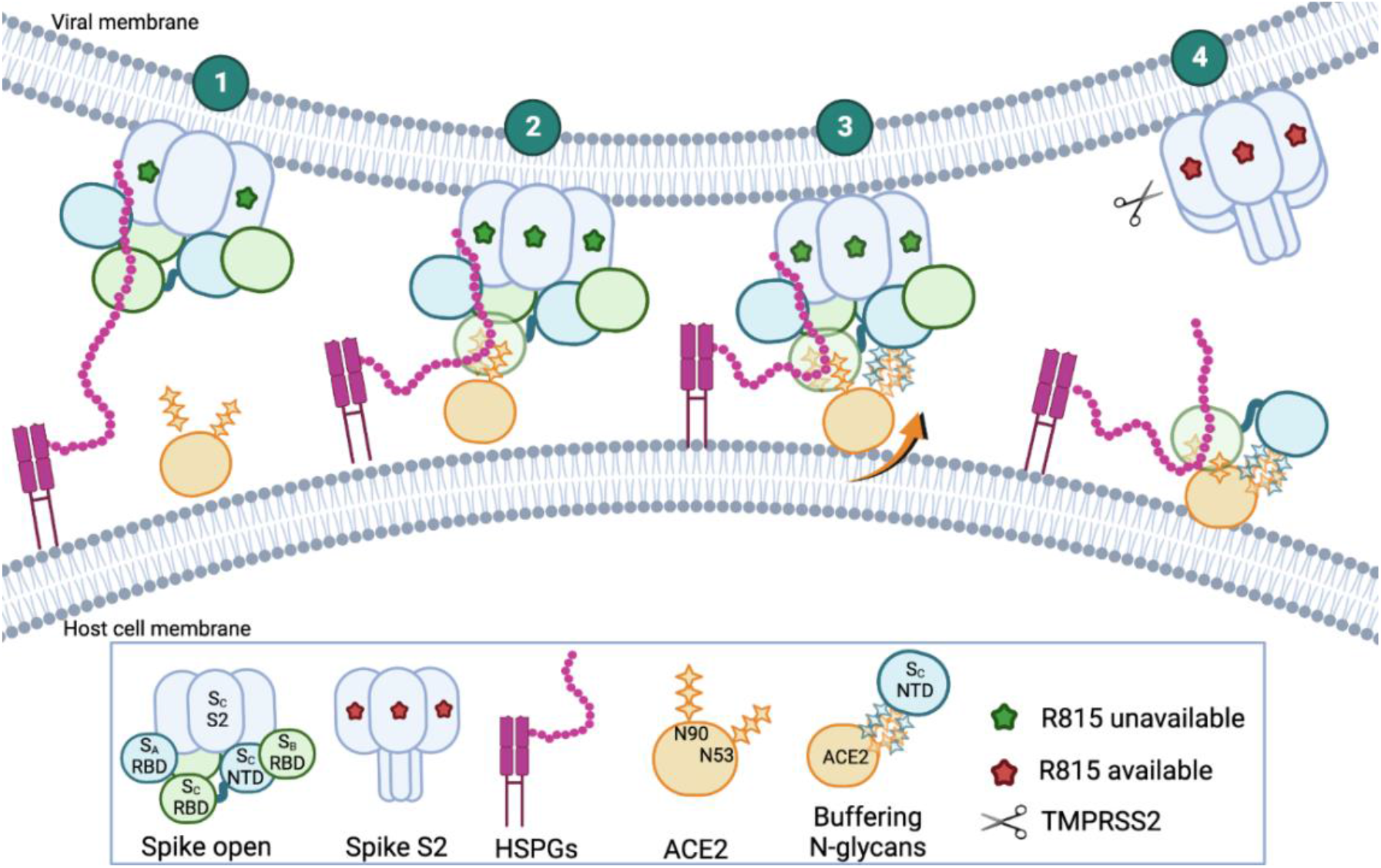
Schematic illustration of the proposed mechanisms by which HSPGs and N-glycans enhance SARS-CoV-2 spike:ACE2 receptor binding and virus priming. The results of the conventional and random acceleration molecular dynamics simulations reported here, along with experimental data reported in the literature, suggest the following mechanisms: HS interacts with SARS-CoV-2 spike glycoprotein, facilitating the encounter with the host cell ACE2 receptor (1). ACE2 N90 glycan drives the handshake with the spike S_C_ up-RBD and subsequently, ACE2 N53 interacts with HS bound to the S_C_ up-RBD, stabilizing the protein:protein complex; HS promotes the formation of residue-residue contacts, structural reordering and the increase in coupled movement in local and distal positions at spike up-RBD:ACE2, resulting in a more energetically favorable complex (2). HS reorients the motion of ACE2 toward the spike NTD of the S_C_ subunit to which the human receptor is bound, while the N-glycans of spike and ACE2 fill the space between the proteins, favoring a more densely packed spike S1:ACE2 complex (3). The motion upon engagement of the human receptor promotes the rearrangement of the allosteric network within spike and releases the constraints imposed on R815, the target of the TMPRSS2 human protease, making it available for cleavage. The cleavage of R815 reduces bonding cohesion and induces conformational metastability of S1-ACE2, promoting the dissociation of the S1-ACE2 component, unlocking the subsequent refolding of S2 (4). Notably, the mechanism proposed is shown for the spike S_C_ subunit only but, based on our data, each spike subunit must undergo this process to expose R815 and be cleaved by TMPRSS2 to promote the rearrangement of S2. RBD, receptor-binding domain; NTD, N-terminal domain; S2, spike fusion domain; HSPGs, heparan sulfate proteoglycans; ACE2, angiotensin-converting enzyme 2; TMPRSS2, transmembrane protease serine subtype 2. Figure created with BioRender (BioRender.com).

Removal or reduction of cell surface HS in physiologically relevant tissues respectively abolishes or significantly reduces viral infection ^24, 28^. Notably, the interaction between spike and HS is driven by electrostatics ^24, 28^. In light of our results, the absence or desulfation of cell surface HS may impede the latching of spike onto the host cell due to the lack of sulfated groups that promote long-range electrostatic recognition of spike, thereby significantly decreasing the likelihood of encountering ACE2 and forming the spike:ACE2:HS ternary complex. On the other hand, reduced sulfation patterns of HS may retain the ability to attach to the spike but either fail in exposing the spike to ACE2, due to too short residence of spike at the cell surface, or successfully promote the encounter of spike with ACE2 but subsequently fail in reorienting ACE2, resulting in a poorly packed complex with short residence time and ineffective cleavage of membrane-anchored TMPRSS2. Since only the most abundant pattern of glycosylation for spike and ACE2, and one sulfation pattern for the highly sulfated GAGs was investigated, further simulations encompassing alternative variable sulfated GAG chains and quantitative assays guided by the simulation results are required to fully characterize the intricate mechanisms exerted by HS in regulating SARS-CoV-2 tropism. Nevertheless, the current computational findings are broadly supported by experimental data reported in the literature, suggesting that the simulated systems represent a reliable model to study the mechanisms involving N-glycans and HS.

The low rate of mutations in the glycosylation sites of SARS-CoV-2 Delta and Omicron spike variants of concern (VoCs)^15^, combined with their high rate of mutation along the GAG binding groove, increasing the positive charge of the spike (Fig.S21 and Ref. ^36^), suggest a potential compensatory mechanism exerted by HS and spike N-glycans to offset the elevated mobility of the VoCs’ RBms. In light of our results, the enhanced affinity for HS of these variants may facilitate the encounter and strengthen the interplay between HS, spike, and ACE2 N-glycans, resulting in more stable complexes with longer residence time, and more efficient cleavage and activation of spike VoCs. Furthermore, the increased charge of the Delta and Omicron spikes potentially increases the electrostatic recognition of highly sulfated HS at long-range, explaining the enhanced affinity for HS and the shifted tropism of SARS-CoV-2 VoCs toward cells that naturally express higher sulfation HS patterns, such as the upper airway tissues. To confirm this hypothesis and elucidate potential susceptibility mechanisms, quantitative assays of the interaction of SARS-CoV-2 VoCs with cells expressing variably sulfated HS and ACE2 mutated at N53 or N90 would be of interest.

In conclusion, from our models and simulations, we have identified modulatory effects of N-glycans and HS on the spike:ACE2 interaction and obtained new insights into the mechanism of SARS-CoV-2 viral tropism and susceptibility. These results support the potential of highly sulfated GAG mimetics, including heparin derivatives, as therapeutics and provide a structural and dynamic framework for developing novel mimetics with enhanced specificity for competing with HS for the binding of spike.

## Materials and Methods

### Modeling of the systems

The two starting structures of the ectodomain of the spike glycoprotein of the Wuhan variant (UniProt A0A679G9E9) in an open, active conformation either with or without three heparin-like GAG chains bound were the initial structures built by Paiardi et al ^35^ (see Tab.S2 for glycomic profile). The three highly sulfated GAG chains were previously modelled using the sliding window method^64^ and span from the RBD of one spike subunit (the S_B,_ S_C_ or S_A_ subunit) to the NTD and S1/S2 functional domains of the adjacent spike subunit (the S_A,_ S_B_ or S_C_ subunit, respectively). Experimental assays showing the direct binding of heparin to spike RBD ^24-27, 34^ and S1/S2 sites ^35^, comparable binding modes obtained with different computational methods ^26, 32, 36^, and an increased binding affinity for GAGs of novel spike variants that exhibit mutations that increase the number of basic residues along the binding path identified (Fig.S21) ^26, 32, 36^ support the modelled positions of the GAG chains. ACE2 was modelled as a monomeric RBD, in line with experimental evidence indicating that dimeric ACE2 can bind up to one RBD of a given spike trimer ^36, 65^; the ACE2 transmembrane anchor and the corresponding membrane were not included in the models. The coordinates for the complex of the ACE2 RBD (residues 19-615) with the spike RBD were retrieved from the RCSB: PDBid 6M0J ^3^ and superimposed on the spike RBD of the starting structures. The spike RBD from the ACE2-spike complex was then removed while retaining the Zn^2+^ ion. The ACE2 was fully glycosylated by adding 6 N-glycans by following the available glycomic profile ^8^ (Tab. S2) and using the Glycam website (http://glycam.org/). Upon visual inspection of the electrostatic potential of ACE2, which has a net negative charge (Fig.S22), and in agreement with SPR data showing the lack of direct binding between ACE2 and heparin ^24^, the modelled GAG chains were not elongated to interact with ACE2. The protonation state was computed at neutral pH using PROPKA ^66^ and PDB2PQR ^67^. None of the spike or ACE2 titratable residues show an unusual protonation state. The two models of the spike:ACE2 and spike:ACE2:GAGs complexes were then solvated and equilibrated for all-atom MD simulation. The last snapshot of each conventional MD replica simulation was used as the starting point for the random acceleration molecular dynamics (RAMD) simulations ^51, 52^. See Tab.S3 for a schematic description of simulations performed and key parameters. For details, see SI.

## Acknowledgments

Stefan Richter, Camilo Aponte and Bernd Doser (Heidelberg Institute for Theoretical Studies) are gratefully acknowledged for technical support. G. P., M.F., and R. C. W. thank the Klaus Tschira Foundation for support. G. P. and R. C. W. thank the German Research Foundation (DFG - Project number: 458623378 to R. C. W.) for financial support. G. P. was supported by the Artificial Intelligence Health Innovation Cluster (post-doc fellowship – 1^st^ cohort) and by the Joachim Herz Stiftung (Add-on fellowship for Interdisciplinary Life Science – 8^th^ cohort). G.P. was supported by the Innogly – Cost Action CA18103 network. M. F. was supported by Erasmus+, the CAPES/DAAD bi-nationally supervised doctoral degree (DAAD, German academic exchange service; CAPES, Coordination for the Improvement of Higher Education Personnel –Research grant number: 57507871). The authors acknowledge support for computing resources from HITS and from the state of Baden-Württemberg through bwHPC and the German Research Foundation (DFG) through grant INST 35/1134-1 and INST 35/1597-1 FUGG. This publication was supported through state funds approved by the State Parliament of Baden-Württemberg for the Innovation Campus Health + Life Science Alliance Heidelberg Mannheim.

## DATA AND CODE AVAILABILITY

Conventional MD simulation replica trajectories of spike:ACE2 complexes in the absence and the presence of GAG chains are available on the BioExcel COVID-19 platform https://bioexcel-cv19.bsc.es/#/ with the identifiers MCV1901746 and MCV190174, respectively. All data for this study are publicly available in the Zenodo repository (doi: 10.5281/zenodo.10214032). All software used is available as described in the Methods. A preprint has been released in bioRXiv at: https://doi.org/10.1101/2024.02.05.578888 under a CC-BY-NC-ND 4.0 International license.

## References

(1) Walls, A. C.; Park, Y.-J.; Tortorici, M. A.; Wall, A.; Mcguire, A. T.; Veesler, D. Structure, Function, and Antigenicity of the SARS-CoV-2 Spike Glycoprotein. Cell 2020, 181 (2), 281-292.e286. DOI: 10.1016/j.cell.2020.02.058 (acccessed 2023-08-14T09:05:30).

(2) Wang, Q.; Zhang, Y.; Wu, L.; Niu, S.; Song, C.; Zhang, Z.; Lu, G.; Qiao, C.; Hu, Y.; Yuen, K.-Y.; et al. Structural and Functional Basis of SARS-CoV-2 Entry by Using Human ACE2. Cell 2020, 181 (4), 894-904.e899. DOI: 10.1016/j.cell.2020.03.045 (acccessed 2023-08-14T09:16:39).

(3) Lan, J.; Ge, J.; Yu, J.; Shan, S.; Zhou, H.; Fan, S.; Zhang, Q.; Shi, X.; Wang, Q.; Zhang, L.; et al. Structure of the SARS-CoV-2 spike receptor-binding domain bound to the ACE2 receptor. Nature 2020, 581 (7807), 215–220. DOI: 10.1038/s41586-020-2180-5 (acccessed 2023-08-14T09:17:56).

(4) Yuan, M.; Wu, N. C.; Zhu, X.; Lee, C. D.; So, R. T. Y.; Lv, H.; Mok, C. K. P.; Wilson, I. A. A highly conserved cryptic epitope in the receptor binding domains of SARS-CoV-2 and SARS-CoV. Science 2020, 368 (6491), 630–633. DOI: 10.1126/science.abb7269.

(5) Watanabe, Y.; Allen, J. D.; Wrapp, D.; McLellan, J. S.; Crispin, M. Site-specific glycan analysis of the SARS-CoV-2 spike. Science 2020, 369 (6501), 330–333. DOI: 10.1126/science.abb9983.

(6) Grant, O. C.; Montgomery, D.; Ito, K.; Woods, R. J. Analysis of the SARS-CoV-2 spike protein glycan shield reveals implications for immune recognition. Scientific Reports 2020, 10 (1). DOI: 10.1038/s41598-020-71748-7 (acccessed 2023-08-14T12:33:24).

(7) Xie, Y.; Butler, M. Quantitative profiling of N-glycosylation of SARS-CoV-2 spike protein variants. Glycobiology 2023, 33 (3), 188–202. DOI: 10.1093/glycob/cwad007.

(8) Zhao, P.; Praissman, J. L.; Grant, O. C.; Cai, Y.; Xiao, T.; Rosenbalm, K. E.; Aoki, K.; Kellman, B. P.; Bridger, R.; Barouch, D. H.; et al. Virus-Receptor Interactions of Glycosylated SARS-CoV-2 Spike and Human ACE2 Receptor. Cell Host & Microbe 2020, 28 (4), 586-601.e586. DOI: 10.1016/j.chom.2020.08.004 (acccessed 2023-08-14T12:39:14).

(9) Casalino, L.; Gaieb, Z.; Goldsmith, J. A.; Hjorth, C. K.; Dommer, A. C.; Harbison, A. M.; Fogarty, C. A.; Barros, E. P.; Taylor, B. C.; Mclellan, J. S.; et al. Beyond Shielding: The Roles of Glycans in the SARS-CoV-2 Spike Protein. ACS Central Science 2020, 6 (10), 1722–1734. DOI: 10.1021/acscentsci.0c01056 (acccessed 2023-08-14T12:40:13).

(10) Sztain, T.; Ahn, S.-H.; Bogetti, A. T.; Casalino, L.; Goldsmith, J. A.; Seitz, E.; Mccool, R. S.; Kearns, F. L.; Acosta-Reyes, F.; Maji, S.; et al. A glycan gate controls opening of the SARS-CoV-2 spike protein. Nature Chemistry 2021, 13 (10), 963–968. DOI: 10.1038/s41557-021-00758-3 (acccessed 2023-08-14T12:40:41).

(11) Harbison, A. M.; Fogarty, C. A.; Phung, T. K.; Satheesan, A.; Schulz, B. L.; Fadda, E. Fine-tuning the spike: role of the nature and topology of the glycan shield in the structure and dynamics of the SARS-CoV-2 S. Chemical Science 2022, 13 (2), 386–395. DOI: 10.1039/d1sc04832e (acccessed 2023-08-14T12:41:23).

(12) Yang, Q.; Hughes, T. A.; Kelkar, A.; Yu, X.; Cheng, K.; Park, S.; Huang, W. C.; Lovell, J. F.; Neelamegham, S. Inhibition of SARS-CoV-2 viral entry upon blocking N- and O-glycan elaboration. Elife 2020, 9. DOI: 10.7554/eLife.61552.

(13) Nangarlia, A.; Hassen, F. F.; Canziani, G.; Bandi, P.; Talukder, C.; Zhang, F.; Krauth, D.; Gary, E. N.; Weiner, D. B.; Bieniasz, P.; et al. Irreversible Inactivation of SARS-CoV-2 by Lectin Engagement with Two Glycan Clusters on the Spike Protein. Biochemistry 2023, 62 (14), 2115–2127. DOI: 10.1021/acs.biochem.3c00109.

(14) Mori, T.; Jung, J.; Kobayashi, C.; Dokainish, H. M.; Re, S.; Sugita, Y. Elucidation of interactions regulating conformational stability and dynamics of SARS-CoV-2 S-protein. Biophysical Journal 2021, 120 (6), 1060–1071. DOI: 10.1016/j.bpj.2021.01.012 (acccessed 2024-02-02T11:19:51).

(15) Newby, M. L.; Fogarty, C. A.; Allen, J. D.; Butler, J.; Fadda, E.; Crispin, M. Variations within the Glycan Shield of SARS-CoV-2 Impact Viral Spike Dynamics. J Mol Biol 2023, 435 (4), 167928. DOI: 10.1016/j.jmb.2022.167928 From NLM.

(16) Turoňová, B.; Sikora, M.; Schürmann, C.; Hagen, W. J. H.; Welsch, S.; Blanc, F. E. C.; von Bülow, S.; Gecht, M.; Bagola, K.; Hörner, C.; et al. In situ structural analysis of SARS-CoV-2 spike reveals flexibility mediated by three hinges. Science 2020, 370 (6513), 203–208. DOI: doi:10.1126/science.abd5223.

(17) Hoffmann, M.; Kleine-Weber, H.; Schroeder, S.; Krüger, N.; Herrler, T.; Erichsen, S.; Schiergens, T. S.; Herrler, G.; Wu, N.-H.; Nitsche, A.; et al. SARS-CoV-2 Cell Entry Depends on ACE2 and TMPRSS2 and Is Blocked by a Clinically Proven Protease Inhibitor. Cell 2020, 181 (2), 271-280.e278. DOI: 10.1016/j.cell.2020.02.052 (acccessed 2023-08-14T12:47:21).

(18) Yang, J.; Petitjean, S. J. L.; Koehler, M.; Zhang, Q.; Dumitru, A. C.; Chen, W.; Derclaye, S.; Vincent, S. P.; Soumillion, P.; Alsteens, D. Molecular interaction and inhibition of SARS-CoV-2 binding to the ACE2 receptor. Nature Communications 2020, 11 (1). DOI: 10.1038/s41467-020-18319-6 (acccessed 2023-08-14T12:47:47).

(19) Shin, Y. H.; Jeong, K.; Lee, J.; Lee, H. J.; Yim, J.; Kim, J.; Kim, S.; Park, S. B. Inhibition of ACE2 ‐Spike Interaction by an ACE2 Binder Suppresses SARS‐CoV‐2 Entry. Angewandte Chemie International Edition 2022, 61 (11). DOI: 10.1002/anie.202115695 (acccessed 2023-08-14T12:48:16).

(20) Yu, S.; Zheng, X.; Zhou, B.; Li, J.; Chen, M.; Deng, R.; Wong, G.; Lavillette, D.; Meng, G. SARS-CoV-2 spike engagement of ACE2 primes S2’ site cleavage and fusion initiation. Proc Natl Acad Sci U S A 2022, 119 (1). DOI: 10.1073/pnas.2111199119.

(21) Vankadari, N.; Ketavarapu, V.; Mitnala, S.; Vishnubotla, R.; Reddy, D. N.; Ghosal, D. Structure of Human TMPRSS2 in Complex with SARS-CoV-2 Spike Glycoprotein and Implications for Potential Therapeutics. The Journal of Physical Chemistry Letters 2022, 13 (23), 5324–5333. DOI: 10.1021/acs.jpclett.2c00967 (acccessed 2023-08-14T12:50:49).

(22) Essalmani, R.; Jain, J.; Susan-Resiga, D.; Andréo, U.; Evagelidis, A.; Derbali, R. M.; Huynh, D. N.; Dallaire, F.; Laporte, M.; Delpal, A.; et al. Distinctive Roles of Furin and TMPRSS2 in SARS-CoV-2 Infectivity. J Virol 2022, 96 (8), e0012822. DOI: 10.1128/jvi.00128-22.

(23) Dodero-Rojas, E.; Onuchic, J. N.; Whitford, P. C. Sterically confined rearrangements of SARS-CoV-2 Spike protein control cell invasion. Elife 2021, 10. DOI: 10.7554/eLife.70362.

(24) Clausen, T. M.; Sandoval, D. R.; Spliid, C. B.; Pihl, J.; Perrett, H. R.; Painter, C. D.; Narayanan, A.; Majowicz, S. A.; Kwong, E. M.; Mcvicar, R. N.; et al. SARS-CoV-2 Infection Depends on Cellular Heparan Sulfate and ACE2. Cell 2020, 183 (4), 1043-1057.e1015. DOI: 10.1016/j.cell.2020.09.033 (acccessed 2023-08-14T12:53:53).

(25) Kim, S. Y.; Jin, W.; Sood, A.; Montgomery, D. W.; Grant, O. C.; Fuster, M. M.; Fu, L.; Dordick, J. S.; Woods, R. J.; Zhang, F.; et al. Characterization of heparin and severe acute respiratory syndrome-related coronavirus 2 (SARS-CoV-2) spike glycoprotein binding interactions. Antiviral Res 2020, 181, 104873. DOI: 10.1016/j.antiviral.2020.104873.

(26) Liu, L.; Chopra, P.; Li, X.; Bouwman, K. M.; Tompkins, S. M.; Wolfert, M. A.; De Vries, R. P.; Boons, G.-J. Heparan Sulfate Proteoglycans as Attachment Factor for SARS-CoV-2. ACS Central Science 2021, 7 (6), 1009–1018. DOI: 10.1021/acscentsci.1c00010 (acccessed 2023-08-14T12:58:49).

(27) Sun, L.; Chopra, P.; Tomris, I.; Van Der Woude, R.; Liu, L.; De Vries, R. P.; Boons, G.-J. Well-Defined Heparin Mimetics Can Inhibit Binding of the Trimeric Spike of SARS-CoV-2 in a Length-Dependent Manner. JACS Au 2023, 3 (4), 1185–1195. DOI: 10.1021/jacsau.3c00042 (acccessed 2023-08-14T12:59:38).

(28) Yue, J.; Jin, W.; Yang, H.; Faulkner, J.; Song, X.; Qiu, H.; Teng, M.; Azadi, P.; Zhang, F.; Linhardt, R. J.; et al. Heparan Sulfate Facilitates Spike Protein-Mediated SARS-CoV-2 Host Cell Invasion and Contributes to Increased Infection of SARS-CoV-2 G614 Mutant and in Lung Cancer. Frontiers in Molecular Biosciences 2021, 8, Original Research. DOI: 10.3389/fmolb.2021.649575.

(29) Zhang, Q.; Chen, C. Z.; Swaroop, M.; Xu, M.; Wang, L.; Lee, J.; Wang, A. Q.; Pradhan, M.; Hagen, N.; Chen, L.; et al. Heparan sulfate assists SARS-CoV-2 in cell entry and can be targeted by approved drugs in vitro. Cell Discov 2020, 6 (1), 80. DOI: 10.1038/s41421-020-00222-5 From NLM.

(30) Kearns, F. L.; Sandoval, D. R.; Casalino, L.; Clausen, T. M.; Rosenfeld, M. A.; Spliid, C. B.; Amaro, R. E.; Esko, J. D. Spike-heparan sulfate interactions in SARS-CoV-2 infection. Current Opinion in Structural Biology 2022, 76, 102439. DOI: 10.1016/j.sbi.2022.102439 (acccessed 2023-08-14T13:00:19).

(31) Hogwood, J.; Mulloy, B.; Lever, R.; Gray, E.; Page, C. P. Pharmacology of Heparin and Related Drugs: An Update. Pharmacological Reviews 2023, 75 (2), 328–379. DOI: 10.1124/pharmrev.122.000684 (acccessed 2023-08-14T13:02:00).

(32) Kim, S. H.; Kearns, F. L.; Rosenfeld, M. A.; Casalino, L.; Papanikolas, M. J.; Simmerling, C.; Amaro, R. E.; Freeman, R. GlycoGrip: Cell Surface-Inspired Universal Sensor for Betacoronaviruses. ACS Central Science 2022, 8 (1), 22–42. DOI: 10.1021/acscentsci.1c01080 (acccessed 2023-08-14T14:49:08).

(33) Tree, J. A.; Turnbull, J. E.; Buttigieg, K. R.; Elmore, M. J.; Coombes, N.; Hogwood, J.; Mycroft‐West, C. J.; Lima, M. A.; Skidmore, M. A.; Karlsson, R.; et al. Unfractionated heparin inhibits live wild type SARS‐CoV‐2 cell infectivity at therapeutically relevant concentrations. British Journal of Pharmacology 2021, 178 (3), 626–635. DOI: 10.1111/bph.15304 (acccessed 2023-08-14T13:03:52).

(34) Mycroft-West, C. J.; Su, D.; Pagani, I.; Rudd, T. R.; Elli, S.; Gandhi, N. S.; Guimond, S. E.; Miller, G. J.; Meneghetti, M. C. Z.; Nader, H. B.; et al. Heparin Inhibits Cellular Invasion by SARS-CoV-2: Structural Dependence of the Interaction of the Spike S1 Receptor-Binding Domain with Heparin. Thrombosis and Haemostasis 2020, 120 (12), 1700–1715. DOI: 10.1055/s-0040-1721319 (acccessed 2023-08-14T13:05:00).

(35) Paiardi, G.; Richter, S.; Oreste, P.; Urbinati, C.; Rusnati, M.; Wade, R. C. The binding of heparin to spike glycoprotein inhibits SARS-CoV-2 infection by three mechanisms. Journal of Biological Chemistry 2022, 298 (2), 101507. DOI: 10.1016/j.jbc.2021.101507 (acccessed 2023-08-14T13:05:43).

(36) Kim, S. H.; Kearns, F. L.; Rosenfeld, M. A.; Votapka, L.; Casalino, L.; Papanikolas, M.; Amaro, R. E.; Freeman, R. SARS-CoV-2 evolved variants optimize binding to cellular glycocalyx. Cell Rep Phys Sci 2023, 4 (4), 101346. DOI: 10.1016/j.xcrp.2023.101346.

(37) Cecon, E.; Burridge, M.; Cao, L.; Carter, L.; Ravichandran, R.; Dam, J.; Jockers, R. SARS-COV-2 spike binding to ACE2 in living cells monitored by TR-FRET. Cell Chemical Biology 2022, 29 (1), 74-83.e74. DOI: 10.1016/j.chembiol.2021.06.008 (acccessed 2023-08-14T13:24:02).

(38) Petitjean, S. J. L.; Eeckhout, S.; Delguste, M.; Zhang, Q.; Durlet, K.; Alsteens, D. Heparin-Induced Allosteric Changes in SARS-CoV-2 Spike Protein Facilitate ACE2 Binding and Viral Entry. Nano Lett 2023, 23 (24), 11678–11684. DOI: 10.1021/acs.nanolett.3c03550 From NLM.

(39) Williams, J. K.; Wang, B.; Sam, A.; Hoop, C. L.; Case, D. A.; Baum, J. Molecular dynamics analysis of a flexible loop at the binding interface of the SARS‐CoV-2 spike protein receptor‐binding domain. Proteins: Structure, Function, and Bioinformatics 2022, 90 (5), 1044–1053. DOI: 10.1002/prot.26208 (acccessed 2023-08-14T13:09:05).

(40) Spinello, A.; Saltalamacchia, A.; Magistrato, A. Is the Rigidity of SARS-CoV-2 Spike Receptor-Binding Motif the Hallmark for Its Enhanced Infectivity? Insights from All-Atom Simulations. The Journal of Physical Chemistry Letters 2020, 11 (12), 4785–4790. DOI: 10.1021/acs.jpclett.0c01148 (acccessed 2023-08-14T13:10:18).

(41) Verkhivker, G.; Alshahrani, M.; Gupta, G.; Xiao, S.; Tao, P. Probing conformational landscapes of binding and allostery in the SARS-CoV-2 omicron variant complexes using microsecond atomistic simulations and perturbation-based profiling approaches: hidden role of omicron mutations as modulators of allosteric signaling and epistatic relationships. Physical Chemistry Chemical Physics 2023, 25 (32), 21245-21266, 10.1039/D3CP02042H. DOI: 10.1039/D3CP02042H.

(42) Qing, E.; Kicmal, T.; Kumar, B.; Hawkins, G. M.; Timm, E.; Perlman, S.; Gallagher, T. Dynamics of SARS-CoV-2 Spike Proteins in Cell Entry: Control Elements in the Amino-Terminal Domains. mBio 2021, 12 (4), e0159021. DOI: 10.1128/mBio.01590-21.

(43) Xu, C.; Wang, Y.; Liu, C.; Zhang, C.; Han, W.; Hong, X.; Hong, Q.; Wang, S.; Zhao, Q.; Yang, Y.; et al. Conformational dynamics of SARS-CoV-2 trimeric spike glycoprotein in complex with receptor ACE2 revealed by cryo-EM. Sci Adv 2021, 7 (1). DOI: 10.1126/sciadv.abe5575.

(44) Yang, Y.; Du, Y.; Kaltashov, I. A. The Utility of Native MS for Understanding the Mechanism of Action of Repurposed Therapeutics in COVID-19: Heparin as a Disruptor of the SARS-CoV-2 Interaction with Its Host Cell Receptor. Analytical Chemistry 2020, 92 (16), 10930–10934. DOI: 10.1021/acs.analchem.0c02449 (acccessed 2023-08-14T13:15:36).

(45) Qing, E.; Li, P.; Cooper, L.; Schulz, S.; Jäck, H. M.; Rong, L.; Perlman, S.; Gallagher, T. Inter-domain communication in SARS-CoV-2 spike proteins controls protease-triggered cell entry. Cell Rep 2022, 39 (5), 110786. DOI: 10.1016/j.celrep.2022.110786 From NLM.

(46) Meng, B.; Datir, R.; Choi, J.; Bradley, J. R.; Smith, K. G. C.; Lee, J. H.; Gupta, R. K.; Baker, S.; Dougan, G.; Hess, C.; et al. SARS-CoV-2 spike N-terminal domain modulates TMPRSS2-dependent viral entry and fusogenicity. Cell Reports 2022, 40 (7), 111220. DOI: 10.1016/j.celrep.2022.111220 (acccessed 2023-08-14T13:16:49).

(47) Huang, H.-C.; Lai, Y.-J.; Liao, C.-C.; Wang, F.-Y.; Huang, K.-B.; Lee, I.-J.; Chou, W.-C.; Wang, S.-H.; Wang, L.-H.; Hsu, J.-M.; et al. Targeting conserved N-glycosylation blocks SARS-CoV-2 variant infection in vitro. eBioMedicine 2021, 74, 103712. DOI: 10.1016/j.ebiom.2021.103712 (acccessed 2023-08-14T13:18:39).

(48) Li, Q.; Wu, J.; Nie, J.; Zhang, L.; Hao, H.; Liu, S.; Zhao, C.; Zhang, Q.; Liu, H.; Nie, L.; et al. The Impact of Mutations in SARS-CoV-2 Spike on Viral Infectivity and Antigenicity. Cell 2020, 182 (5), 1284-1294.e1289. DOI: 10.1016/j.cell.2020.07.012 (acccessed 2023-08-14T13:19:30).

(49) Mehdipour, A. R.; Hummer, G. Dual nature of human ACE2 glycosylation in binding to SARS-CoV-2 spike. Proceedings of the National Academy of Sciences 2021, 118 (19), e2100425118. DOI: 10.1073/pnas.2100425118 (acccessed 2023-08-14T13:20:01).

(50) Hsu, Y. P.; Frank, M.; Mukherjee, D.; Shchurik, V.; Makarov, A.; Mann, B. F. Structural remodeling of SARS-CoV-2 spike protein glycans reveals the regulatory roles in receptor-binding affinity. Glycobiology 2023, 33 (2), 126–137. DOI: 10.1093/glycob/cwac077.

(51) Kokh, D. B.; Amaral, M.; Bomke, J.; Grädler, U.; Musil, D.; Buchstaller, H.-P.; Dreyer, M. K.; Frech, M.; Lowinski, M.; Vallee, F.; et al. Estimation of Drug-Target Residence Times by τ-Random Acceleration Molecular Dynamics Simulations. Journal of Chemical Theory and Computation 2018, 14 (7), 3859–3869. DOI: 10.1021/acs.jctc.8b00230 (acccessed 2023-08-14T13:23:19).

(52) Kokh, D. B.; Doser, B.; Richter, S.; Ormersbach, F.; Cheng, X.; Wade, R. C. A workflow for exploring ligand dissociation from a macromolecule: Efficient random acceleration molecular dynamics simulation and interaction fingerprint analysis of ligand trajectories. The Journal of Chemical Physics 2020, 153 (12), 125102. DOI: 10.1063/5.0019088 (acccessed 2023-08-14T14:01:36).

(53) Zhu, R.; Canena, D.; Sikora, M.; Klausberger, M.; Seferovic, H.; Mehdipour, A. R.; Hain, L.; Laurent, E.; Monteil, V.; Wirnsberger, G.; et al. Force-tuned avidity of spike variant-ACE2 interactions viewed on the single-molecule level. Nat Commun 2022, 13 (1), 7926. DOI: 10.1038/s41467-022-35641-3 From NLM.

(54) Costescu, B. I.; Gräter, F. Time-resolved force distribution analysis. BMC Biophysics 2013, 6 (1), 5. DOI: 10.1186/2046-1682-6-5 (acccessed 2023-08-14T14:10:09).

(55) Cheng, M. H.; Zhang, S.; Porritt, R. A.; Noval Rivas, M.; Paschold, L.; Willscher, E.; Binder, M.; Arditi, M.; Bahar, I. Superantigenic character of an insert unique to SARS-CoV-2 spike supported by skewed TCR repertoire in patients with hyperinflammation. Proc Natl Acad Sci U S A 2020, 117 (41), 25254–25262. DOI: 10.1073/pnas.2010722117 From NLM.

(56) Hsieh, C.-L.; Goldsmith, J. A.; Schaub, J. M.; Divenere, A. M.; Kuo, H.-C.; Javanmardi, K.; Le, K. C.; Wrapp, D.; Lee, A. G.; Liu, Y.; et al. Structure-based design of prefusion-stabilized SARS-CoV-2 spikes. Science 2020, 369 (6510), 1501–1505. DOI: 10.1126/science.abd0826 (acccessed 2024-02-02T14:51:47).

(57) Lu, M.; Chamblee, M.; Zhang, Y.; Ye, C.; Dravid, P.; Park, J.-G.; Mahesh, K.; Trivedi, S.; Murthy, S.; Sharma, H.; et al. SARS-CoV-2 prefusion spike protein stabilized by six rather than two prolines is more potent for inducing antibodies that neutralize viral variants of concern. Proceedings of the National Academy of Sciences 2022, 119 (35). DOI: 10.1073/pnas.2110105119 (acccessed 2024-02-02T14:52:36).

(58) Low, J. S.; Jerak, J.; Tortorici, M. A.; McCallum, M.; Pinto, D.; Cassotta, A.; Foglierini, M.; Mele, F.; Abdelnabi, R.; Weynand, B.; et al. ACE2-binding exposes the SARS-CoV-2 fusion peptide to broadly neutralizing coronavirus antibodies. Science 2022, 377 (6607), 735–742. DOI: 10.1126/science.abq2679.

(59) Yang, Q.; Kelkar, A.; Sriram, A.; Hombu, R.; Hughes, T. A.; Neelamegham, S. Role for N-glycans and calnexin-calreticulin chaperones in SARS-CoV-2 Spike maturation and viral infectivity. Science Advances 2022, 8 (38). DOI: 10.1126/sciadv.abq8678 (acccessed 2023-08-14T13:30:18).

(60) Zheng, L.; Ma, Y.; Chen, M.; Wu, G.; Yan, C.; Zhang, X. E. SARS-CoV-2 spike protein receptor-binding domain N-glycans facilitate viral internalization in respiratory epithelial cells. Biochem Biophys Res Commun 2021, 579, 69–75. DOI: 10.1016/j.bbrc.2021.09.053 From NLM.

(61) Capraz, T.; Kienzl, N. F.; Laurent, E.; Perthold, J. W.; Föderl-Höbenreich, E.; Grünwald-Gruber, C.; Maresch, D.; Monteil, V.; Niederhöfer, J.; Wirnsberger, G.; et al. Structure-guided glyco-engineering of ACE2 for improved potency as soluble SARS-CoV-2 decoy receptor. Elife 2021, 10. DOI: 10.7554/eLife.73641 From NLM.

(62) Barros, E. P.; Casalino, L.; Gaieb, Z.; Dommer, A. C.; Wang, Y.; Fallon, L.; Raguette, L.; Belfon, K.; Simmerling, C.; Amaro, R. E. The flexibility of ACE2 in the context of SARS-CoV-2 infection. Biophys J 2021, 120 (6), 1072–1084. DOI: 10.1016/j.bpj.2020.10.036.

(63) Conca, D. V.; Bano, F.; Wirén, J. v.; Scherrer, L.; Svirelis, J.; Thorsteinsson, K.; Dahlin, A.; Bally, M. Variant-specific interactions at the plasma membrane: Heparan sulfate’s impact on SARS-CoV-2 binding kinetics. bioRxiv 2024, 2024.2001.2010.574981. DOI: 10.1101/2024.01.10.574981.

(64) Bugatti, A.; Paiardi, G.; Urbinati, C.; Chiodelli, P.; Orro, A.; Uggeri, M.; Milanesi, L.; Caruso, A.; Caccuri, F.; D’Ursi, P.; et al. Heparin and heparan sulfate proteoglycans promote HIV-1 p17 matrix protein oligomerization: computational, biochemical and biological implications. Scientific Reports 2019, 9 (1). DOI: 10.1038/s41598-019-52201-w (acccessed 2023-08-14T13:42:02).

(65) Asor, R.; Olerinyova, A.; Burnap, S. A.; Kushwah, M. S.; Soltermann, F.; Powell Rudden, L.; Hensen, M.; Vasiljevic, S.; Brun, J.; Hill, M.; et al. Cooperativity and induced oligomerisation control the interaction of SARS-CoV-2 with its cellular receptor and patient-derived antibodies. bioRxiv 2023, 2023.2009.2014.557399. DOI: 10.1101/2023.09.14.557399.

(66) Li, H.; Robertson, A. D.; Jensen, J. H. Very fast empirical prediction and rationalization of protein pKa values. Proteins 2005, 61 (4), 704–721. DOI: 10.1002/prot.20660.

(67) Dolinsky, T. J.; Nielsen, J. E.; McCammon, J. A.; Baker, N. A. PDB2PQR: an automated pipeline for the setup of Poisson-Boltzmann electrostatics calculations. Nucleic Acids Res 2004, 32 (Web Server issue), W665–667. DOI: 10.1093/nar/gkh381.

(68) Shang, J.; Ye, G.; Shi, K.; Wan, Y.; Luo, C.; Aihara, H.; Geng, Q.; Auerbach, A.; Li, F. Structural basis of receptor recognition by SARS-CoV-2. Nature 2020, 581 (7807), 221–224. DOI: 10.1038/s41586-020-2179-y (acccessed 2023-08-14T13:12:54).

(69) Kollman, P. A.; Massova, I.; Reyes, C.; Kuhn, B.; Huo, S.; Chong, L.; Lee, M.; Lee, T.; Duan, Y.; Wang, W.; et al. Calculating structures and free energies of complex molecules: combining molecular mechanics and continuum models. Acc Chem Res 2000, 33 (12), 889–897. DOI: 10.1021/ar000033j From NLM.

(70) Genheden, S.; Ryde, U. The MM/PBSA and MM/GBSA methods to estimate ligand-binding affinities. Expert Opin Drug Discov 2015, 10 (5), 449–461. DOI: 10.1517/17460441.2015.1032936 From NLM.

