## Supplementary information for "The accomplices: Heparan sulfates and N-glycans foster SARS-CoV-2 spike:ACE2 receptor binding and virus priming"

### SI appendix for:

[1] Heidelberg University, Heidelberg, Germany
[2] Zentrum für Molekulare Biologie (ZMBH), DKFZ-ZMBH-Alliance, Heidelberg University, Heidelberg, Germany
[3] Molecular and Cellular Modeling Group, Heidelberg Institute for Theoretical Studies (HITS), Heidelberg, Germany
[4] Department of Virology, Aggeu Magalhães Institute, Oswaldo Cruz Foundation, FIOCRUZ, Recife, PE, Brazil
[5] Department of Fundamental Chemistry, Federal University of Pernambuco, UFPE, Recife, PE, Brazil
[6] Current affiliation: NEC OncoImmunity AS, Gaustadalléen 21, 0349, Oslo, Norway
[7] Macromolecular Interaction Analysis Unit, Section of Experimental Oncology and Immunology, Department of Molecular and Translational Medicine, Brescia, Italy
[8] Interdisciplinary Center for Scientific Computing (IWR), Heidelberg University, Heidelberg, Germany
[9] Lead contact

**This PDF file includes:**

1. Supporting Material and Methods p.1
2. Supporting Figures: S1-S18 p.4
3. Supporting Tables: S1-S3 p.27
4. Supporting Movies: S1-S2 p.30
5. Supporting References p.31

**Other supporting materials for this manuscript include the following:**

Conventional MD simulation replica trajectories of open spike:ACE2 complexes in the absence and the presence of GAG chains are available on the BioExcel COVID-19 platform https://bioexcel-cv19.bsc.es/#/ with the identifiers MCV1901746 and MCV190174, respectively. All data that support the findings of this study are publicly available in the Zenodo repository: https://zenodo.org/records/10617411. All software used is available as described in the Methods. A preprint has been released in bioRXiv at https://www.biorxiv.org/content/10.1101/2024.02.05.578888 under a CC-BY-NC-ND 4.0 International license.

### 1. Supporting Material and Methods

**Conventional MD simulation**. All conventional MD simulations were carried out using the Amber 20 software[1]. The Amber ff14SB[2], GLYCAM-06j[3] and general AMBER (GAFF)[4] force fields were used to assign the parameters. The starting configurations were solvated in the center of a cubic periodic solvent box using a TIP3P water model[5] with at least 10 Å between the solutes and the edges of the box. Na+ and Cl- ions were added to neutralize the systems and to immerse them in solvent with an ionic strength of 150 mM. The two final models of the spike:ACE2 and spike:ACE2:GAGs complexes consist of 873223 and 872404 atoms, respectively. Energy minimization was performed in 14 consecutive energy minimization steps, each of 100 steps of steepest descent followed by 900 steps of conjugate gradient, with decreasing positional restraints from 1000 to 0 kcal/mol $A^2$ on all the atoms of the systems excluding waters, counterions, and hydrogens, with a cutoff for Coulombic and Lennard-Jones nonbonded interactions of 8 Å. Subsequently, the systems were subjected to two consecutive steps of heating, each of 100,000 time steps, from 10 to 100 K and from 100 to 310 K in an NVT ensemble with a Langevin thermostat with a friction coefficient of $1ps^{-1}$. Bonds involving hydrogen atoms were constrained with the SHAKE algorithm[6], and a 2 fs time step was used. The systems were then equilibrated at 310 K in four consecutive steps of 2.5 ns each in the NPT ensemble with a Langevin thermostat with a friction coefficient of $1ps^{-1}$ and a Berendsen barostat with random velocities assigned at the beginning of each step. Then multiple independent 1μs MD production runs for the systems without (4x) and with (6x) GAGs were carried out starting from randomly chosen restart files with randomly assigned velocities from the last 5 ns of equilibration. Notably, although one spike:ACE2:GAGs complex was modelled, the extraction of 6 restart files chosen randomly from the last 5ns of equilibration resulted in independent starting structures with somewhat different initial positioning of the GAGs in the basic binding groove for running the conventional MD simulations. Short-ranged nonbonded interactions (Coulombic and Lennard-Jones) were evaluated using the default cutoff parameter of 8 Å and long-range electrostatic interactions were treated with cubic spline switching and the Particle Mesh Ewald approximation [7] as recommended for the ff14SB force field [2]. The PME grid dimensions were automatically computed (Amber20 manual (https://ambermd.org/doc12/Amber20.pdf), p. 662). Temperature was maintained at 310 K using the Langevin thermostat while pressure was controlled using the Berendsen barostat with a 0.4 ps relaxation time. Coordinates were written at intervals of 100 ps. Simulations were carried out on in-house GPU-nodes.

**RAMD simulations.** The last snapshot of each conventional MD replica simulation was used as the starting point for the random acceleration molecular dynamics (RAMD) simulations[8, 9]. For each of these snapshots, the Amber format files (coordinates and topology) were converted to GROMACS format using the AnteChamber Python Parser interfacE (acpype) script[10]. The periodic box was enlarged by 70 Å along the z-axis to allow the glycoproteins to separate during the simulations and refilled with TIP3P water molecules and ions to maintain 150 mM NaCl concentration, reaching 900170 and 1061017 atoms for the spike:ACE2 and spike:ACE2:GAGs systems, respectively. Each system was geometry-optimized using up to 50,000 steepest descent steps until the maximum force was lower than 100 $kJ.mol^{-1}nm^{-1}$. The system was then heated in the NVT ensemble for 100 ps to a target temperature of 310 K using a time step of 1 fs, a Langevin thermostat with a friction coefficient of 0.5 $ps^{-1}$, and position restraints of 1000 $kJmol^{-1}nm^{-2}$ on the solute atoms. The system was then equilibrated with no restraints for 1 ns in the NPT ensemble using the Langevin thermostat with a friction coefficient of 0.5 $ps^{-1}$ and the Berendsen barostat with a 0.4 ps relaxation time. The equilibration was extended for a further 2.5 ns using a 2 fs time step. For all equilibration simulations, the GROMACS 2020.3[11] engine was used. After equilibration, 15 dissociation trajectories were generated for each replica using the RAMD method implemented in GROMACS-RAMD (https://github.com/HITS-MCM/gromacs-ramd). During the equilibration and dissociation simulations, short-ranged nonbonded interactions (Coulombic and Lennard-Jones) were treated using a 12 Å cut-off, and electrostatic interactions were treated with the fast smooth Particle-Mesh Ewald scheme[7]. Upon testing additional forces with random initial orientations and magnitudes of 120, 167, 239, 598, 789, and 1196 kcal/mol·Å, the lowest force magnitude of 120 kcal/mol·Å was adopted. This force magnitude ensured that most of the replica simulations, both with and without GAG chains bound, dissociate within the 24-hour computing time limit of our in-house compute-cluster while avoiding significant distortion of the proteins. The movement of ACE2 was monitored at intervals of 50 time steps and, if ACE2 did not move more than 0.025 Å in one of these intervals, the direction of the force was changed randomly and the simulation continued. The simulations were stopped when the proteins had dissociated such that the distance between the COMs of spike and ACE2 exceeded 120 Å. For the RAMD simulations, the leap-frog integrator was used with a 2 fs timestep. The pressure (1 atm) and temperature (310 K) were kept constant using the Parrinello-Rahman barostat[12] and Nóse-Hoover thermostat[13], respectively.

**Analysis of MD simulations**. VMD[14] was employed for visual inspection of the trajectories and used along with CPPTRAJ[15] from AmberTools20[1] for quantitative analysis.

*Root mean square deviation (RMSD) and Root mean square fluctuation (RMSF)* were computed using CPPTRAJ[15] for all C-alpha atoms of the spike subunits - $S_A$, $S_B$, $S_C$ - as well as for ACE2 and for all the carbon, oxygen, sulfur and nitrogen atoms of each GAG chain (GAG-1, GAG-2, GAG-3) (Fig. S1). The RMSF of the spike $S_C$ RBD with and without GAG chains bound was calculated for the C-alpha atoms of residues 341-521 (Fig. S7).

*Hydrogen bond (H-bond) analysis* was performed with VMD[14] using a single trajectory approach and subsequently computing the statistics (average and standard deviations) over the replica simulations of each system (Fig. 2A,5B-C and S2-3, S9, S12, S17). H-bonds were computed using the default parameters (donor hydrogen-acceptor distance of less than 3 Å and D-H…A angle cut off maximum of 20°) for the converged parts of the trajectories (400 ns to 1 microsec) or for the last 400ns in association with FDA analyses or on the last 100 ps of the RAMD trajectories and are reported only when present in at least 50% of the replicas (i.e. in at least 2 and 3 replicas for systems without and with GAGs, respectively). Occupancy was determined by counting the number of frames in which a specific hydrogen bond was formed with respect to the total number of frames.

*- H-bonds of GAGs* were computed between each GAG chain and the spike and ACE2 glycoproteins (Fig. S2 and S12). Given the hydrophilic nature of GAGs, H-bonds are expected to represent the major interactions established with the amino acid residues. Therefore, no further interactions were computed. H-bonds were computed using the default parameters and a cut-off occupancy level of 10% was used.

*- H-bonds between protein residues* were computed for amino acid residues 417-505 and 19-393 of the spike $S_C$ RBD and ACE2 RBD, respectively, using a cut-off occupancy of 10% (Figs. 2A, S3A, and S12B), while the number of contacts during the trajectories for Fig. S3A is reported in Fig. S3B.

*- H bonds of N*-glycans were computed for spike $S_C$ N17, N165, N322 and N657 glycans with ACE2 glycoprotein and for ACE2 N90, N322 and N546 glycans with spike glycoprotein (Figs. S9 and S12C) and for $S_B$ N657 with the $S_C$ D439 residue (Fig. S17). Given the polar nature of the monosaccharides, H-bonds are expected to represent the major interactions with the amino acid residues. No further interactions were computed. H-bonds were considered using a cut-off occupancy level of 10%.

*Residue contact probability* (percentage of total simulation time during which residues are in contact) was computed with an in-house *tcl* script for VMD[14] between amino acid residues 417-505 and 19-393 of spike $S_C$ RBD and ACE2 RBD, respectively, using a single trajectory approach and subsequently computing the statistics (average and standard deviations) over the replica simulations of each system (Fig. 2A-B and S4, S12A). Contacts were computed considering a maximum residue-residue contact distance of 3.5 Å for the converged parts of the conventional MD trajectories (from 400ns to 1 microsec) and for the last 100ps of the RAMD trajectories and are plotted in the matrix when identified in at least 1 frame of a trajectory and present in at least 50% of the replicas (i.e. in at least 2 or 3 replicas for systems without or with GAGs, respectively).

*Dynamic cross-correlation matrices* were computed using CPPTRAJ[15] and were based on per-residue Pearson's correlation coefficients (CCs) as derived from the mass-weighted covariance matrices calculated for residues at the interface between the ACE2 RBm and the spike $S_C$ RBm (Figs. 2C and S5).

*Distance and angle calculations* were performed using an in-house *tcl* script with VMD[14] (Fig.3). To define the orientation of the protein, the complex was aligned with the reference structure, perpendicular to the xy-axis. The distance between ACE2 and the spike $S_C$ NTD was computed between the COMs of ACE2 (residues 18-616) and the $S_C$ NTD (residues 16-271). The alpha and beta angles were computed between the z-axis and the vectors $\boldsymbol{v_1}$ and $\boldsymbol{v_2}$. Vector $\boldsymbol{v_1}$ spans from the COM of the up-RBD of the $S_C$ subunit (residues 341-521) to the COM of ACE2, whereas vector $\boldsymbol{v_2}$ spans from the COM of the three spike central helices (residues 987-1034) to the COM of the $S_C$ NTD. Distances and angles were computed at each frame along the trajectories as a deviation (positive or negative) from their initial values. The trajectories were aligned to the reference structure at frame 0 of each independent trajectory.

*Essential dynamics (ED)* analysis consisted of Principal Component Analysis (PCA) of the conventional MD simulations. PCA was performed along the individual trajectories with CPPTRAJ[15]. The principal modes of motion were visualized using VMD[14]. The first normalized eigenvectors for the subsystem of spike $S_C$ and ACE2 without and with GAGs bound were plotted along the trajectory, and the direction of motion was defined by visual inspection (Figs. 3E-F and S8).

*Force distribution analysis (FDA)* implemented in a modified version of GROMACS 2020.4[11] available at (https://github.com/HITS-MBM/gromacs-fda) was used to calculate the changes in internal forces in the spike and ACE2 glycoproteins upon binding of GAGs (Fig. 5 and S14-S16). For details on the methodology, see Costescu et al.[16]. To perform FDA, the Amber format files (coordinates, topology, trajectory files) were first stripped of water and ions and then converted to GROMACS format using the AnteChamber Python Parser interfacE (acpype)[10] script. FDA was performed on each individual trajectory per condition and the resulting average pairwise forces of the apo state were subtracted from those of the holo state. The networks shown are connected edges for at least 3 residues with force differences above a given threshold (Fig. 5A). The punctual stress is the sum of the absolute values of scalar pairwise forces acting on each atom (Fig. S14-S16). Forces from water and ions were not considered in this analysis due to their rapid interchange of positions. However, the force distribution pattern indirectly reflects the effect of the solvent, as the MD simulations were carried out in an explicit aqueous solution at physiological ion concentration.

*Molecular mechanics-generalized Born surface area (MM/GBSA) energies* were computed with AmberTools20[1] (python script) using a single trajectory approach and then computing the average and standard deviation over the replica simulations of each system (Tab. 2). Binding affinities were calculated for systems without and with GAGs using the last 5000 frames (corresponding to the last 500 ns) of each conventional MD trajectory. Water molecules and ions were treated implicitly (igb=2 and saltconc=0.15 M). All other parameters were assigned default values. MM/GBSA energies were computed for the following systems without and with GAGs: (i) spike:ACE2 including N-glycans; (ii) spike $S_C$ RBD:ACE2 excluding glycans; (iii) spike $S_C$ NTD+RBD:ACE2 including N-glycans (iv) spike $S_C$ NTD+RBD:ACE2 excluding N-glycans. The $S_C$ RBD was defined by residues 341-521, whereas the $S_C$ NTD consisted of residues 16-271 (Tab. 2).

#### 2. Supporting Figures

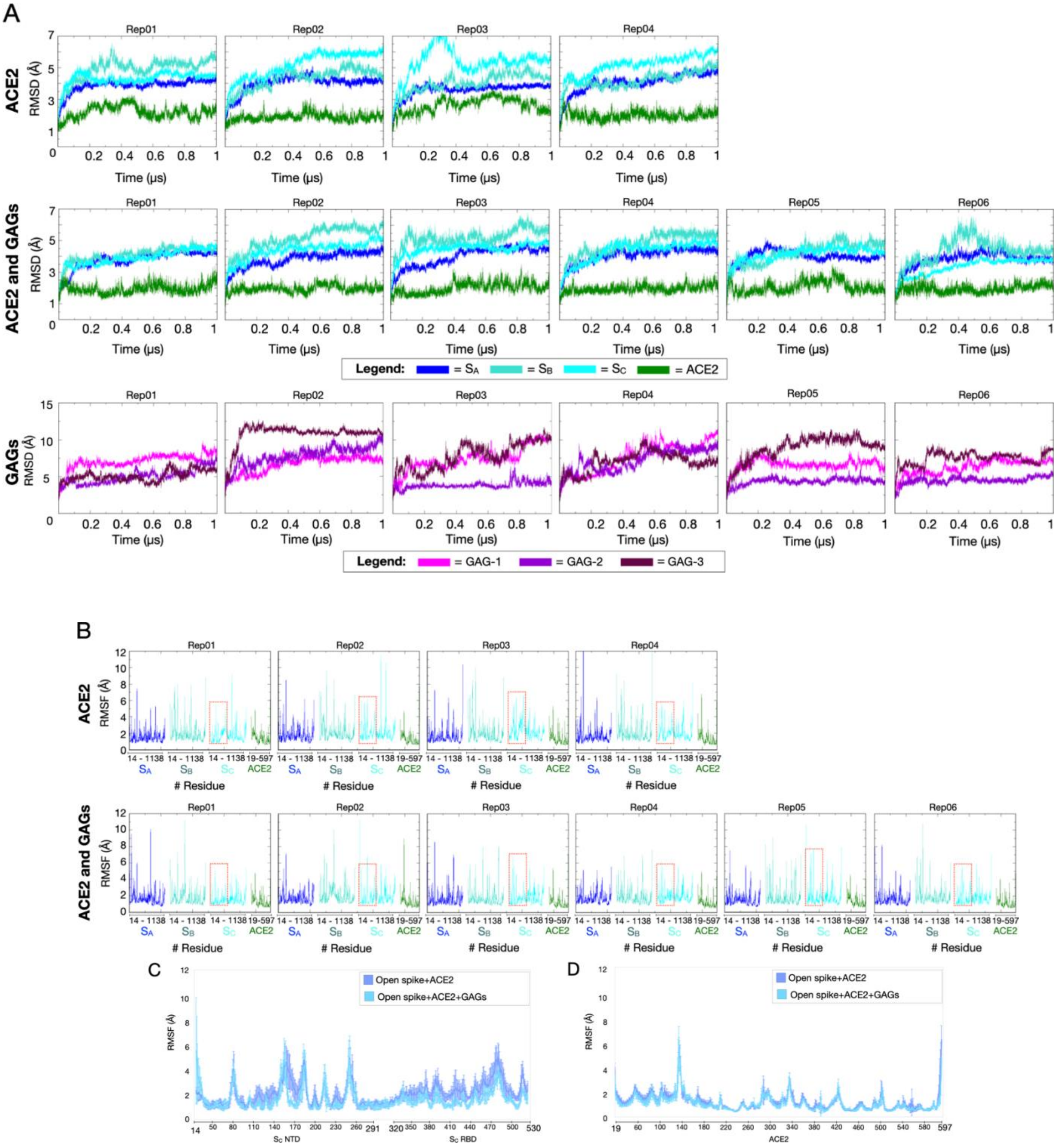


**Fig. S1 Structural convergence and conformational fluctuations of the simulated systems.** (A) Root mean squared deviation (RMSD (Å)) *versus* time (μs) for the replica MD trajectories of the open spike-ACE2 complex in the absence (4 trajectories) and presence (6 trajectories) of GAGs. The RMSD values for the individual spike subunits - $S_A$, $S_B$, $S_C$,-

and ACE2 were calculated for the C-alpha atoms of residues 51-1063 of each spike subunit and of residues 19-597 of ACE2 and are shown in blue, teal, cyan, and green, respectively, whereas the RMSD values of the three GAG chains were calculated for all the C, N and O atoms in all monosaccharides and are shown in magenta, purple and brown. (B) Root mean squared fluctuation (RMSF) (Å) *versus* residue number of the three subunits of the spike homotrimer for the replica MD trajectories of the simulated systems in the absence (4 trajectories) and presence (6 trajectories) of GAGs. The RMSF values for the individual subunits - $S_A$, $S_B$, $S_C$, - and ACE2 were calculated for the C-alpha atoms of residues 14-1138 for each spike subunit and residues 19-597 for ACE2 and are shown in blue, teal, and cyan and green, respectively. The red boxes enclose the NTD (residues 14-291) and RBD (residues 330-530) of the $S_C$ subunit (also see Fig. S4). The average and standard deviation of the RMSF values for the $S_C$ NTD and RBD (C) and ACE2 (D) over the replica MD trajectories of the simulated systems in the absence (4 trajectories) and presence (6 trajectories) of GAGs are plotted per residue along the sequences.

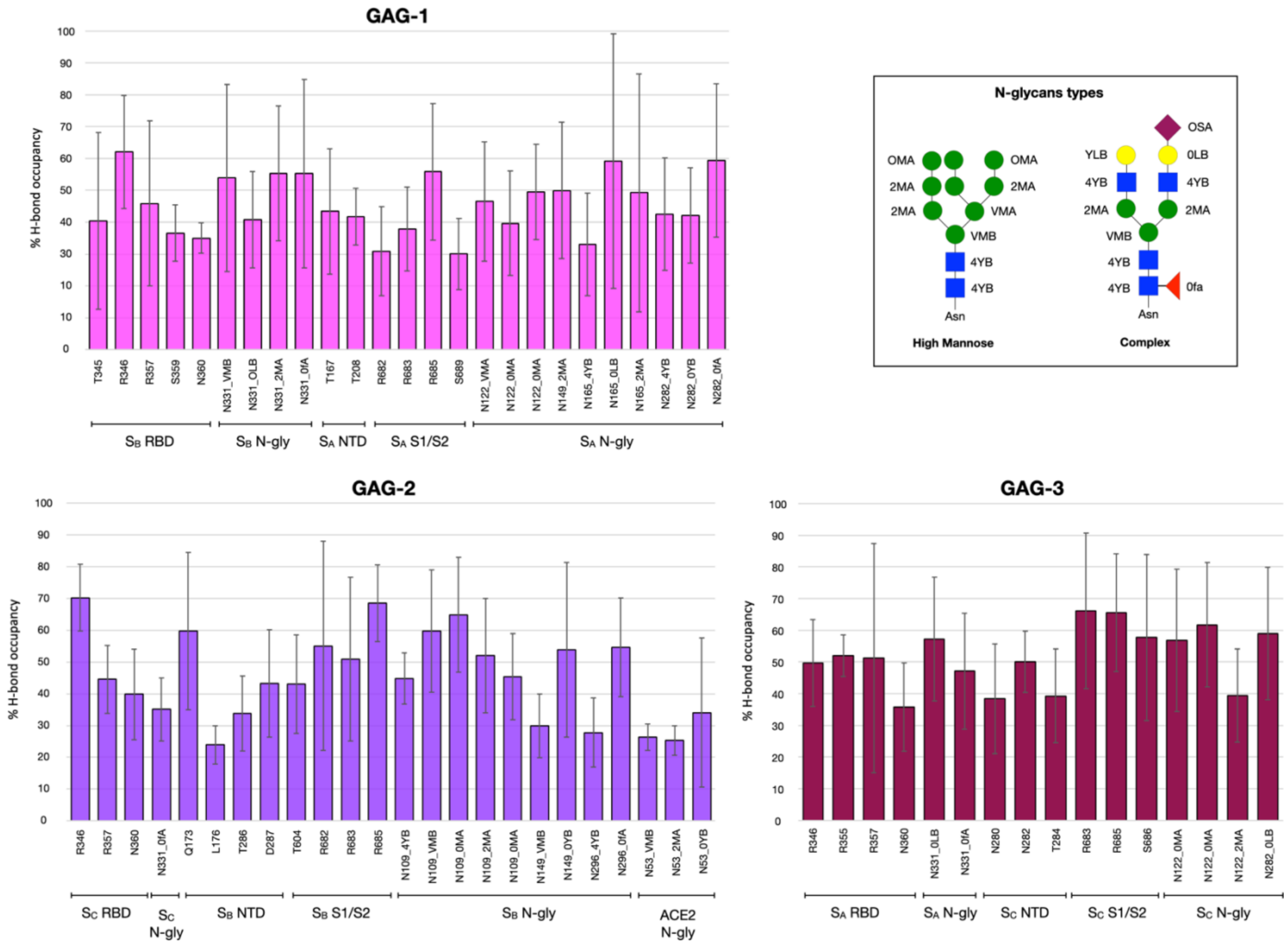


**Fig. S2 H-bond interactions between GAG polysaccharides and spike or ACE2 glycoproteins.** The trajectory-averaged H-bond occupancies were calculated for the last 200ns of each trajectory from conventional MD simulations. Two residues were considered in contact when the distance between their heavy atoms was within 3.2 Å and persistent for more than 80% of the trajectories analyzed. H-bonds were computed independently for each GAG chain (GAG-1, GAG-2, and GAG-3). The inset shows the nomenclature for the monosaccharides for two representative N-glycan types (high mannose and a complex type), drawn according to the canonical Symbol Nomenclature of Glycans (SNFG), and includes the nomenclature used to name each of the monosaccharides in the GAGs (GAG-1: $S_B$ N-gly and $S_A$ N-gly; GAG-2: $S_C$ Ngly, $S_B$ N-gly and ACE2 N-gly; GAG-3: $S_A$ N-gly and $S_C$ N-gly). GAG-2 binds to the up-RBD and establishes interaction with ACE2 (N53 glycan), whereas GAG-1 and GAG-3 do not interact with ACE2.

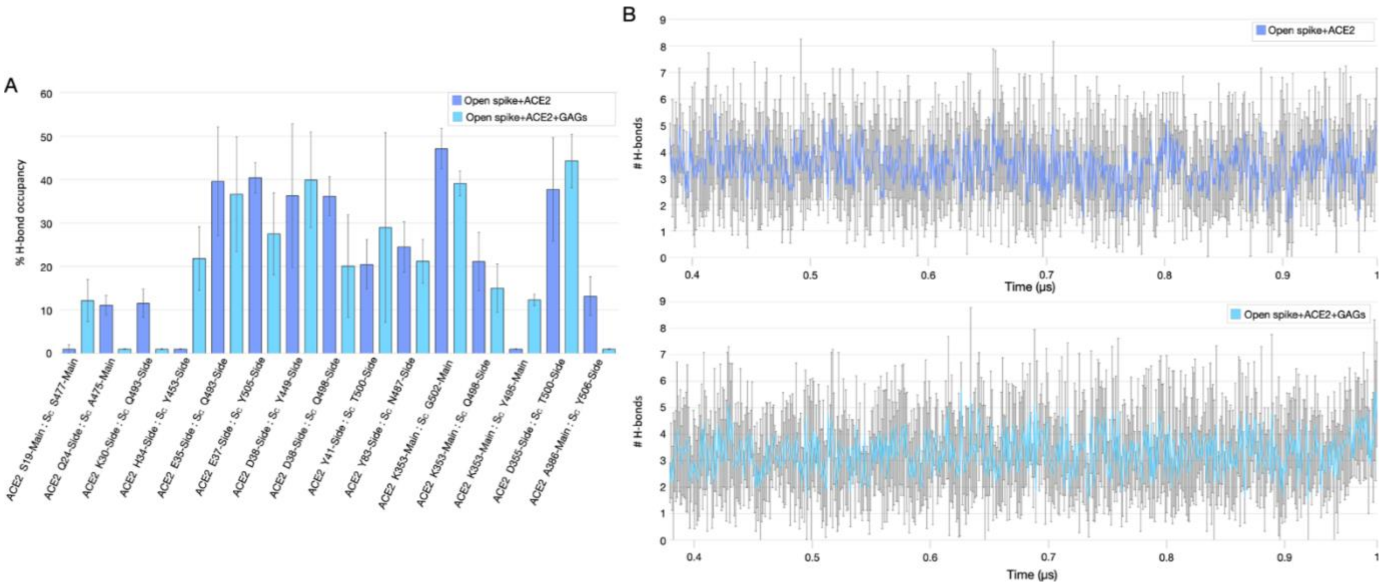


**Fig. S3 H-bonds at the interface between spike $S_C$ RBm and ACE2 RBm.** (A) H-bond occupancy computed for the last 400ns of each trajectory and (B) number of H-bonds during the simulations between the spike $S_C$ RBm and the ACE2 RBm in the absence (blue) and presence (cyan) of GAGs. For comparison with interactions in crystal structures, see Tab. S1.

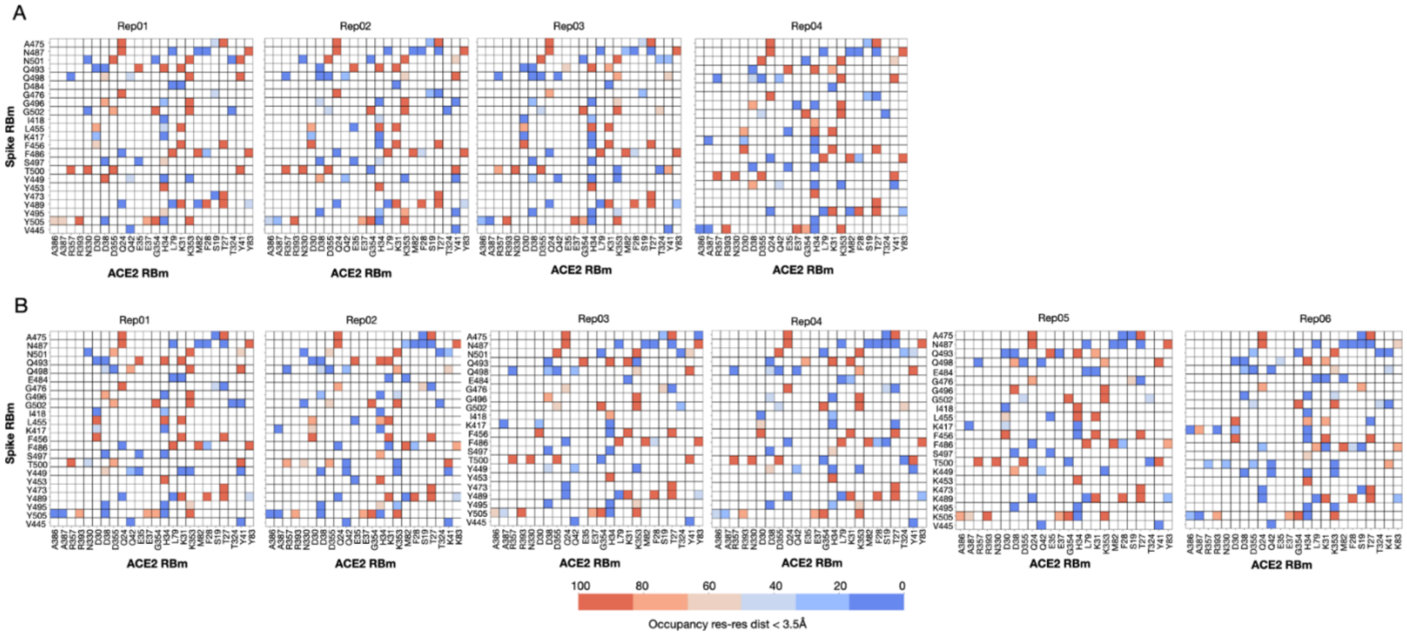


**Fig. S4 Residue-residue interactions at the interface between the spike $S_C$ RBm and ACE2 RBm.** Contact maps between spike $S_C$ RBm and ACE2 RBm residues were computed for the last 400ns of each trajectory in (A) the absence and (B) the presence of GAGs. Two residues were considered in contact when the distance between their heavy atoms was within 3.5 Å and persistent for more than 80% of the trajectories analyzed. Contact values (percentage of persistency along the trajectory) range from 0 (blue, no interaction) to 100 (red, interaction over the entire trajectory). White squares indicate residues that are far from each other. For comparison with interactions in crystal structures, see Tab. S1.

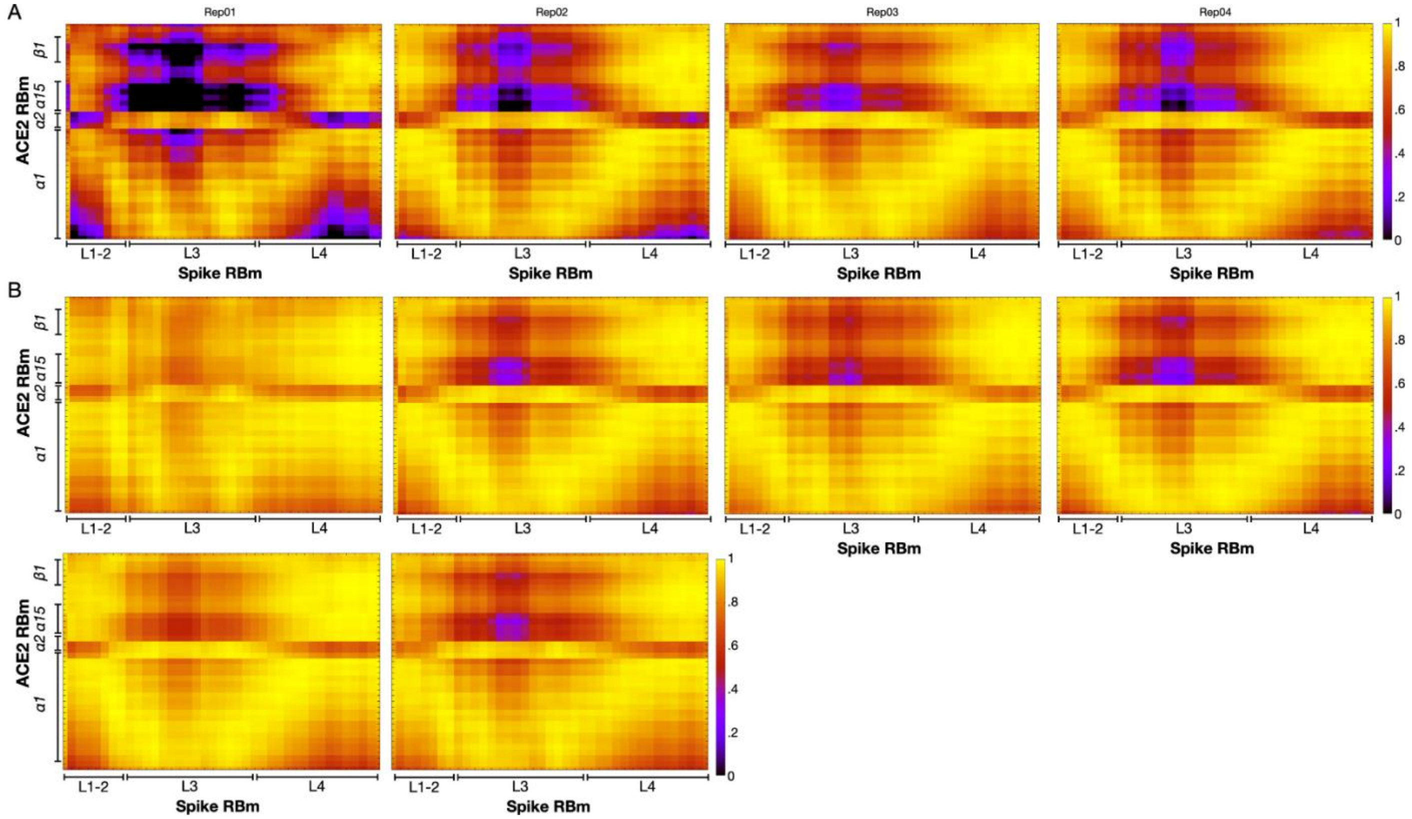


**Fig. S5 GAG-binding results in increased cross-correlated motion of the spike $S_C$ RBm and the ACE2 RBm.** The dynamic cross-correlation matrix (DCCM) based on per-residue Pearson's correlation coefficients (CCs) as derived from the mass-weighted covariance matrix calculated for residues at the interface between ACE2 RBm and spike $S_C$ RBm for the replica MD trajectories of the complexes in, respectively, (A) the absence (4 trajectories) and (B) the presence (6 trajectories) of GAGs. CC values range from 0 (black, uncorrelated motion) to +1 (yellow, correlated motion). For the location of the regions selected for the analysis in the 3D structure, see Fig. S6.

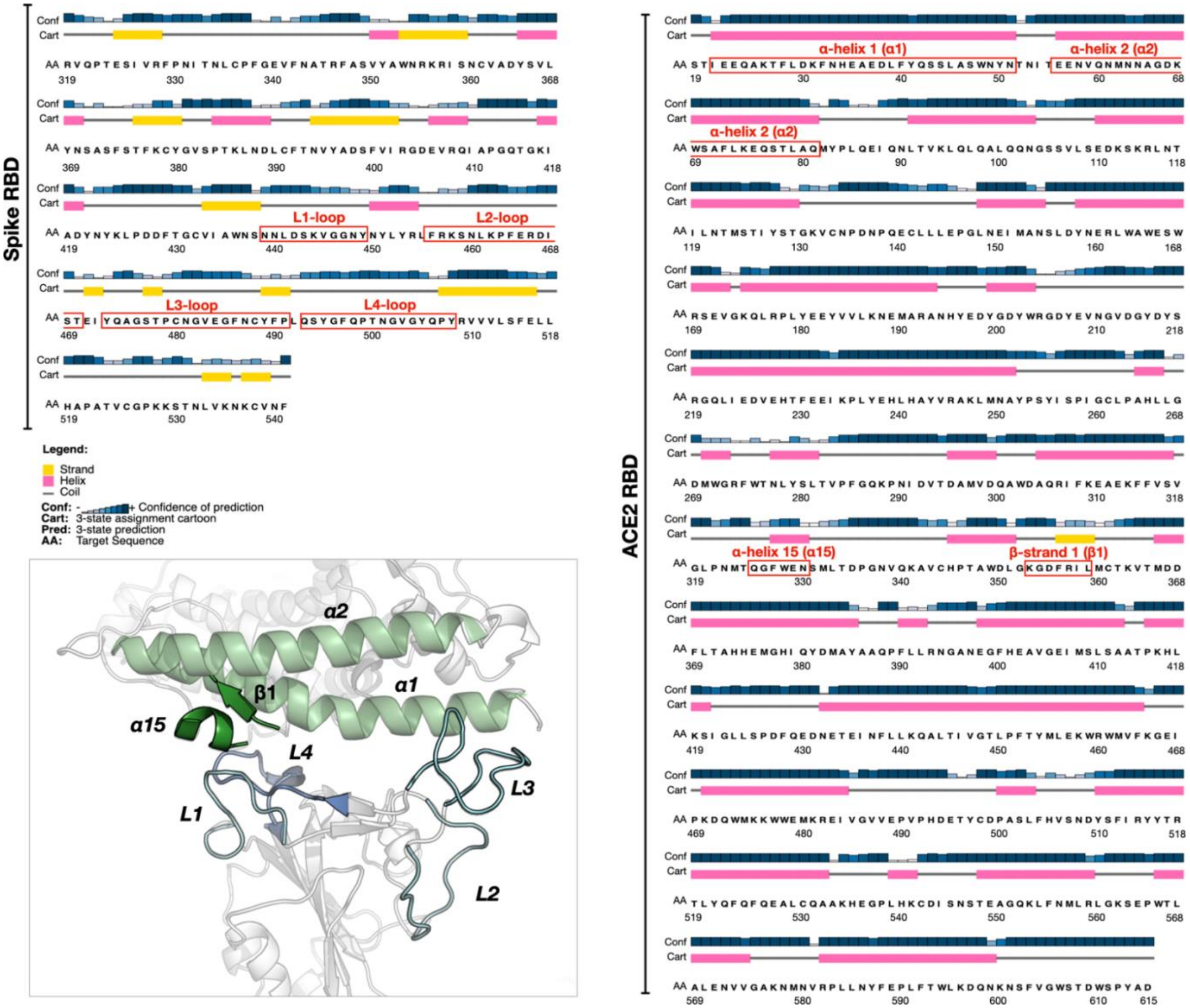


**Fig. S6 Sequences of spike RBD and ACE2 and the structure of their interface**. The secondary structures of the Wuhan spike and ACE2 RBDs are reported as predicted using Psipred (http://bioinf.cs.ucl.ac.uk/psipred/). The spike and ACE2 regions considered for performing the dynamic cross-correlation matrix (DCCM) analysis are highlighted and labelled on the 3D structure (see Fig. S5). In detail, the interfacial residues of ACE2 α1 (21-51), ACE2 α2 (56-81), ACE2 α15 (315-330), and ACE2 β1 (353-359) are shown as cartoon representations and colored green, while the interfacial residues of spike L1-loop (441-449), L2-loop (456-470), L3-loop (472-490) and L4-loop (493-508) are shown as cartoon colored cyan.

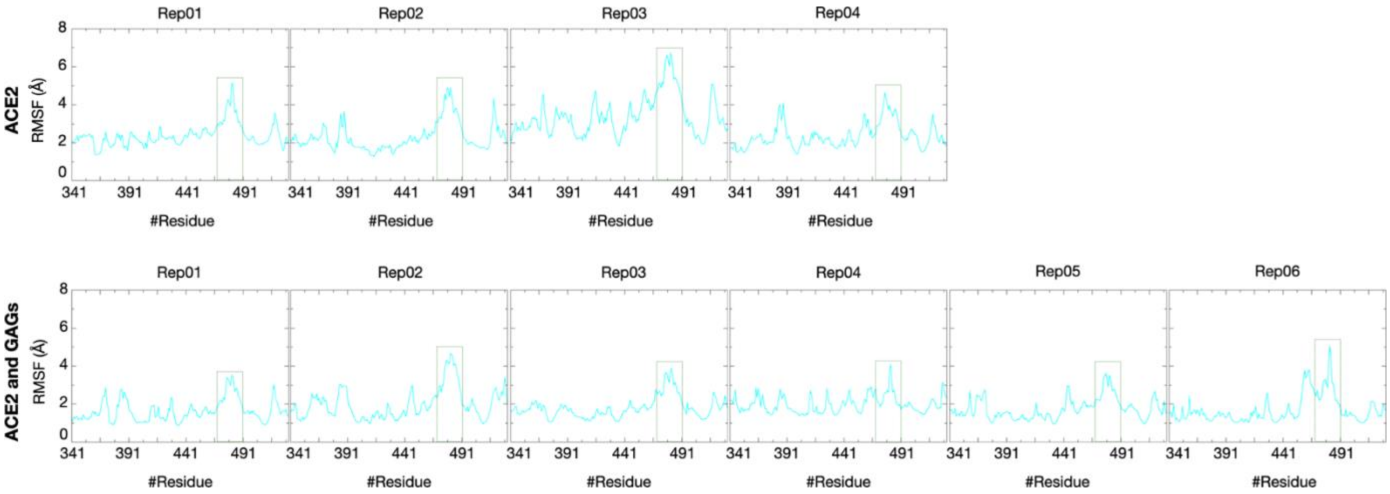


**Fig. S7 Rigidification of spike $S_C$ up-RBD and L3-loop upon GAG binding.** Root mean square fluctuation (RMSF) (Å) *versus* residue number of the $S_C$ – RBD for the replica MD trajectories of the simulated systems without (4 trajectories) and with (6 trajectories) GAGs. The RMSF values were calculated for the C-alpha atoms of the RBD residues (residues 341-541). The green dashed line boxes highlight the L3-loop (472-490 residues) of the $S_C$ subunit

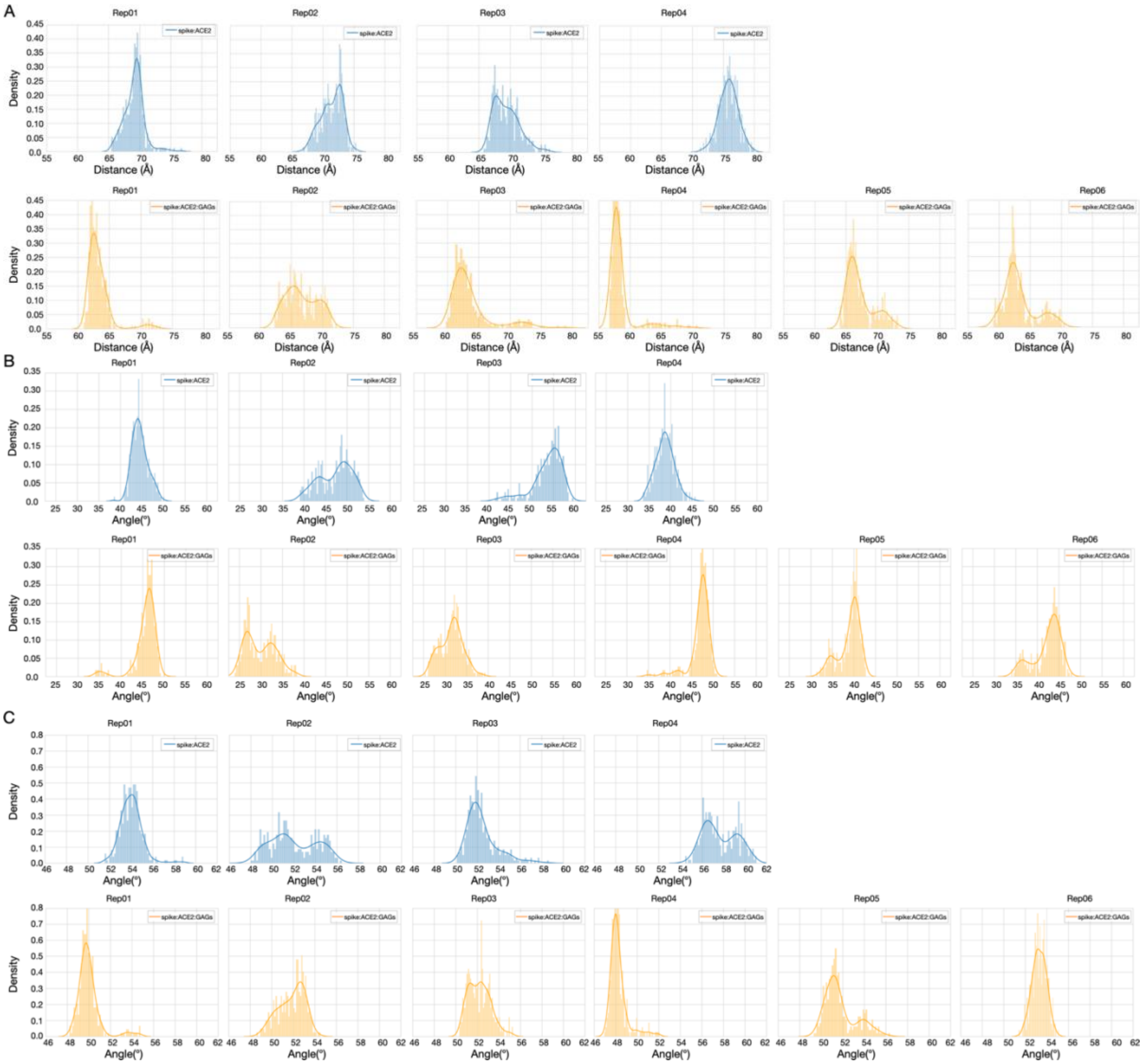


**Fig. S8 Binding of GAG chains brings the ACE2 and spike $S_C$ NTD closer and narrows the distribution of relative orientations of ACE2 and spike as defined by angles α and β.** The distribution of the distances d (A) and angles (B) α and (C) β are computed and shown for the replica simulations for the open spike:ACE2 (4 trajectories - blue) and open spike:ACE2:GAGs (6 trajectories - orange) systems. In the presence of the GAG chains, all replicas show reduced distances (A) and narrower distributions of angles (B-C) although the latter distributions are wider for replica02 and replica03, indicating some differences in the ensembles of conformations in the bending of ACE2 between replicas.

A

Eigenvalues spike $S_C$ subunit-ACE2
Open spike + ACE2 systems

| eigenvalue | Replica 1 | Replica 2 | Replica 3 | Replica 4 |
|---|---|---|---|---|
| 1 | 38595 (3.5x) | 70999 (4.1x) | 76978 (1.5x) | 53023 (2.5x) |
| 2 | 10847 (1.3x) | 17340 (2.4x) | 51059 (3.2x) | 21538 (2.0x) |
| 3 | 8215 (1.1x) | 7216 (1.4x) | 15860 (2.9x) | 10888 (1.5x) |
| 4 | 7314 (1.7x) | 5123 (1.2x) | 5450 (1.5x) | 7063 (2.0x) |
| 5 | 4404 | 4349 | 3547 | 3491 |
| 6 | 2815 | 2994 | 2737 | 2649 |
| 7 | 2396 | 2559 | 1970 | 2381 |
| 8 | 1701 | 1833 | 1489 | 1798 |
| 9 | 1355 | 1522 | 1294 | 1443 |
| 10 | 1159 | 1423 | 950 | 1151 |

C

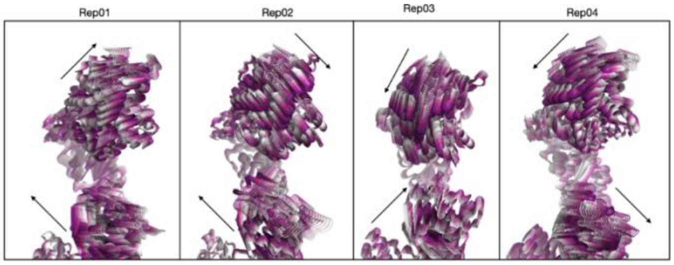


B

Eigenvalues spike $S_C$ subunit-ACE2
Open spike + ACE2 + GAGs systems

| eigenvalue | Replica 1 | Replica 2 | Replica 3 | Replica 4 | Replica 5 | Replica 6 |
|---|---|---|---|---|---|---|
| 1 | 25748 (2.8x) | 71098 (6.3x) | 23939 (1.5x) | 31605 (2.8x) | 39219 (2.2x) | 60564 (6.2x) |
| 2 | 9281 (1.2x) | 11160 (1.3x) | 16127 (1.5x) | 11100 (1.4x) | 17617 (2.6x) | 9756 (1.8x) |
| 3 | 7562 (1.9x) | 8719 (1.6x) | 10612 (2.3x) | 7683 (2.3x) | 6744 (1.5x) | 5468 (1.7x) |
| 4 | 3993 (1.5x) | 5301 (1.7x) | 4576 (1.4x) | 3351 (1.5x) | 4371 (1.4x) | 3219 (1.4x) |
| 5 | 2623 | 3069 | 3236 | 2268 | 3015 | 2270 |
| 6 | 1763 | 2228 | 2669 | 1904 | 1902 | 1989 |
| 7 | 1280 | 2049 | 2004 | 1509 | 1844 | 1492 |
| 8 | 1194 | 1566 | 1602 | 1271 | 1603 | 1448 |
| 9 | 1082 | 1376 | 1155 | 1078 | 1250 | 932 |
| 10 | 846 | 997 | 962 | 945 | 1083 | 840 |

D

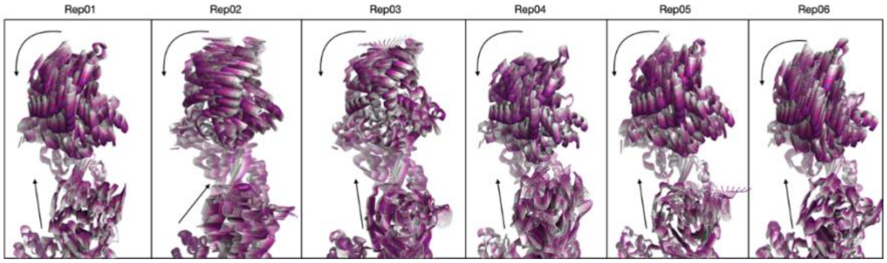


**Fig. S9 Differences in the motion of spike $S_C$ up-RBD bound to ACE2 in the absence and presence of GAGs.** The results of essential dynamics analysis are computed and shown for the $S_C$ subunit bound to ACE2 for the replica simulations for the open spike:ACE2 (4 trajectories) and open spike:ACE2:GAGs (6 trajectories) systems. (A-B) Values of the first eigenvalues for the $S_C$ subunit with the up-RBD and ACE2 for the two simulated systems. Multipliers denote the magnitude of each eigenvalue with respect to the next eigenvalue. (C-D) Structure superpositions showing the direction of motion of the first eigenvector of the $S_C$-RBD bound to ACE2 without (C) or with (D) GAGs. The arrows indicate the main directions of motion. (D) In the presence of the GAG chains, replica02 and replica03 show a somewhat different orientation of motion for ACE2 in the first eigenvector compared to the other replica simulations. Plotting of angles α and β per replica shows that the difference is due to a wider distribution of these angles for these replicas (Fig.S8). Nevertheless, plotting the distance d per replica shows that the distance between COMs of spike $S_C$-NTD and ACE2 is reduced in all replicas (Fig.S8), confirming that the movement results in the reorientation of ACE2 toward spike-NTD.

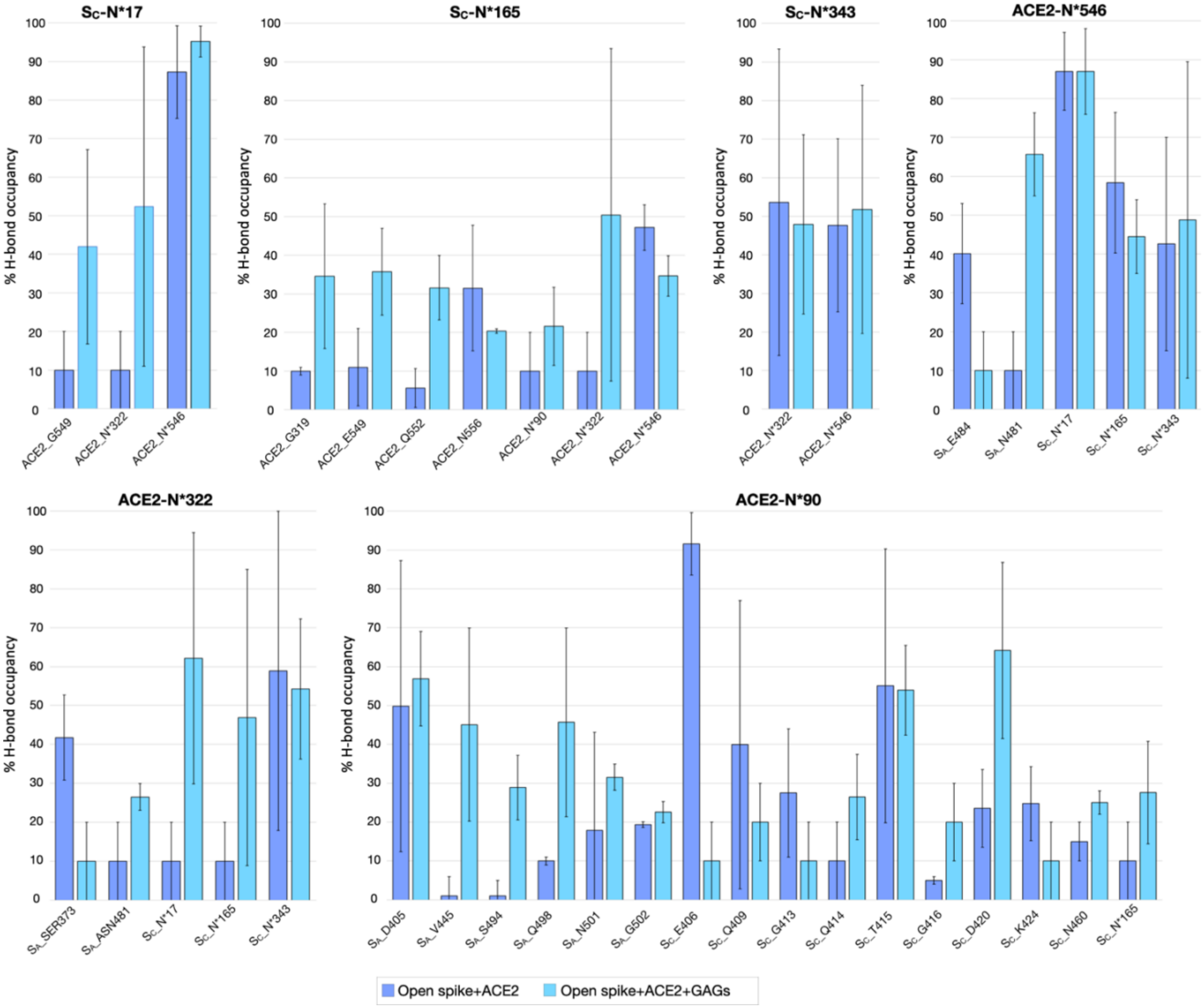


**Fig. S10 H-bonds formed by spike and ACE2 N-glycans.** H-bond interactions of $S_C$ glycans (N17, N165, N343) with ACE2 glycoprotein and of ACE2 glycans (N90, N322, N546) with spike glycoprotein computed for converged parts of the trajectories of the simulated systems in the absence (blue) and presence (cyan) of GAGs. H-bond occupancy across all the replicas is shown (*see methods section for further details*). $S_A$ $S_B$ and $S_C$ denote the three spike subunits. Residues are labelled by single-letter (N*) when an N-glycan is referred to and otherwise by a canonical single-letter code.

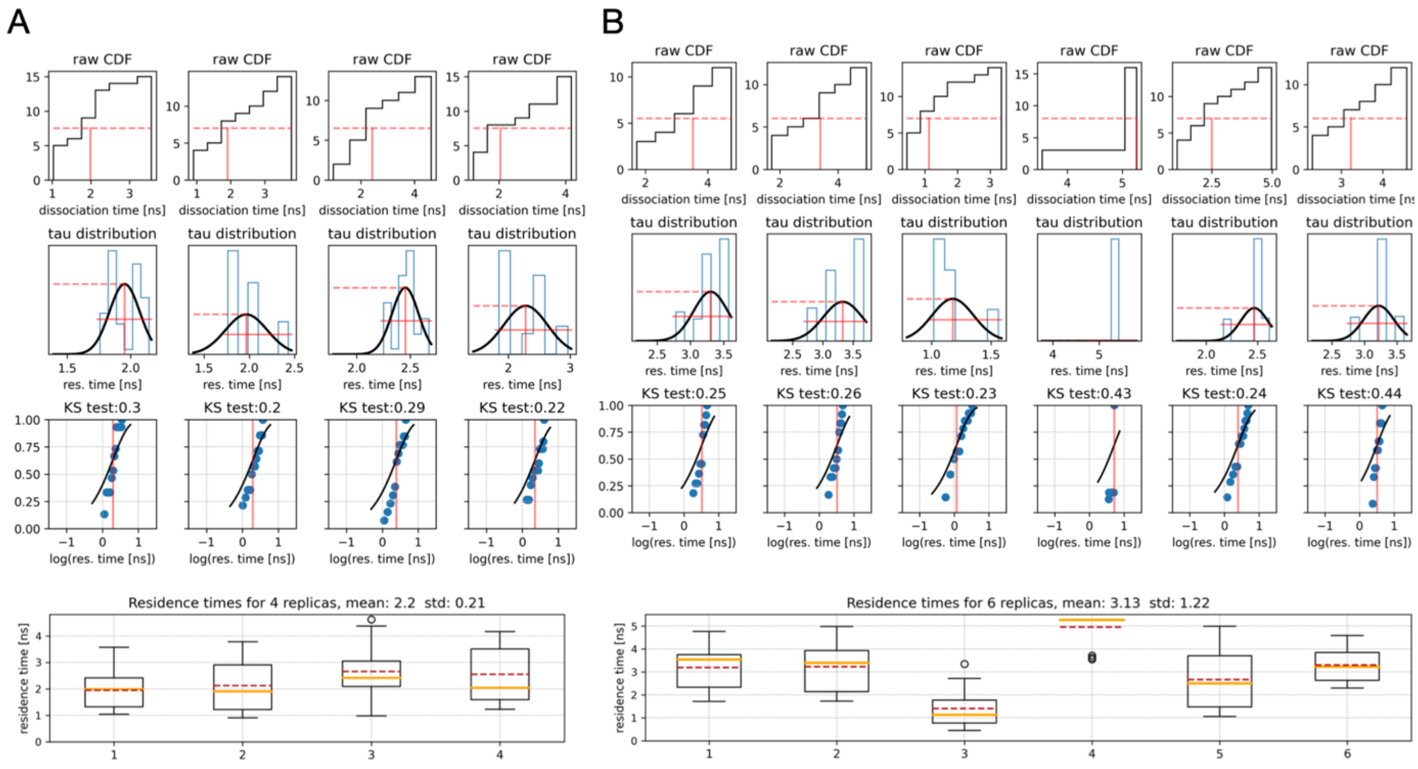

**Fig. S11 RAMD residence times of the open spike-ACE2 complexes derived from RAMD simulations.** The RAMD residence time was calculated from RAMD simulations starting from the last frame of each replica conventional MD simulation and performing 15 replica RAMD simulations for each of the starting structures, resulting in a total of 60 independent replica trajectories for systems in the absence of GAGs (A) and 90 replica trajectories for systems with GAGs (B). The average RAMD residence time is 2.2±0.21 and 3.13±1.22 ns in the absence and presence of GAGs, respectively. Note that these times are orders of magnitude shorter than the expected residence times of the complexes due to the application of an additional random force during the RAMD simulations. Notably, the relative residence times are consistent with the MMGBSA calculations [see Tab. S2].

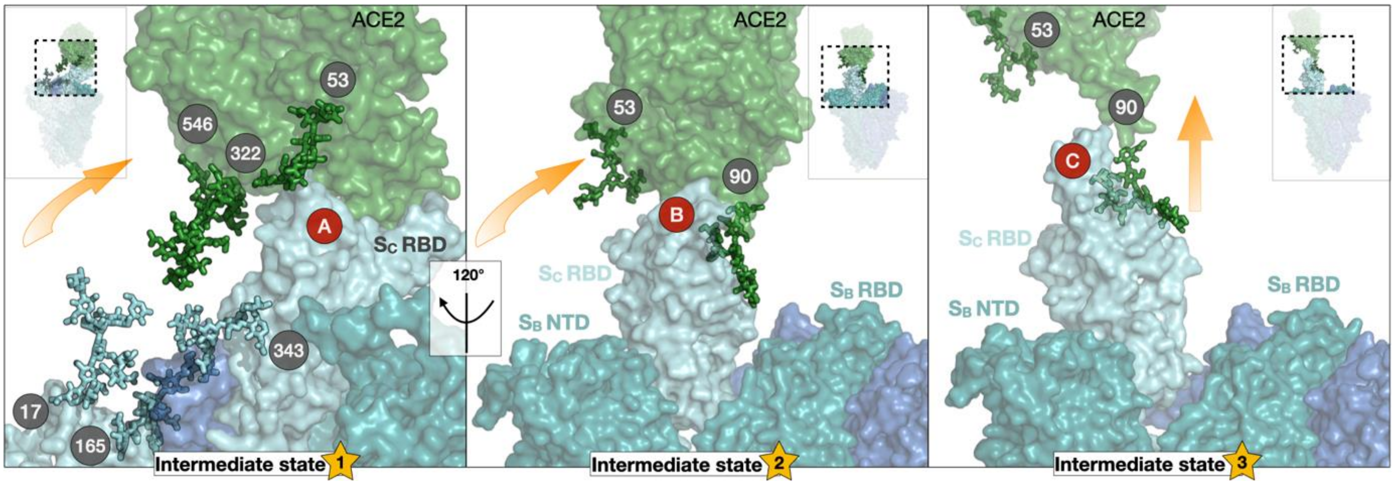


**Fig. S12 Dissociation pathway of ACE2 from the spike in the absence of GAGs (replica 1).** $S_A$, $S_B$, and $S_C$ subunits and ACE2 are shown as surfaces in blue, teal, cyan, and green, respectively. Key N-glycans covalently attached to the spike and ACE2 and involved in the mechanisms described are labeled with grey circles and shown in stick representation, colored according to the subunit to which they are attached. Yellow arrows show the dissociation route of ACE2 as a fan-like movement (rotational and translational) in the first two intermediate states and a translating movement in the third. Red circles schematically highlight the protein-protein dissociation pathway described in detail with the heat map in Fig. S12.

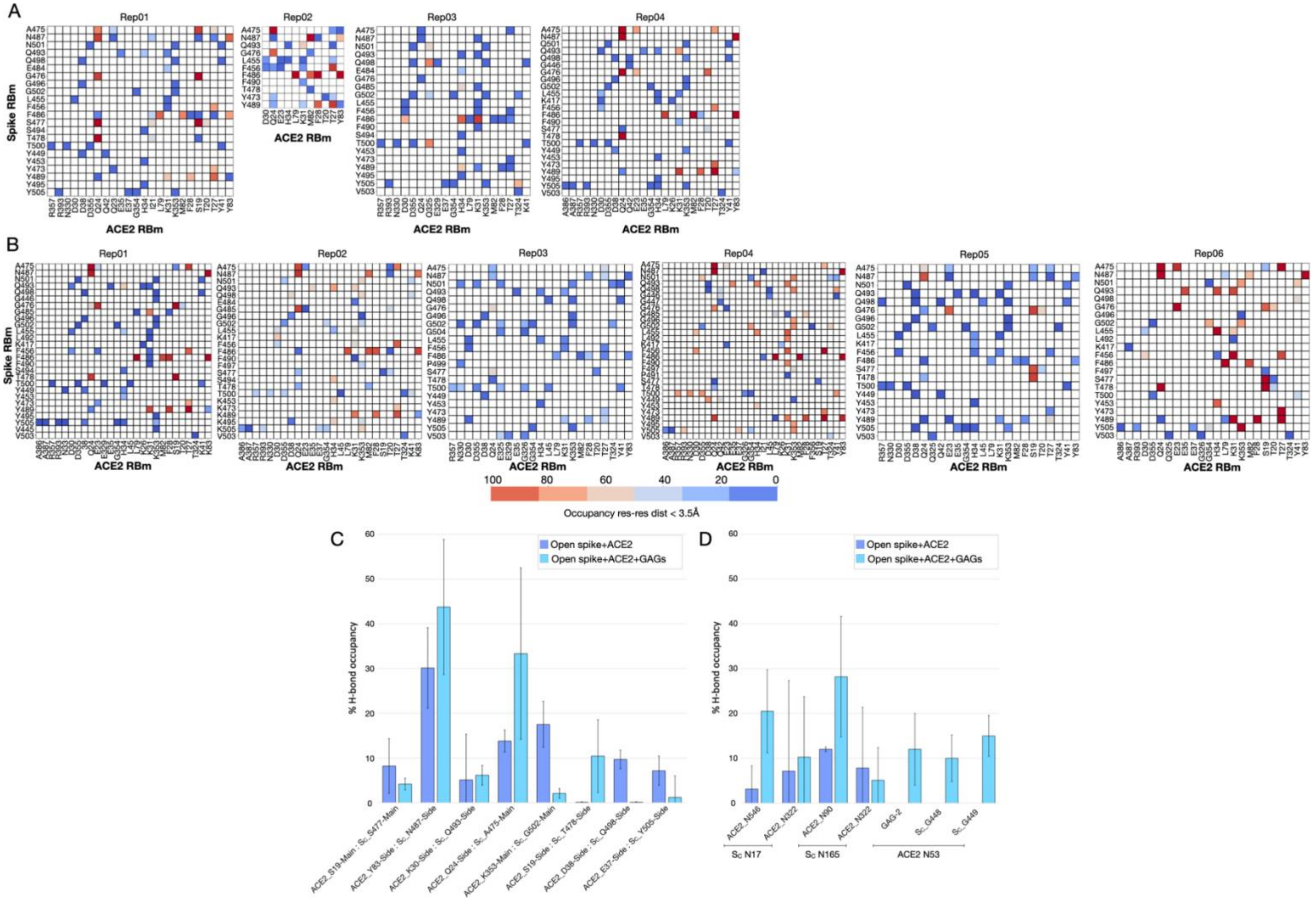


**Fig. S13 Spike:ACE2 interactions in the last 100ps of the RAMD trajectories before dissociation.** (A-B) Contact maps (color scheme as in Fig. S4) and (C) H-bond occupancy for protein-protein interactions between $S_C$ RBm and ACE2 RBm computed for each replica trajectory for the systems without and with GAGs. (D) H-bond occupancy for interactions between $S_C$ and ACE2 N-glycans and between the ACE2 N-glycan N53, GAG-2 and $S_C$ averaged over all trajectories for the systems without (blue) and with (cyan) GAGs (*see methods section for further details*).

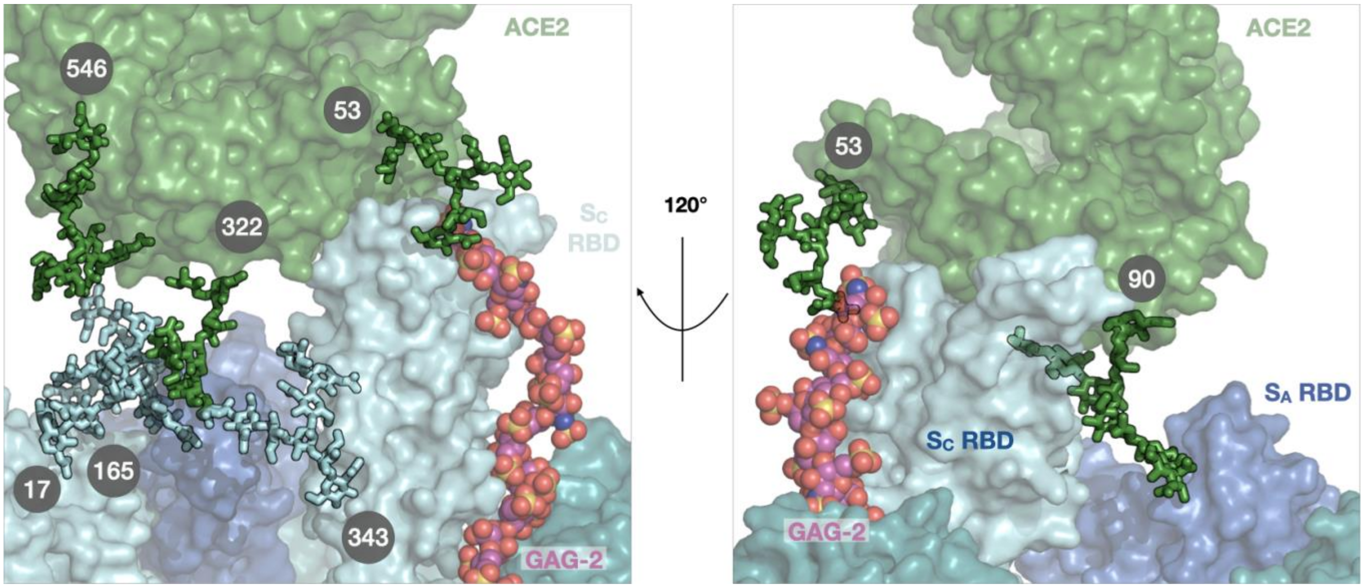


**Fig. S14 Effect of N-glycans on spike:ACE2 stabilization during replica trajectory 3 with three GAG chains bound.** Front (left) and side (right) view of the initial complex. $S_{A,}$ $S_B$, and $S_C$ subunits, and ACE2 are shown as surfaces in blue, teal, cyan, and green, respectively. Key N-glycans covalently attached to the spike and ACE2 are labeled and shown in stick representation, colored according to the subunit to which they are attached. The 31mer GAG-2 chain bound to the up-RBD ($S_C$-RBD) is depicted as spheres colored by elements with magenta carbons. The side view (right) shows that, unlike the other replicas (see Fig. 4A), one branch of the N90 glycan points toward the $S_A$-RBD (blue).

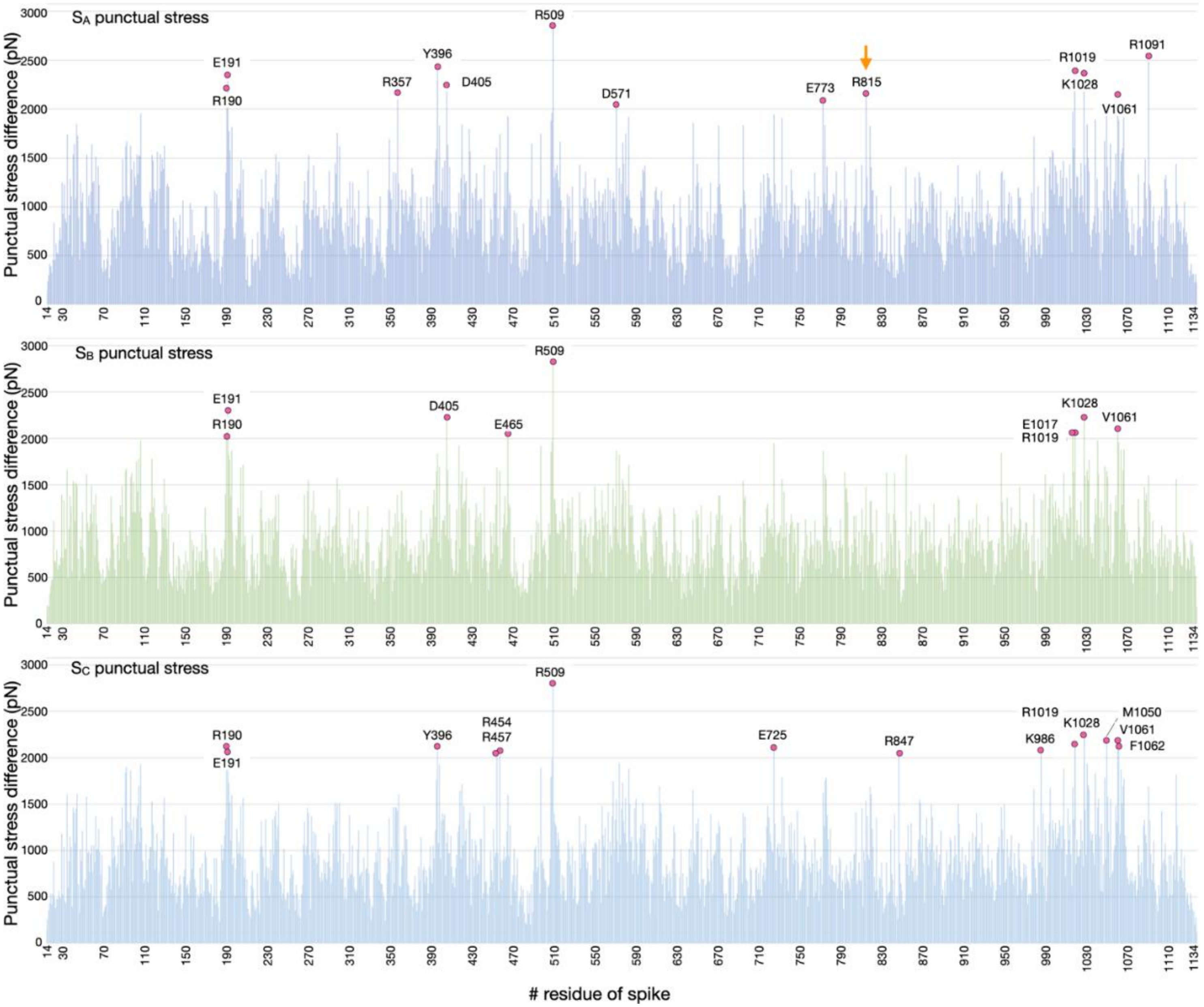


**Fig. S15 Absolute differences in punctual stress between open spike with and without ACE2 bound.** The absolute difference in punctual stress is the sum of the absolute values of scalar pairwise forces acting on each atom computed by merging the last 400ns of each trajectory for the open spike without ACE2 bound (4 replicas), retrieved from Paiardi et al.[17], and for the open spike with ACE2 bound in the absence of GAGs (4 replicas). The spike residues of each subunit are given along the x-axis ($S_A$ in blue, $S_B$ in green, $S_C$ in light blue). N-glycans are included in the computation (see Fig. S17). No rearrangements were identified by visual inspection or H-bond analysis for the residues within the spike NTD (aa 16-291) and RBD (aa 330-530), suggesting that the stress point is due to long-range electrostatic interactions promoted by the negatively charged ACE2 and the charged residues in the surrounding spike domains. Punctual stress residues in the central helix (987-1034) of each spike subunit were investigated by visual inspection and H-bond analysis, allowing the identification of a previously unknown bifurcated $S_A$ E773 - $S_A$ R1019 - $S_A$ E1017 salt-bridge that becomes more transient upon ACE2 binding (see Fig. S16).

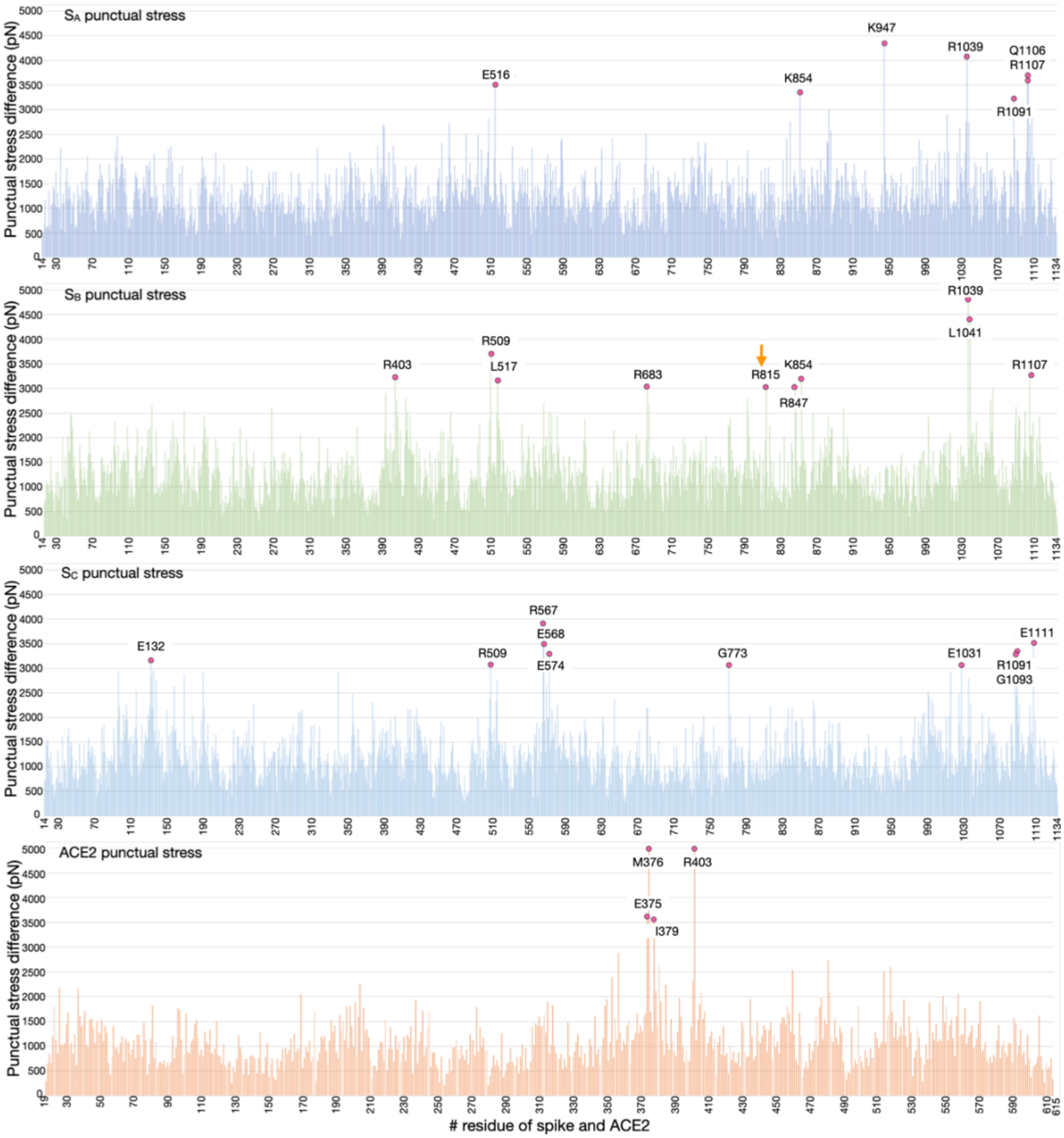


**Fig. S16 Absolute differences in punctual stress in the open spike:ACE2 complex with and without GAGs bound.** The absolute difference in punctual stress is the sum of the absolute values of the scalar pairwise forces acting on each atom computed by merging the last 400ns of each trajectory for the open spike:ACE2 systems without GAGs (4 replicas) and for the open spike:ACE2 with GAGs bound (6 replicas). The residues of spike and ACE2 are given on the x-axis ($S_A$ in blue, $S_B$ in green, $S_C$ in light blue, ACE2 in orange). N-glycans are included in the computation (see Fig. S17). No rearrangements were identified by visual inspection or H-bond analysis for the residues within the S1 domain (19-680), suggesting that the highlighted stress points are due to long-range electrostatic interactions promoted by GAG chains and the charged residues surrounding their binding domain. Visual inspection of the trajectories reveals that the N-glycans surrounding $S_B$ R815 compete with D839 and D843 for binding with R815, either upon binding of ACE2 or ACE2 and GAGs. No relevant rearrangements were identified by visual inspection or H-bond analysis for the residues within the spike central helices (987-1034). However, the bifurcated $S_A$ E773 - $S_A$ R1019 - $S_A$ E1017 salt bridge identified upon ACE2 binding shows reduced occupancy upon ACE2 and GAG binding.

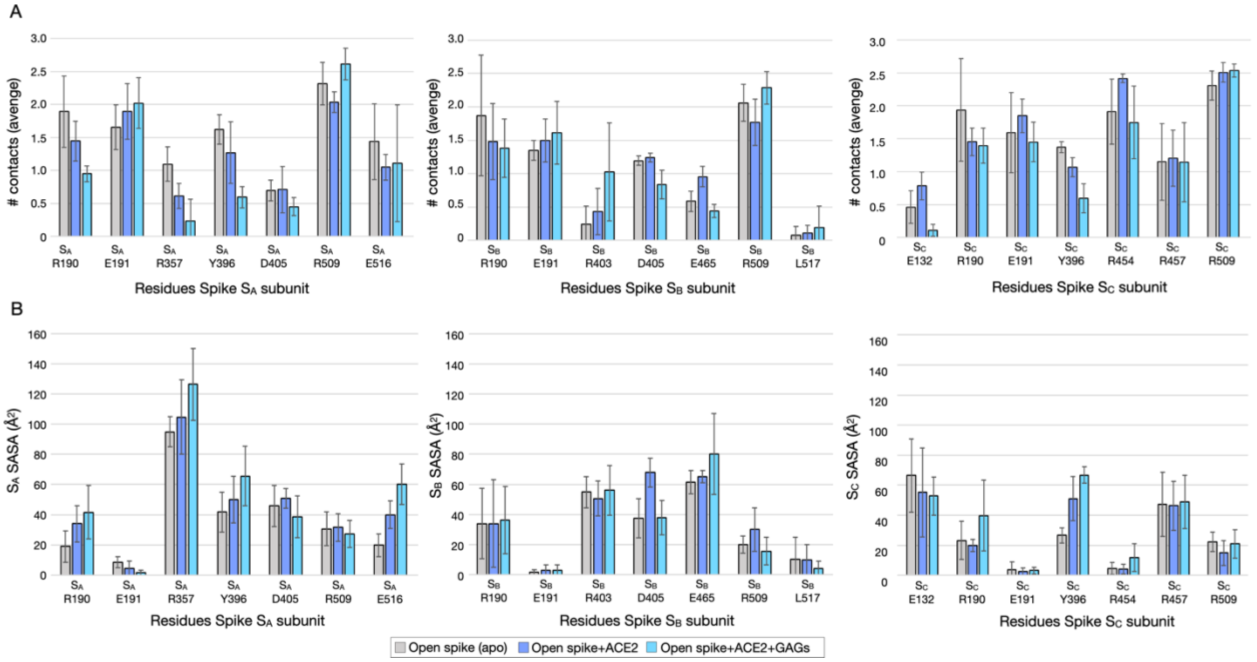


**Fig.S17** – **H-bonds (contacts) and surface exposure to the solvent (SASA) of spike residues in the NTD and RBD.** (A) Average number of contacts and (B) SASA computed for the last 400 ns for spike NTD+RBD residues showing notable differences in punctual stress (Fig. S14-S15) in the absence of ACE2 and GAGs (grey), in the presence of ACE2 (blue) and in the presence of ACE2 and GAGs (cyan). None of the residues was in direct contact with ACE2 or GAGs, suggesting that the stress differences are driven by long-range electrostatic interactions between the negatively charged ACE2 or GAGs and the charged residues of spike in proximity to the binding partners.

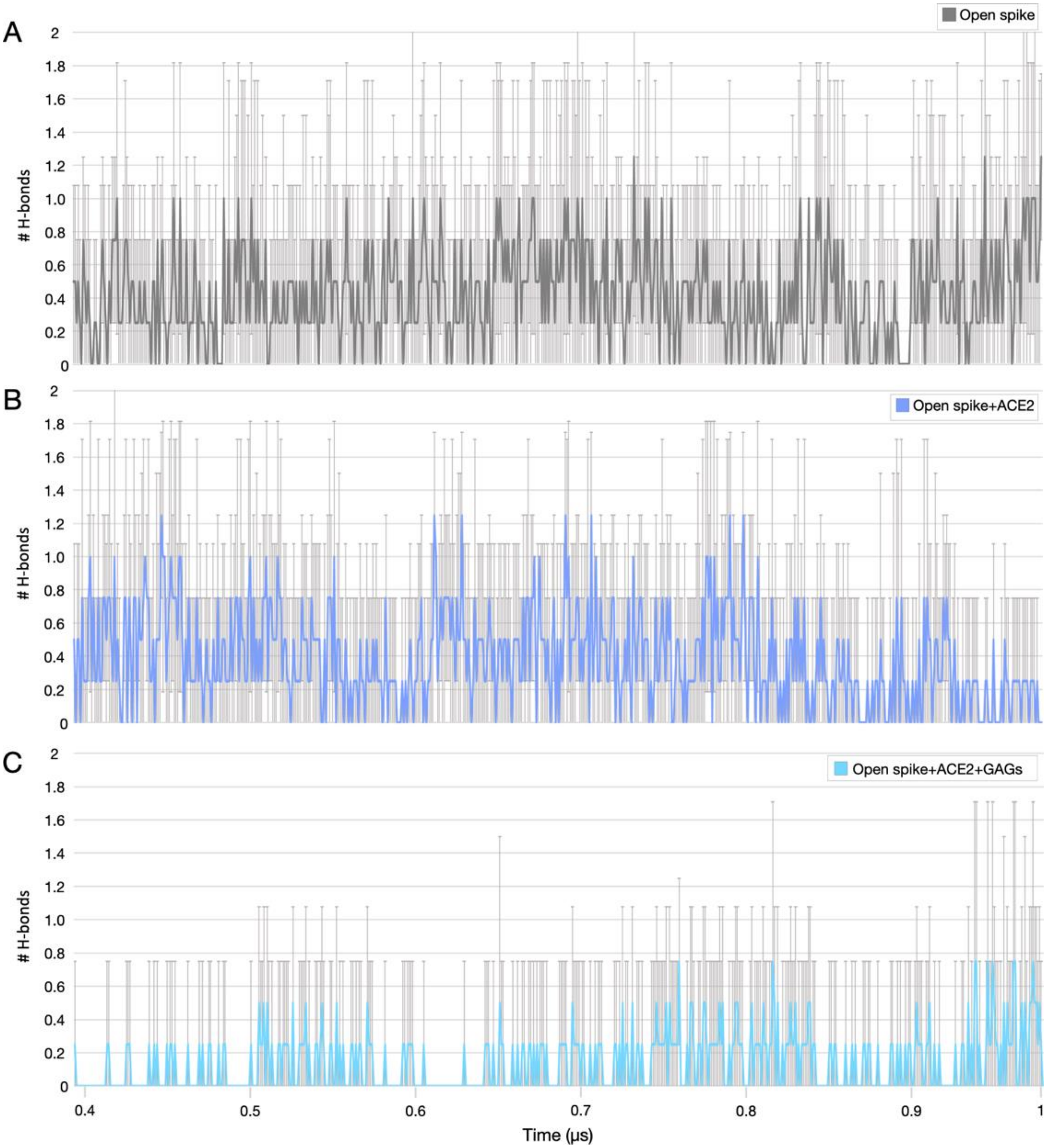


**Fig. S18 H-bonds between $S_A$ E773 - $S_A$ R1019.** The number of H-bonds between $S_A$ E773 - $S_A$ R1019 computed for the last 400ns of each trajectory in (A) the open spike, (B) open spike with ACE2 bound, and (C) open spike with ACE2 and GAGs bound.

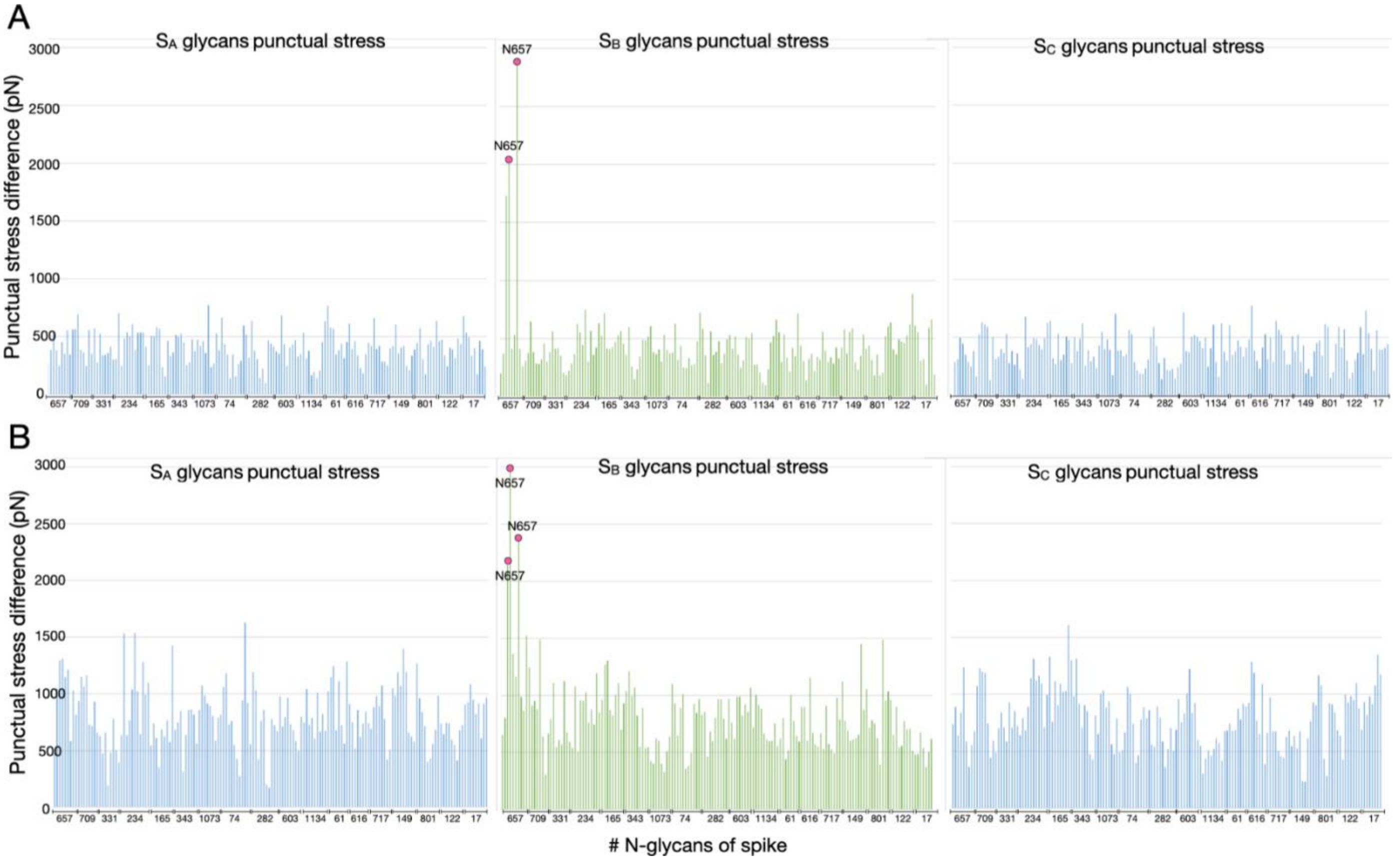


**Fig. S19 Absolute differences in punctual stress for N-glycans** on (A) open spike without ACE2 bound vs open spike with ACE2 bound (see Fig.S14), and on (B) the spike:ACE2 complex without GAGs bound vs the spike: ACE2 complex with GAGs bound (see Fig.S15). The absolute difference in punctual stress is the sum of the absolute values of scalar pairwise forces acting on each atom of the systems mentioned above and is computed by merging the last 400ns of each trajectory per condition. The residue numbers of the N-glycosylated asparagines are given on the x-axis for each spike subunit ($S_A$ in blue, $S_B$ in green, $S_C$ in light blue). Visual inspection of the trajectories reveals that the $S_B$ N657 glycan interacts with $S_A$ D843 upon ACE2 binding while upon ACE2 and GAG binding, the interaction is lost.

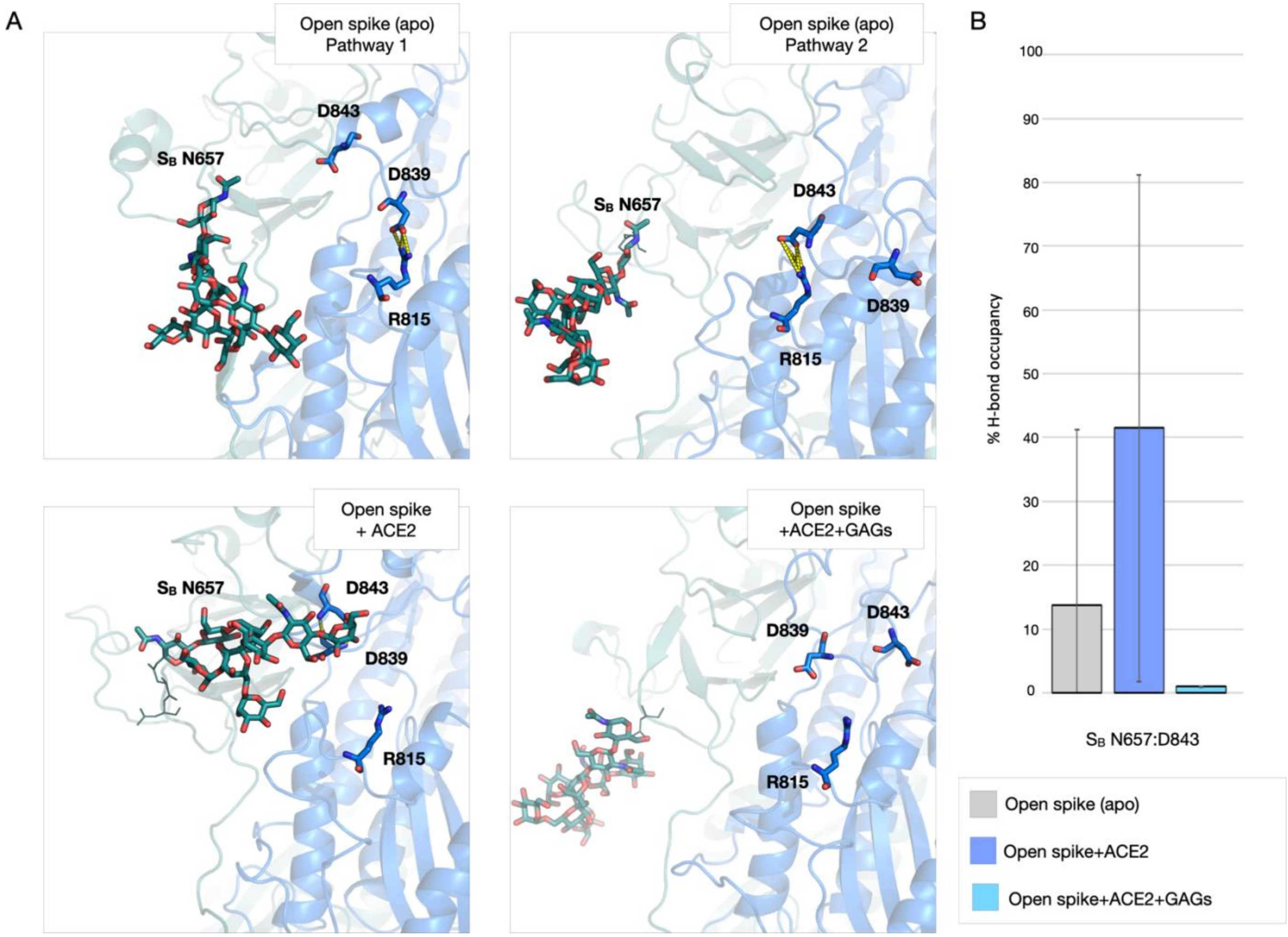


**Fig. S20 $S_B$ N657 glycan interactions with $S_A$ D843.** The punctual stress analysis for N-glycans (see Fig. S17) shows that $S_B$ N657 experiences a much higher stress difference than all other N-glycans. Visual inspection of the trajectories shows that the N657 glycan gains an interaction with D843 upon ACE2 binding in the absence of GAGs. (A) $S_A$ and $S_B$ are shown as cartoons colored in blue and teal, respectively. GAGs are not shown for clarity. Residues involved in the interactions are labeled and depicted as sticks colored by element (oxygen:red, nitrogen:blue, and sulfur:yellow). $S_B$ N657 glycan is shown as teal stick colored by element. Polar contacts are depicted as yellow dashed lines connecting interacting residues. (B) Trajectory averaged H-bond occupancy calculated for the last 400ns of MD simulations. For complexes with open spike only, all 4 replicas were considered but only one replica out of the 4 shows direct interactions between $S_B$ N657 glycan and $S_A$ D843 while upon ACE2 binding, the interaction is stable over all the replicas. Finally, upon ACE2 and GAG binding, these interactions are completely lost.

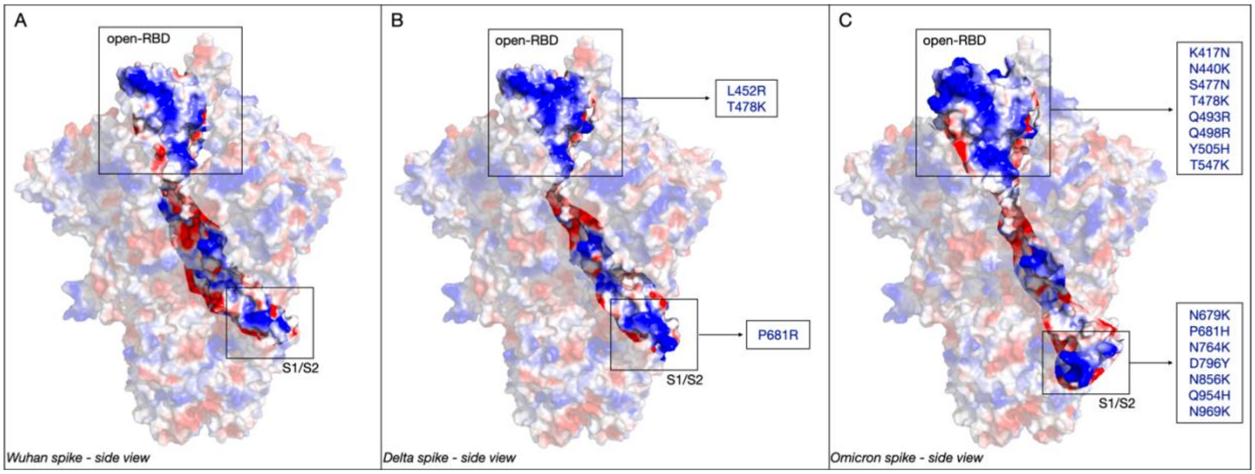


**Fig. S21 Spike variants evolve to optimize their binding to GAGs.** Representative structures of (A) Wuhan, (B) Delta, and (C) Omicron spike variants in an open conformation with 1 up-RBD are displayed as molecular surfaces with electrostatic potential mapped onto them to show the partially grooved positively charged path followed by the GAGs [17]. The Delta and Omicron spike models were prepared by starting from the Wuhan model and replacing the residues mutated based on sequencing studies (https://covariants.org/shared-mutations). The insets report the mutations in (B) Delta and (C) Omicron variants that increase the positive charge along the basic binding path which leads to enhanced binding to the negatively charged GAGs [18, 19]. Regions of positive and negative molecular electrostatic potentials are shown in blue and red, respectively.

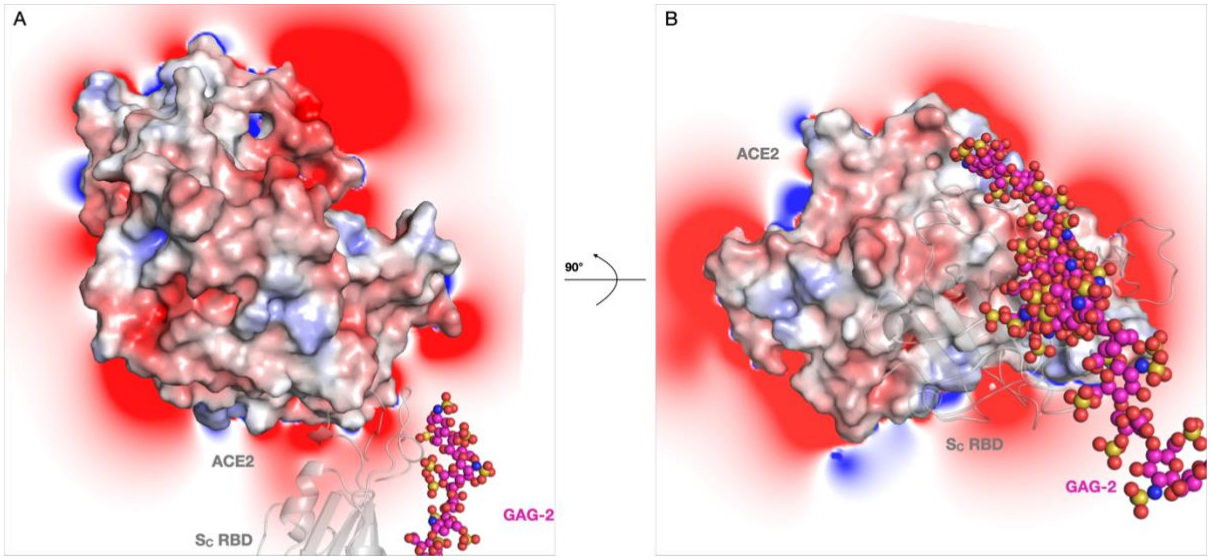


**Fig. S22 ACE2 lacks suitable basic domains for binding highly sulfated GAGs.** (A) Side and (B) front views of the spike:ACE2 interface in the initial spike:ACE2:GAGs complex. Spike $S_C$ up-RBD is shown in grey cartoon representation. The 31mer GAG-2 chain bound to the up-RBD is depicted as spheres colored by element with magenta carbons. ACE2 is displayed as a molecular surface colored by electrostatic potential with the electrostatic potential around it shown by a density map (positive: blue, negative: red). ACE2 has a net negative charge of -27e and is mostly surrounded by regions of negative electrostatic potential. ACE2 lacks a suitable positively charged domain to bind to the GAG chain, which has a net negative charge of -58e. Furthermore, the face of ACE2 interacting with spike RBD is negatively charged, and therefore unfavorable for binding a negatively charged GAG chain.

#### Supporting Tables

| PDBid 6M0J (HB 3.5Å) | | PDBid 6M17 (n.d) | | PDBid 6LZG (HB 3.5Å - VDW 4.5Å) | | PDBid 6VW1 (HB 3.35Å - VDW 3.9Å) | | PDBid 7DF4 (HB 3.5Å - VDW 4.0Å) | | MDs ACE2 (HB 3.0Å - VDW 3.5Å) | | MDs ACE2+GAGs (HB 3.0Å - VDW 3.5Å) | |
|---|---|---|---|---|---|---|---|---|---|---|---|---|---|
| ACE2 | RBD | ACE2 | RBD | ACE2 | RBD | ACE2 | RBD | ACE2 | RBD | ACE2 | RBD | ACE2 | RBD |
| / | / | / | / | S19 | A475, G476 | S19 | A475 | S19 | A475, G476 | S19 | A475 | S19 | A475, **G476** |
| Q24 | N487 | Q24 | Q474 | Q24 | A475, G476, N478 | Q24 | G476, N487 | Q24 | G476, N487 | Q24 | G476, N487 | Q24 | G476, N487 |
| / | / | / | / | T27 | F456, Y473, A475, Y489 | T27 | F456, A475, Y489 | T27 | F456, Y473, A475, Y489 | T27 | F456, A475, Y489 | T27 | F456, A475, Y489 |
| / | / | / | / | F28 | Y489 | F28 | Y489 | F28 | Y489 | F28 | Y489 | F28 | Y489 |
| D30 | K417 | D30 | K417 | D30 | K417, L455, F456 | / | / | D30 | F456 | / | / | / | |
| / | / | / | / | K31 | L455, F456, E484, Y489, F490, Q493 | K31 | F456, Q493 | K31 | L455, F456, Y489, F490, Q493 | K31 | L455, F456 | K31 | L455, F456 |
| / | / | H34 | Y453 | H34 | Y453, L455, Q493 | H34 | Y453, L455 | H34 | Y453, Q493, S494, Y495 | H34 | Y453, L455 | H34 | Y453, Q493, **S497** |
| E35 | Q493 | / | / | E35 | Q493 | E35 | Q493 | / | / | E35 | Q493 | E35 | Q493 |
| E37 | Y505 | / | / | E37 | Y505 | E37 | Y505 | / | / | E37 | Y505 | E37 | Y505 |
| D38 | Y449 | / | / | D38 | Y449, G496, Q498 | D38 | Y449 | / | / | D38 | Y449 | D38 | Y449 |
| Y41 | 500, N50 | Y41 | Q498, T500, N501 | Y41 | Q498, T500, N501 | Y41 | Q498, T500, N501 | Y41 | Q498, T500 | Y41 | Q498, T500, **N501** | Y41 | Q498, T500 |
| Q42 | 446, Y44 | Q42 | Q498 | Q42 | G446, Y449, Q498 | Q42 | Q498 | Q42 | G446 | / | / | / | / |
| / | / | / | / | L45 | Q498, T500 | L45 | Q498 | / | / | / | / | / | / |
| / | / | / | / | L79 | F486 | L79 | F486 | L79 | F486 | L79 | F486 | L79 | F486 |
| / | / | M82 | F486 | M82 | F486 | M82 | F486 | M82 | F486 | M82 | F486 | M82 | F486 |
| Y83 | 489, N48 | / | / | Y83 | F486, N487, Y489 | Y83 | F486, N487 | Y83 | F486, N487, Y489 | Y83 | F486, N487 | Y83 | F486, N487 |
| / | / | / | / | / | / | | | H84 | S494 | / | / | / | / |
| / | / | / | / | N330 | T500 | N330 | T500 | / | / | N330 | T500 | N330 | T500 |
| K353 | G502 | K353 | N501 | K353 | G496, N501, G502, Y505 | K353 | G496, N501, G502, Y505 | K353 | G496, N501, G502, Y505 | K353 | G496, N501, G502, Y505 | K353 | **Y495**, G496, N501, G502, Y505 |
| / | / | / | / | G354 | G502, Y505 | G354 | G502, Y505 | G354 | G502, Y505 | G354 | G502, Y505 | G354 | G502, Y505 |
| / | / | / | / | D355 | T500, G502 | D355 | T500 | D355 | T500 | D355 | T500 | D355 | T500, **G502** |
| / | / | R357 | T500 | R357 | T500 | R357 | T500 | R357 | T500 | R357 | T500 | R357 | T500 |
| / | / | / | / | / | / | / | / | A386 | Y505, R393 | / | / | / | / |
| R393 | Y505 | / | / | R393 | Y505 | / | / | R393 | Y505 | / | / | / | / |

**Tab. S1 Residue-residue contacts at the spike RBD:ACE2 interface.** Comparison of residue contacts at the interface between spike RBD and ACE2 retrieved from experimentally determined PDB structures (PDBid: 6M0J[21], 6M17[22], 6LZG[23], 6VW1[24], 7DF4[25]), and for simulated systems of open spike with ACE2 bound and open spike with ACE2 and GAGs bound. GAG binding results in the gain of interactions of residues highlighted in bold red.

| Spike glycomic profile | |
|---|---|
| Aminoacid residue | Glycan type |
| N17 | 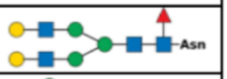 |
| N61 | 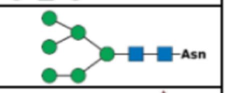 |
| N74 | 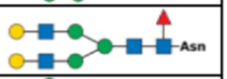 |
| N122 | 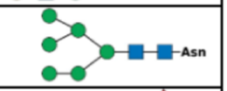 |
| N149 | 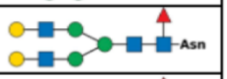 |
| N165 | 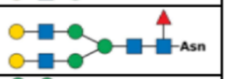 |
| N234 | 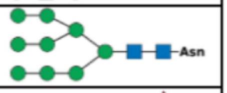 |
| N282 | 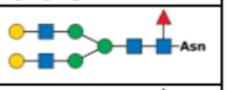 |
| N331 | 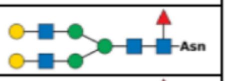 |
| N343 | 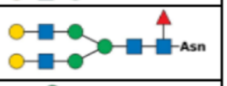 |
| N603 | 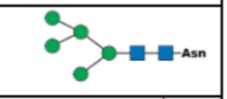 |
| N616 | 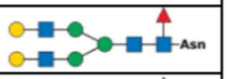 |
| N657 | 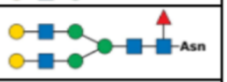 |
| N709 | 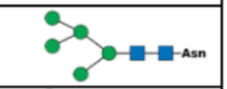 |
| N717 | 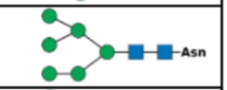 |
| N801 | 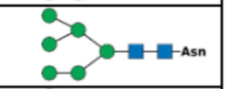 |
| N1074 | 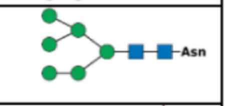 |
| N1134 | 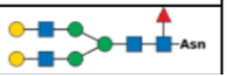 |

| ACE2 glycomic profile | |
|---|---|
| Aminoacid residue | Glycan type |
| N53 | 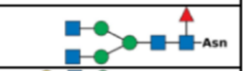 |
| N90 | 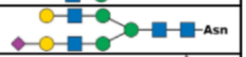 |
| N103 | 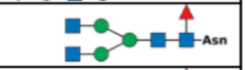 |
| N322 | 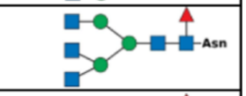 |
| N432 | 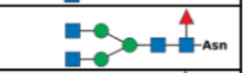 |
| N546 | 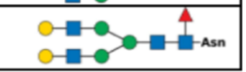 |

| Monosaccharides | | | | |
|---|---|---|---|---|
| GlcNAc | Fuc | Man | Gal | Neu5Ac |
| 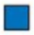 | 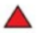 | 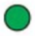 | 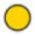 | 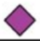 |

**Tab.S2 Glycosylation pattern of the simulated spike and ACE2 glycoproteins.** N-glycan types and the respective Asn to which they are covalently bound on spike and ACE2 are from refs [26,27]. GlcNAc = N-Acetylglucosamine; Fuc = Fucose; Man = Mannose; Gal = Galactose; Neu5Ac = Sialic Acid.

| Simulation method | Simulated system | Simulated system components | | | Starting structure | Random Force applied | # Atoms | Length ns | # Replica |
|---|---|---|---|---|---|---|---|---|---|
| | | Spike | ACE2 | GAG chains | | | | | |
| Conventional MD carried out by Paiardi et al. [17] | Spike | + | - | - | Modelled | - | 716254 | 1000 | 4 |
| | Spike:GAGs | + | - | 3x 31mer | Modelled | - | 715862 | 1000 | 4 |
| Conventional MD | Spike:ACE2 | + | + | - | Modelled | - | 873223 | 1000 | 4 |
| | Spike:ACE2:GAGs | + | + | 3x 31mer | Modelled | - | 872404 | 1000 | 6 |
| RAMD | Spike:ACE2 | + | + | - | Last snapshot of conventional MD replica | + | 900170 | ≤ 5 | 4x15 |
| | Spike:ACE2:GAGs | + | + | 3x 31mer | Last snapshot of conventional MD replica | + | 1061017 | ≤ 5 | 6x15 |

**Tab. S3** Summary of the MD trajectories carried out by Paiardi et al.[17] and the MD simulations performed in this study. For details, see the Supporting Material and Methods section.

#### 3. Supporting Movies

**Movie S1 – N-glycans and GAGs concur in strengthening the spike:ACE2 interaction.** The movie shows first the spike:ACE2:GAGs structure (replica 1) with a histogram showing the difference in the interactions in the absence and presence of GAGs of the buffering N-glycans between ACE2 and spike-NTD and between ACE2 N53 glycan and GAG-2 followed by a focus on the interactions established by ACE2 N90 glycan with spike residues (corresponding to Figure 4A in the main text and Figure S10 in the SI Appendix).

**Movie S2 – N-glycans and GAGs concur in extending the residence time of the complex.** The movie first shows the spike:ACE2 and spike:ACE2:GAGs structures extracted from the last snapshots of conventional MD simulations (replica1) and used as starting structures for performing RAMD simulations followed by examples of egress pathways in the absence (left) and presence (right) of GAG chains (corresponding to Figure S12 in the SI Appendix and Figure 4B in the main text, respectively).

#### 4. Supporting References


(1) *AMBER 2020*; University of California, San Francisco.: 2020. (accessed.

(2) Maier, J. A.; Martinez, C.; Kasavajhala, K.; Wickstrom, L.; Hauser, K. E.; Simmerling, C. ff14SB: Improving the Accuracy of Protein Side Chain and Backbone Parameters from ff99SB. *Journal of Chemical Theory and Computation* **2015**, *11* (8), 3696-3713. DOI: 10.1021/acs.jctc.5b00255 (acccessed 2023-08-14T13:44:16).

(3) Kirschner, K. N.; Yongye, A. B.; Tschampel, S. M.; González-Outeiriño, J.; Daniels, C. R.; Foley, B. L.; Woods, R. J. GLYCAM06: A generalizable biomolecular force field. Carbohydrates. *Journal of Computational Chemistry* **2008**, *29* (4), 622-655. DOI: 10.1002/jcc.20820 (acccessed 2023-08-14T13:46:01).

(4) Wang, J.; Wolf, R. M.; Caldwell, J. W.; Kollman, P. A.; Case, D. A. Development and testing of a general amber force field. *J Comput Chem* **2004**, *25* (9), 1157-1174. DOI: 10.1002/jcc.20035.

(5) Pekka, M.; Lennart, N. Structure and Dynamics of the TIP3P, SPC, and SPC/E Water Models at 298 K. *The Journal of Physical Chemistry A* **2001**, *105* (43), 9954-9960. DOI: 10.1021/jp003020w.

(6) Kräutler , V.; van Gunsteren , W. F.; Hünenberger , P. H. A fast SHAKE algorithm to solve distance constraint equations for small molecules in molecular dynamics simulations. Journal of Computational Chemistry ed.; Journal of Computational Chemistry: Journal of Computational Chemistry, 2001; Vol. 22, pp 501-508.

(7) Essmann, U.; Perera, L.; Berkowitz, M. L.; Darden; Tom; Lee, H.; Pedersen; G., L. A smooth particle mesh Ewald method The Journal of Chemical Physics: 1995; Vol. 103, pp 8577–8593.

(8) Kokh, D. B.; Amaral, M.; Bomke, J.; Grädler, U.; Musil, D.; Buchstaller, H.-P.; Dreyer, M. K.; Frech, M.; Lowinski, M.; Vallee, F.; et al. Estimation of Drug-Target Residence Times by τ-Random Acceleration Molecular Dynamics Simulations. *Journal of Chemical Theory and Computation* **2018**, *14* (7), 3859-3869. DOI: 10.1021/acs.jctc.8b00230 (acccessed 2023-08-14T13:23:19).

(9) Kokh, D. B.; Doser, B.; Richter, S.; Ormersbach, F.; Cheng, X.; Wade, R. C. A workflow for exploring ligand dissociation from a macromolecule: Efficient random acceleration molecular dynamics simulation and interaction fingerprint analysis of ligand trajectories. *The Journal of Chemical Physics* **2020**, *153* (12), 125102. DOI: 10.1063/5.0019088 (acccessed 2023-08-14T14:01:36).

(10) Sousa Da Silva, A. W.; Vranken, W. F. ACPYPE - AnteChamber PYthon Parser interfacE. *BMC Research Notes* **2012**, *5* (1), 367. DOI: 10.1186/1756-0500-5-367 (acccessed 2023-08-14T14:01:51).

(11) Van Der Spoel, D.; Lindahl, E.; Hess, B.; Groenhof, G.; Mark, A. E.; Berendsen, H. J. GROMACS: fast, flexible, and free. *J Comput Chem* **2005**, *26* (16), 1701-1718. DOI: 10.1002/jcc.20291 From NLM.

(12) Parrinello, M.; Rahman; A. Polymorphic transitions in single crystals: A new molecular dynamics method. *Journal of Applied Physics*: 1981; Vol. 52, pp 7182–7190.

(13) Evans, D. J.; Holian, B. L. The Nose–Hoover thermostat. 1985; Vol. 83, pp 4069–4074.

(14) Humphrey, W.; Dalke, A.; Schulten, K. VMD: visual molecular dynamics. *J Mol Graph* **1996**, *14* (1), 33-38, 27-38. DOI: 10.1016/0263-7855(96)00018-5.

(15) Roe, D. R.; Cheatham, T. E. PTRAJ and CPPTRAJ: Software for Processing and Analysis of Molecular Dynamics Trajectory Data. *J Chem Theory Comput* **2013**, *9* (7), 3084-3095. DOI: 10.1021/ct400341p.

(16) Costescu, B. I.; Gräter, F. Time-resolved force distribution analysis. *BMC Biophysics* **2013**, *6* (1), 5. DOI: 10.1186/2046-1682-6-5 (acccessed 2023-08-14T14:10:09).

(17) Paiardi, G.; Richter, S.; Oreste, P.; Urbinati, C.; Rusnati, M.; Wade, R. C. The binding of heparin to spike glycoprotein inhibits SARS-CoV-2 infection by three mechanisms. *Journal of Biological Chemistry* **2022**, *298* (2), 101507. DOI: 10.1016/j.jbc.2021.101507 (acccessed 2023-08-14T13:05:43).

(18) Kim, S. H.; Kearns, F. L.; Rosenfeld, M. A.; Casalino, L.; Papanikolas, M. J.; Simmerling, C.; Amaro, R. E.; Freeman, R. GlycoGrip: Cell Surface-Inspired Universal Sensor for Betacoronaviruses. *ACS Central Science* **2022**, *8* (1), 22-42. DOI: 10.1021/acscentsci.1c01080 (acccessed 2023-08-14T14:49:08).

(19) Kim, S. H.; Kearns, F. L.; Rosenfeld, M. A.; Votapka, L.; Casalino, L.; Papanikolas, M.; Amaro, R. E.; Freeman, R. SARS-CoV-2 evolved variants optimize binding to cellular glycocalyx. *Cell Rep Phys Sci* **2023**, *4* (4), 101346. DOI: 10.1016/j.xcrp.2023.101346.

(20) Clausen, T. M.; Sandoval, D. R.; Spliid, C. B.; Pihl, J.; Perrett, H. R.; Painter, C. D.; Narayanan, A.; Majowicz, S. A.; Kwong, E. M.; Mcvicar, R. N.; et al. SARS-CoV-2 Infection Depends on Cellular Heparan Sulfate and ACE2. *Cell* **2020**, *183* (4), 1043-1057.e1015. DOI: 10.1016/j.cell.2020.09.033 (acccessed 2023-08-14T12:53:53).

(21) Lan, J.; Ge, J.; Yu, J.; Shan, S.; Zhou, H.; Fan, S.; Zhang, Q.; Shi, X.; Wang, Q.; Zhang, L.; et al. Structure of the SARS-CoV-2 spike receptor-binding domain bound to the ACE2 receptor. *Nature* **2020**, *581* (7807), 215-220. DOI: 10.1038/s41586-020-2180-5 (acccessed 2023-08-14T09:17:56).
(22) Yan, R.; Zhang, Y.; Li, Y.; Xia, L.; Guo, Y.; Zhou, Q. Structural basis for the recognition of SARS-CoV-2 by full-length human ACE2. *Science* **2020**, *367* (6485), 1444-1448. DOI: 10.1126/science.abb2762.
(23) Wang, Q.; Zhang, Y.; Wu, L.; Niu, S.; Song, C.; Zhang, Z.; Lu, G.; Qiao, C.; Hu, Y.; Yuen, K.-Y.; et al. Structural and Functional Basis of SARS-CoV-2 Entry by Using Human ACE2. *Cell* **2020**, *181* (4), 894-904.e899. DOI: 10.1016/j.cell.2020.03.045 (acccessed 2023-08-14T09:16:39).
(24) Shang, J.; Ye, G.; Shi, K.; Wan, Y.; Luo, C.; Aihara, H.; Geng, Q.; Auerbach, A.; Li, F. Structural basis of receptor recognition by SARS-CoV-2. *Nature* **2020**, *581* (7807), 221-224. DOI: 10.1038/s41586-020-2179-y (acccessed 2023-08-14T13:12:54).
(25) Xu, C.; Wang, Y.; Liu, C.; Zhang, C.; Han, W.; Hong, X.; Hong, Q.; Wang, S.; Zhao, Q.; Yang, Y.; et al. Conformational dynamics of SARS-CoV-2 trimeric spike glycoprotein in complex with receptor ACE2 revealed by cryo-EM. *Sci Adv* **2021**, *7* (1). DOI: 10.1126/sciadv.abe5575.
(26) Watanabe, Y.; Allen, J. D.; Wrapp, D.; McLellan, J. S.; Crispin, M. Site-specific glycan analysis of the SARS-CoV-2 spike. *Science* **2020**, *369* (6501), 330-333. DOI: 10.1126/science.abb9983.
(27) Zhao, P.; Praissman, J. L.; Grant, O. C.; Cai, Y.; Xiao, T.; Rosenbalm, K. E.; Aoki, K.; Kellman, B. P.; Bridger, R.; Barouch, D. H.; et al. Virus-Receptor Interactions of Glycosylated SARS-CoV-2 Spike and Human ACE2 Receptor. *Cell Host & Microbe* **2020**, *28* (4), 586-601.e586. DOI: 10.1016/j.chom.2020.08.004 (acccessed 2023-08-14T12:39:14).
